# Atlas of Neuromuscular Organization in the ctenophore *Pleurobrachia bachei*

**DOI:** 10.1101/385435

**Authors:** Tigran P. Norekian, Leonid L. Moroz

## Abstract

Enigmatic ctenophores are descendants of one of the earliest branching metazoan lineage. Their nervous systems are equally elusive. The lack of convenient neurogenic molecules and neurotransmitters suggests an extensive parallel evolution and independent origins of neurons and synapses. However, the field is logged behind due to the lack of microanatomical data about the neuro-muscular systems in this group of animals. Here, using immunohistochemistry and scanning electron microscopy, we describe the organization of both muscular and nervous systems in the sea gooseberry, *Pleurobrachia bachei*, from North Pacific. The diffused neural system of *Pleurobrachia* consists of two subsystems: the subepithelial neural network and the mesogleal net with about 5000-7000 neurons combined. Our data revealed the unprecedented complexity of neuromuscular organization in this basal metazoan lineage. The anatomical diversity of cell types includes at least nine broad categories of neurons, five families of surface receptors and more than two dozen types of muscle cells as well as regional concentrations of neuronal elements to support ctenophore feeding, complex swimming, escape and prey capture behaviors. In summary, we recognize more than 80 total morphological cell types. Thus, in terms of cell type specification and diversity, ctenophores significantly exceed what we currently know about other prebilaterian groups (placozoan, sponges, and cnidarians), and some basal bilaterians.

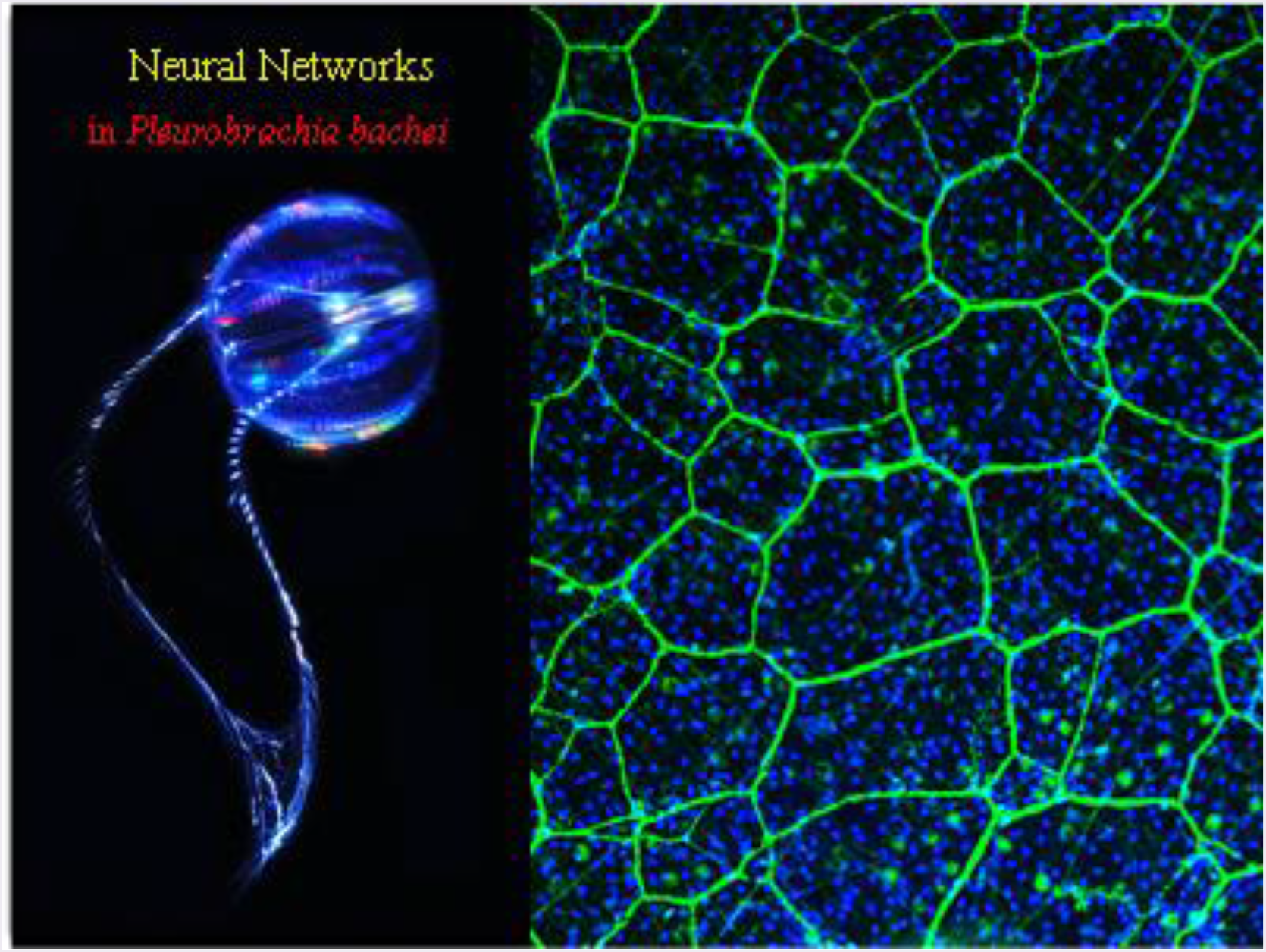

## 1 INTRODUCTION

Ctenophores are enigmatic basal metazoans (Kozloff, 1990; Brusca and Brusca, 2003; Nielsen, 2012), and their nervous systems are equally elusive even in term of their anatomy and functions (Tamm, 1982; Hernandez-Nicaise, 1991; Moroz, 2015). Due to the fragile nature of these jelly-like marine organisms, it is difficult to work with comb jellies outside of their marine habitats. The second challenge is a unique organization (Hernandez-Nicaise, 1991) and remarkably different molecular makeup of ctenophore nervous systems (Moroz, 2015), which significantly limit the use of conventional protocols to label neurons.

These challenges lead to numerous controversies about the early evolution of animals in general, and the origins of neurons and muscles in particular (Moroz, 2014). The exact phylogenetic position of Ctenophora has been extensively debated over the last decade (Nielsen, 2012; Ryan et al., 2013; Moroz et al., 2014; Borowiec et al., 2015; Dunn et al., 2015; Whelan et al., 2015a; Whelan et al., 2015b; Halanych et al., 2016; Telford et al., 2016; Alamaru et al., 2017; Arcila et al., 2017; Cavalier-Smith, 2017; Feuda et al., 2017; King and Rokas, 2017; Shen et al., 2017; Simion et al., 2017; Whelan et al., 2017). However, the most recent models of evolution and interdisciplinary data strongly suggest that Ctenophora is a sister group to all extant Metazoa including nerveless sponges, which are morphologically simpler than comb jellies (Moroz et al., 2014; Arcila et al., 2017; Shen et al., 2017; Whelan et al., 2017).

The combination of genomic, physiological, neurochemical, pharmacological, proteomic and metabolomic data indicate that neural systems and synapses in ctenophores evolved independently from the rest of animals (Moroz et al., 2014; Moroz, 2015; Moroz and Kohn, 2015; 2016). The muscles and mesoderm might also evolve independently in ctenophores and cnidarians/bilaterians (Steinmetz et al., 2012; Moroz et al., 2014). Molecular analyses further confirmed extensive parallel evolution and ctenophore-lineage-specific diversification of ion channels, gap junction proteins, transmitter receptors, neuropeptides, RNA editing, RNA modifications and signaling pathways (Moroz and Kohn, 2016).

But the field is logged behind due to the lack of microanatomical data about the structure of neuro-muscular systems in ctenophores. Here, using immunohistochemistry and scanning electron microscopy, we expand earlier developmental (Norekian and Moroz, 2016) and comparative studies (Jager et al., 2011), and characterize the organization of both muscular and nervous systems in the sea gooseberry, *Pleurobrachia bachei*, from North Pacific. Our data revealed the unprecedented complexity of neuro-muscular organization in this basal metazoan lineage with diverse types of neurons and muscles as well as regional concentrations of neuronal elements to support ctenophore feeding, complex swimming, escape and prey capture behaviors.

## 2 MATERIALS AND METHODS

### 2.1 Animals

Adult specimens of *Pleurobrachia bachei* were collected from the breakwater and held in 1-gallon glass jars in the large tanks with constantly circulating seawater at 10° C. Experiments were carried out at Friday Harbor Laboratories, the University of Washington in the spring-summer seasons of 2012-2017.

### 2.2 Scanning Electron Microscopy (SEM)

The details of the protocol have been described elsewhere (Norekian and Moroz, 2016). Briefly, adult animals were fixed in 2.5% glutaraldehyde in 0.2 M phosphate-buffered saline (pH=7.6) for 4 hours at room temperature and washed for 2 hours in 2.5% sodium bicarbonate. Large animals, more than 1 cm in diameter, were then dissected into a few smaller pieces, while small animals were processed as whole mounts. For secondary fixation, we used 2% osmium tetroxide in 1.25% Sodium Bicarbonate for 3 hours at room temperature. Tissues were then rinsed several times with distilled water, dehydrated in ethanol and placed in Samdri-790 (Tousimis Research Corporation) for Critical point drying. After the drying process, the tissues were placed on the holding platforms and processed for metal coating on Sputter Coater (SPI Sputter). SEM analyses and photographs were done on NeoScope JCM-5000 microscope (JEOL Ltd., Tokyo, Japan).

### 2.3 Immunocytochemistry and phalloidin staining

Adult *Pleurobrachia* were fixed overnight in 4% paraformaldehyde in 0.1 M phosphate-buffered saline (PBS) at +4-5° C and washed for 2-6 hours in PBS. The fixed animals were then dissected to get better access to specific organs and processed as described for embryonic and larval *Pleurobrachia* (Norekian and Moroz, 2016), with minor modifications. To label the muscle fibers, we used well-known marker phalloidin (Alexa Fluor 488 and Alexa Fluor 594 phalloidins from Molecular Probes), which binds to F-actin (Wulf et al., 1979) The tissue was incubated in phalloidin solution in PBS for 4 to 8 hours at a final dilution 1:80 and then washed in several PBS rinses for 6 hours.

For immunocytochemistry, adult animals were fixed the same way in 4% paraformaldehyde in PBS for 8-12 hrs and washed in PBS several times for 6-12 hrs. Sometimes, we used smaller, 2-6 mm, pieces for better access to different areas and better antibody penetration. Tissues were pre-incubated overnight in a blocking solution of 6% goat serum in PBS and then incubated for 48 hours at +4-5° C in the primary antibodies diluted in 6% of the goat serum at a final dilution 1:40. The rat monoclonal anti-tubulin antibody (AbD Serotec Cat# MCA77G, RRID: AB_325003) recognizes the alpha subunit of tubulin, specifically binding tyrosylated tubulin (Wehland and Willingham, 1983; Wehland et al., 1983). Following a series of PBS washes for total 6 hours, the tissues were incubated for 12 hours in secondary antibodies of one of two types of goat anti-rat IgG antibodies: Alexa Fluor 488 conjugated (Molecular Probes, Invitrogen, Cat# A11006, RRID: AB_141373) and goat anti-rat IgG antibody Alexa Fluor 594 conjugated (Molecular Probes, Invitrogen, Cat# A11007, RRID: AB_141374), at a final dilution 1:20. To stain nuclei, the tissues were incubated for 8 hours in SYTOX Green nucleic acid stain from Life Technologies (Cat# S7020) or was mounted in VECTASHIELD Hard-Set Mounting Medium with DAPI (Cat# H-1500).

After washing in PBS, the preparations were mounted in Fluorescent Mounting Media(KPL) on glass microscope slides to be viewed and photographed using a Nikon Research Microscope Eclipse E800 with Epi-fluorescence using standard TRITC and FITC filters, BioRad (Radiance 2000 system) Laser Scanning confocal microscope or Nikon C1 Laser Scanning confocal microscope. Multiple optic sections were taken through the depth of the samples for 3D reconstructions. To test for the specificity of immunostaining either the primary or the secondary antibody were omitted from the procedure. In both cases, no labeling was detected.

### 2.4 Antibody specificity

Rat monoclonal anti-tyrosinated alpha-tubulin antibody is raised against yeast tubulin, clone YL1/2, isotype IgG2a (Serotec Cat # MCA77G; RRID: AB_325003). The antibody is routinely tested in ELISA on tubulin, and ctenophore *Pleurobrachia* is listed in species reactivity (manufacturer’s technical information). The epitope recognized by this antibody appears to be a linear sequence requiring an aromatic residue at the C terminus, with the two adjacent amino acids being negatively charged, represented by Glu-Glu-Tyr in Tyr-Tubulin (manufacturer’s information). As reported by Wehland et al. (Wehland and Willingham, 1983; Wehland et al., 1983)-this rat monoclonal antibody “reacts specifically with the tyrosylated form of brain alpha-tubulin from different species.” They showed that “YL 1/2 reacts with the synthetic peptide Gly-(Glu)3-Gly-(Glu)2-Tyr, corresponding to the carboxyterminal amino acid sequence of tyrosylated alpha-tubulin, but does not react with Gly-(Glu)3-Gly-(Glu)2, the constituent peptide of detyrosinated alpha-tubulin”. The epitope recognized by this antibody has been extensively studied including details about antibody specificity and relevant assays (Wehland and Willingham, 1983; Wehland et al., 1983). Equally important, this specific monoclonal antibody has been used before in two different species of both adult and larval *Pleurobrachia*: the previously obtained staining patterns are very similar to our experiments (Jager et al., 2011; Moroz et al., 2014; Norekian and Moroz, 2016).

## 3 RESULTS

### 3.1 Brief Introduction to the *Pleurobrachia* anatomy

The pelagic *Pleurobrachia bachei* has the biradial symmetrical body-plan (Kozloff, 1990; Hernandez-Nicaise, 1991), characteristic for the phylum including the morphological features of the canonical cydippid larvae (Fig. 1A). Two symmetrical polar fields are connected to the aboral organ forming the aboral pole of the animal, which is opposite to the mouth or the oral pole. Here, we use the term of **the aboral organ** to distinguish it from the non-homologous apical organ in bilaterian larvae (Nielsen, 2012).

**Figure 1.**
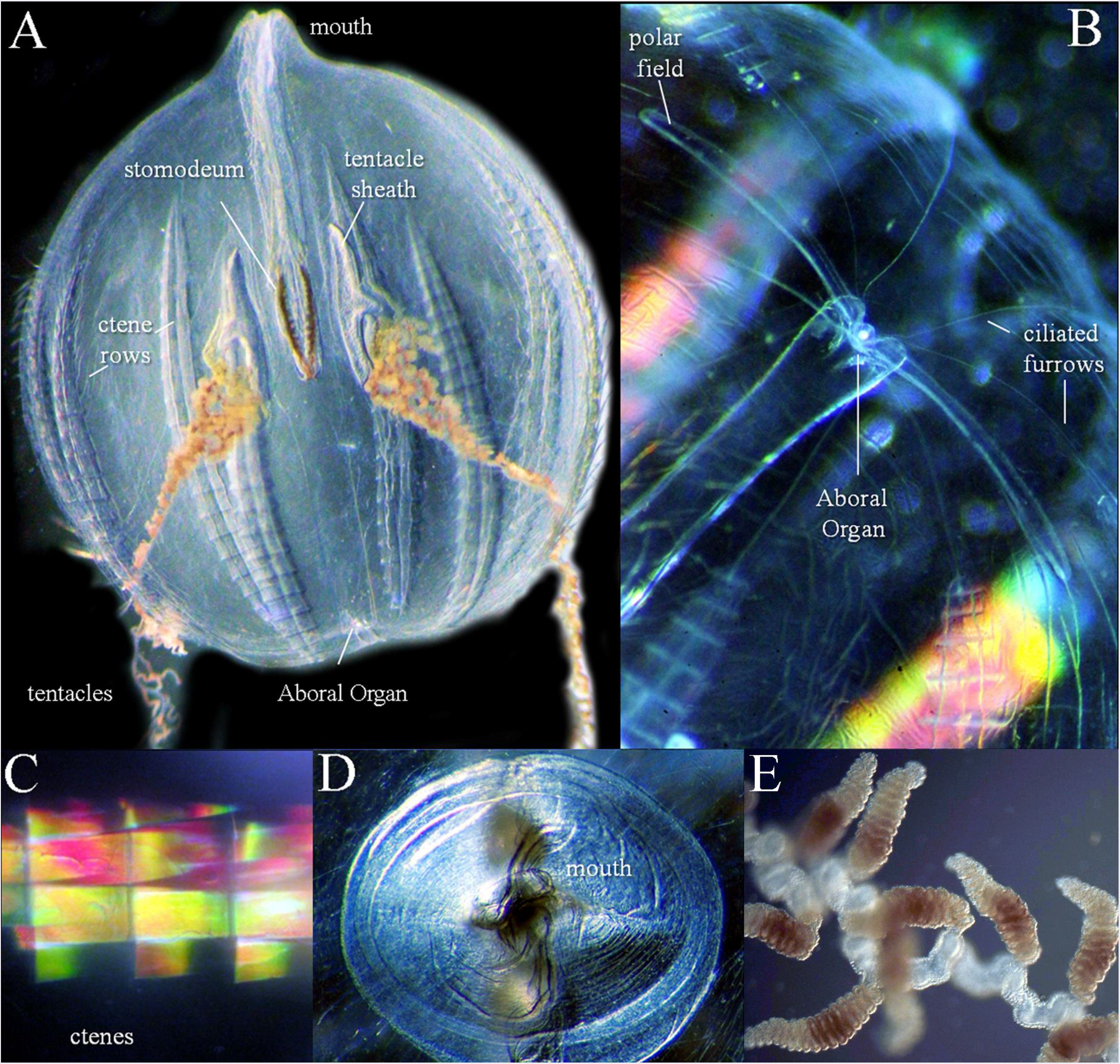
The ctenophore *Pleurobrachia bachei* and its anatomy. **A**: General view and major organs visible in an alive animal (1 cm). See text and other figures for microscopic details of every structure. Ctenes rows are also referred to combs plates and consist of many fused cilia. The tentacle shield often called the tentacle pocket. **B**: The aboral pole. The bright iridescence of swimming comb cilia is noticeable even through the transparent mesoglea and separately in **C. D**: The oral view of the mouth. **E**: A part of a tentacle with tentilla (small tentacles), which sometimes are pigmented.

Together with the mouth (Fig. 1D), the polar fields are making the first symmetry plane of the body, perpendicular to the tentacle plane. The size of polar fields is age-dependent, from less than 1 mm in juveniles to very long and narrow loops in adult animals. A bright single statolith (Tamm, 2015) is visible in the center of the aboral organ (Fig. 1B). The aboral organ performs multiple integrative functions and has a complex ultrastructural organization (Aronova, 1974; Tamm, 1982; Hernandez-Nicaise, 1991; Lowe, 1997; Tamm, 2016b). Eight thin and long ciliated furrows (Fig. 1B) connect the aboral organ with eight swim cilia rows (Fig. 1A, C), which represent the main means of locomotion in the ctenophores. A pair of tentacles (Fig. 1A, E), with sticky colloblasts, is used to collect the food and bring it to the mouth. Upon complete retraction, tentacles withdraw into a pair of tentacle pockets, which extend to the center of the body (Fig. 1A). The mouth (Fig. 1D) opens into the muscular pharynx and then short stomach chamber, which then connects to a system of gastrovascular canals.

### 3.2 The Aboral Organ and Polar Fields are complex integrative structures with multiple neural elements

#### *Scanning electron microscopy* (SEM)

Since ctenophores are composed of >95% of water, SEM imaging is very challenging because the dehydration and fixation cause significant tissue deformation. Nevertheless, we successfully adopted the SEM protocol for both juvenile and adult *Pleurobrachia bachei* (Norekian and Moroz, 2016) to reveal the microscopic anatomy for most of the tissues including the aboral organ and polar fields (Fig. 2). The SEM sagittal section shows a dome structure over the aboral organ, which consists of long cilia protecting the statolith (Fig. 2A, B).

**Figure 2.**
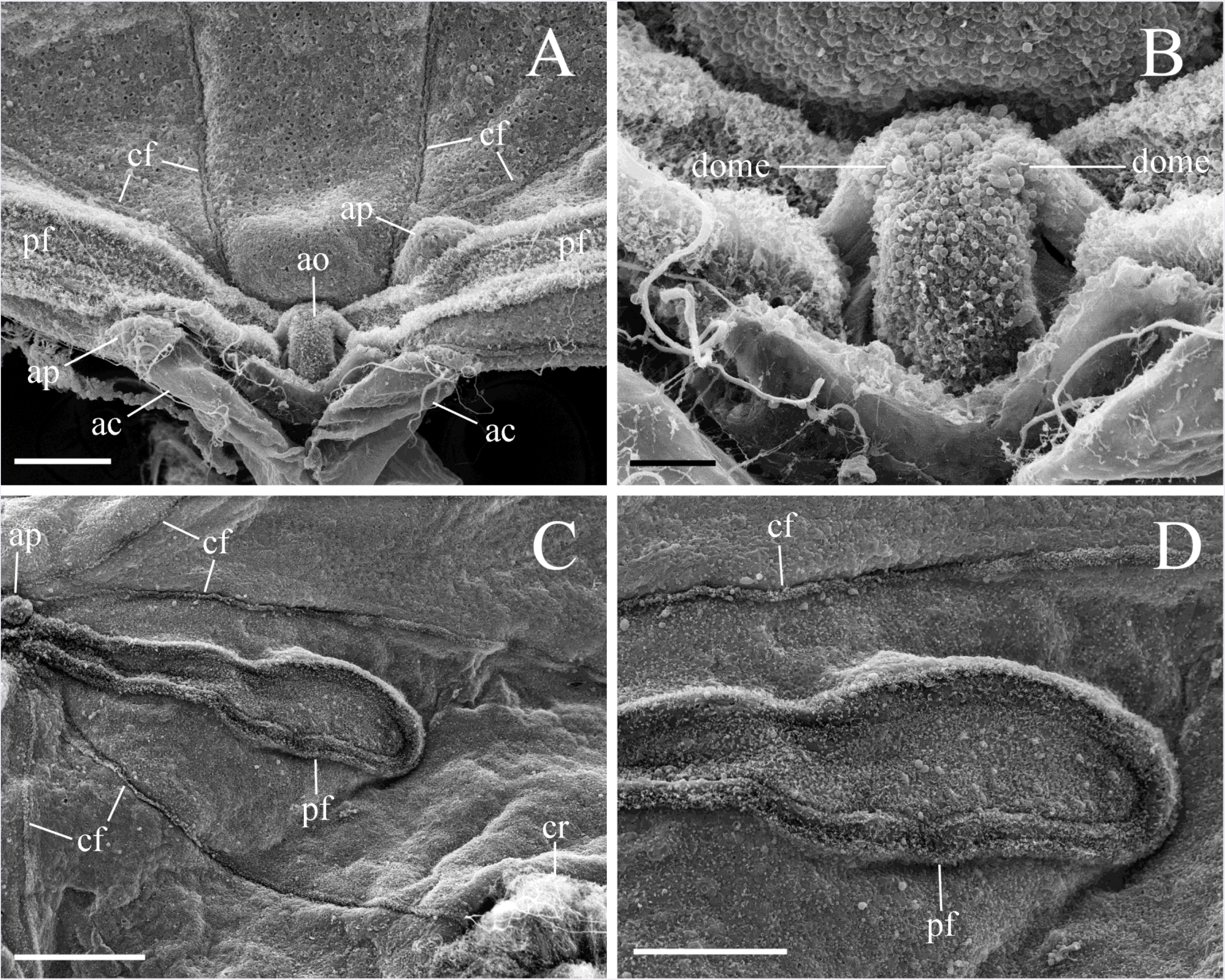
SEM images of the aboral organ and polar fields. **A**: Two polar fields and several ciliated furrows converge to the aboral organ. Two short anal canals terminate in two anal pores located diagonally on the opposite sides of the aboral organ. **B**: The aboral organ is covered by a protective dome. **C, D**: The polar field with a band of cilia outlining its edge and neighboring ciliated furrows. Abbreviations: *ao* - aboral organ; *pf* - polar field; *ac* - anal canal; *ap* - anal pore; *cf* - ciliated furrow; *cr* - comb row. Scale bars: A - 100 μm, B - 20 μm, C - 200 μm, D - 100 μm.

The aboral end of the digestive tract has two short branches on both sides of the aboral organ (Fig. 2A). Each of these branches opens as an anal pore (a small bulge on the SEM image in the Fig. 2A, C). In fact, the anal pores act as the functional anus similarly to the majority of bilaterian animals (Presnell et al., 2016; Tamm, 2016a).

The horizontal SEM view shows that the polar fields are formed by the short ciliated bands on their outside edges (Fig. 2C, D). The densely ciliated furrows connect the aboral organ with the comb rows and might represent conductive paths (Tamm, 1982; Tamm, 2014).

#### Immunoreactivity for neurons and cilia

As was shown in previous studies (Jager et al., 2011; Moroz et al., 2014; Norekian and Moroz, 2016), the tubulin antibodies are useful tools for the identification of neurons in ctenophores. In adult *Pleurobrachia*, the tubulin antibody staining always produced a very low background and a good signal/noise ratio in most parts of the body. Specific labeling occurred in neural elements and different cilia associated with such structures as the aboral organ, two polar fields, eight ciliated furrows and two tentacular nerves approaching the aboral organ (Fig. 3A). The tubulin immunoreactivity (IR) also revealed a diffused neural network covering the entire surface of the body.

**Figure 3.**
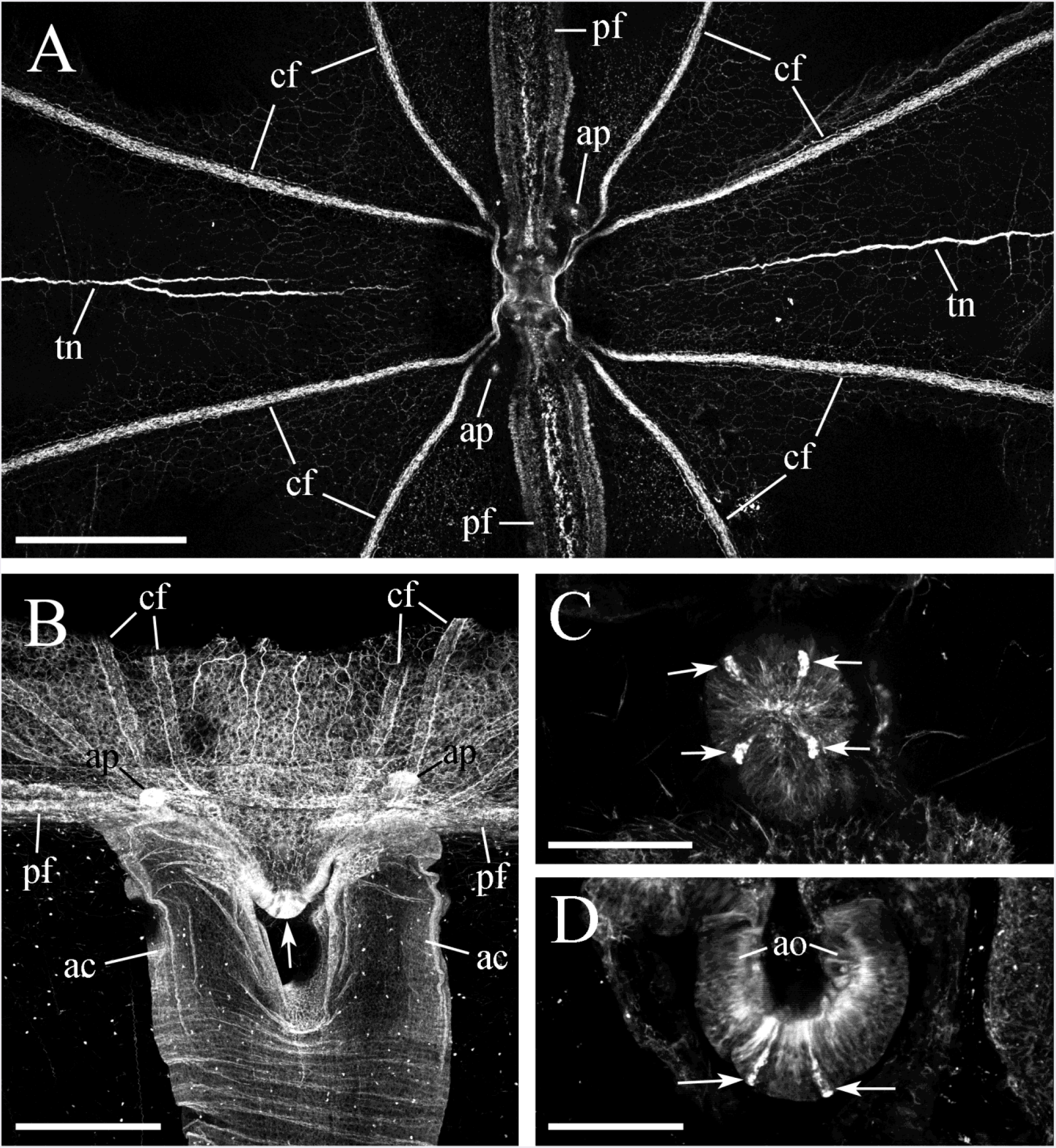
Tubulin IR in the aboral complex. **A**: Horizontal view of the aboral pole shows two polar fields, eight ciliated furrows, and two tentacular nerves. **B**: Sagittal view of the aboral pole shows two short anal canals under the aboral organ (arrow) with the anal pores located on opposite sides. **C, D**: Balancers (arrows) at the horizontal (C) and sagittal sections (D) of the aboral organ. Abbreviations: *ao* - aboral organ; *pf* - polar field; *cf* - ciliated furrow; *ac* - anal canal; *ap* - anal pore; *tn* - tentacular nerve. Scale bars: A - 500 μm; B - 300 μm; C - 100 μm; D - 60 μm.

In the sagittal view, the aboral organ resembles a horseshoe. The aboral canal of the gastrovascular system branches into two short anal canals and subsequently two openings near the aboral organ (Fig. 3A, B, and 6A). These anal pores can be seen in both sagittal and horizontal planes of the images (Fig. 3A, B, and 6A). Inside the aboral organ, tubulin IR staining revealed four elongated cell-like structures (Fig. 3C, D), which were located under four balances that support the statolith (Tamm, 1982; Tamm, 2015; 2016b). The statolith itself was often washed away during the incubation and washing in numerous solutions.

The bright tubulin-specific fluorescence of aboral organ observed in all preparations (e.g., Fig. 4A, C) was related to the large number of very tightly packed immunoreactive small cells covering the entire area (Fig. 4C, D). The size of these cells varied between 1 and 4 μm in diameter. Some of them revealed elongated processes suggesting their neuron-like nature (Table 1).

**Figure 4.**
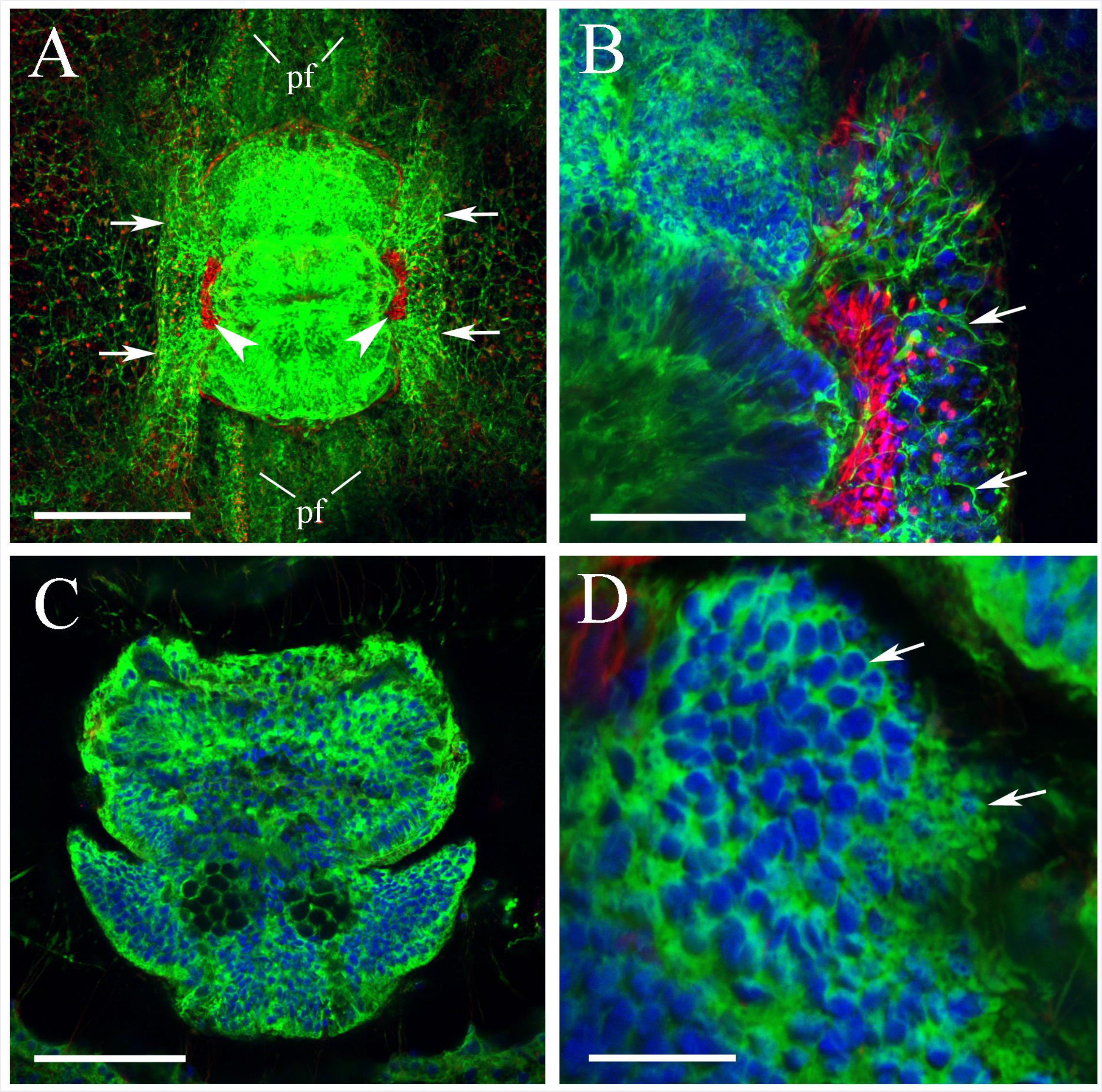
**The organization of the aboral organ** as revealed by tubulin antibody (green) and phalloidin (red) staining (nuclear DAPI staining is blue). **A**: A diffused neural net in the skin forms two invaginations (arrows) on both sides of the aboral organ. These areas also contain two symmetrical structures stained by phalloidin (arrowheads). **B**: Some neurons and their neurites including cells from the neural net (arrows) are located nearby the cluster of phalloidin-stained (red) small cells with processes. **C, D**: Bright tubulin-related fluorescence (green) of the aboral organ in all preparations is related to many very tightly packed small immunoreactive cells (some of them are indicated by arrows) across the entire aboral organ. Abbreviations: *pf* - polar field. Scale bars: A - 100 μm; B - 20 μm; C - 50 μm; D - 10 μm.

**Table 1.** Neuronal cell types identified in *Pleurobrachia*.

|  | *Neuronal Cell Type* | *Size* | *Location* | *Figure* |
| --- | --- | --- | --- | --- |
| 1. | Neuron-like cells in the aboral organ | 1-4 $\mu\text{m}$ | Aboral organ | Fig. 4 C, D |
| 2. | Neuron-like cells in the polar fields | 2-3 $\mu\text{m}$ | Polar Fields | Fig. 8 C, D |
| 3. | Bipolar neurons in the strands of the subepithelial net | 3-7 $\mu\text{m}$ | Subepithelial network | Fig. 10 C |
| 4. | Tripolar neurons in the nodes of the subepithelial net | 3-7 $\mu\text{m}$ | Subepithelial network | Fig. 10 B |
| 5. | Neurons located between strands of the subepithelial net | 4-8 $\mu\text{m}$ | Subepithelial network | Fig. 10 A, B |
| 6. | Neurons located between strands of the subepithelial net with cilia | 4-8 $\mu\text{m}$ | Subepithelial network | Fig. 11 A, B, C |
| 7. | Star-like mesogleal neurons with multiple radial processes | 3-7 $\mu\text{m}$ | Mesoglea | Fig. 17 B |
| 8. | Bipolar mesogleal neurons | 3-7 $\mu\text{m}$ | Mesoglea | Fig. 17 B |
| 9. | Mesogleal neurons projecting a single long neurite | 3-7 $\mu\text{m}$ | Mesoglea | Fig. 18 D, E, F |

Diffused neural network located within the body wall formed two deep foldings on both sides of the aboral organ (Fig. 4A, B, and Video-1 in Supplements). These two network foldings neural had a wide web of direct connections to the aboral organ (Fig. 4B). The areas, where diffused neural network joined the aboral organ, contained two narrow structures stained by phalloidin (F-actin marker), which included groups of small cells and fibers (Fig. 4A, B).

Also, phalloidin labeled four cylindrical structures at the base of each polar field (Fig. 5A, B). These “cylinders” consisted of many parallel, very thin, phalloidin-labeled filaments, and looked like ‘anchors’ to which polar field bands were attached. In the center of the aboral organ, phalloidin labeled four groups of muscle filaments related to fours balancers (Fig. 5B).

**Figure 5.**
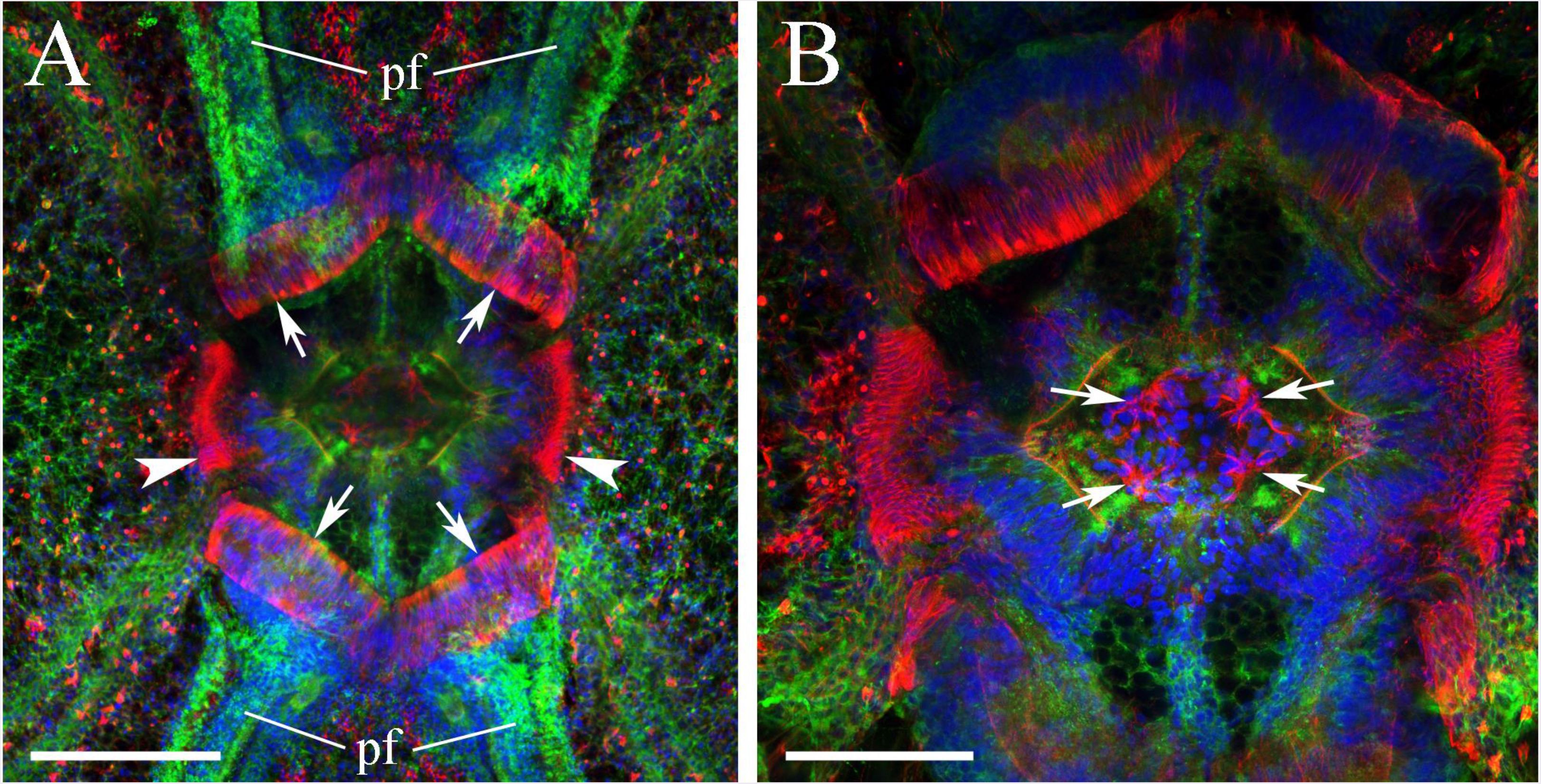
**Horizontal optical section of the aboral organ within the plane of the polar fields** (tubulin IR - green; phalloidin – red; and DAPI - blue). **A**: In addition to previously described phalloidin-labeled groups of cells and fibers on both sides of the aboral organ (arrowheads) there are also four cylindrical structures at the base of each polar field stained by phalloidin (arrows). They all consist of many parallel phalloidin-labeled filaments with F-actin. **B**: In the center of the aboral organ phalloidin labels four balancer-related structures (arrows), while tubulin IR presumably reveals four lamellate bodies and a “bridge” consisting of small cells (Tamm and Tamm, 2002) and connecting the aboral organ and polar fields see also Fig. 3C, D). Abbreviations: *pf* - polar field; *l* - lamellate body; *b* - bridge. Scale bars: A - 50 μm; B - 25 μm.

Phalloidin is known to be a good marker of muscle cells (Wulf et al., 1979). It stained a group of very thin and short fibers connecting the aboral organ to the walls of two short anal canals in *Pleurobrachia* (Fig. 6B). There were also several long individual muscle filaments under the aboral organ and around the anal canals (Fig. 6C). Finally, several thick muscle fibers were attached to the polar fields; these fibers presumably mediate the defensive withdrawal of polar fields inside the body. SYTOX green staining revealed multiple nuclei in each muscle filament (Fig. 6D).

**Figure 6.**
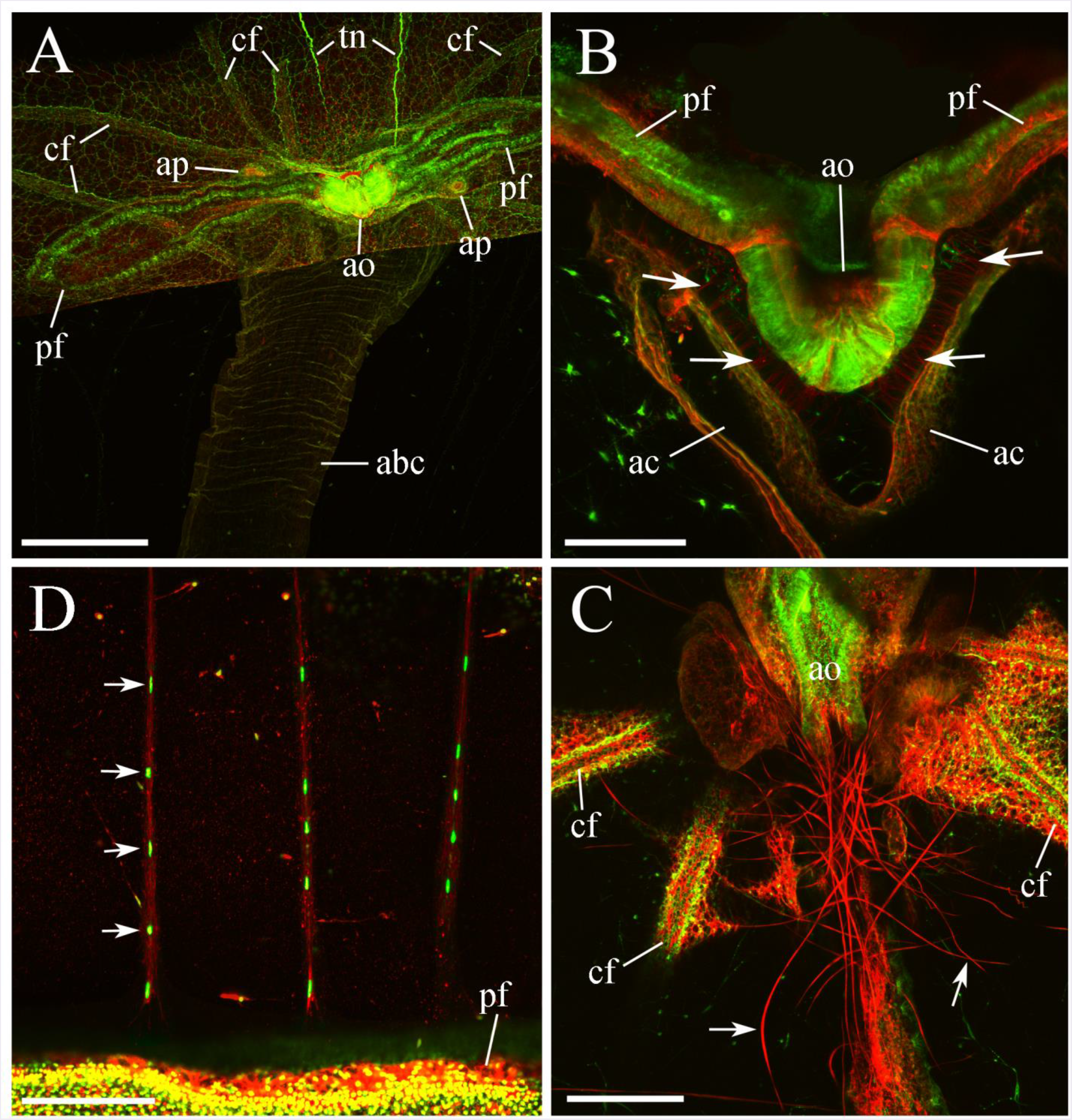
The aboral complex in *Pleurobrachia*. **A: The general view of the aboral complex** (tubulin IR – green, and phalloidin - red). The photo shows two elongated polar fields, ciliated furrows, two tentacular nerves, a wide aboral canal splitting into two anal canals and terminating as two anal pores, with muscle sphincters, on opposite sides of the aboral organ. **B: Sagittal view of the aboral organ.** Phalloidin stains a group of very thin and short muscles (arrows) connecting the aboral organ to the walls of two short anal canals. **C**: A group of long muscle cells (arrows) under the aboral organ. **D**: Several muscle cells with many nuclei are attached to the polar fields. SYTOX green was used to stains nuclei (arrows). Abbreviations: *ao* - aboral organ; *pf* - polar field; *abc* - aboral canal; *ac* - anal canal; *ap* - anal pore; *cf* - ciliated furrow; *tn* - tentacular nerve. Scale bars: A - 300 μm; B - 100 μm, C - 150 μm, D - 50 μm.

#### Neural cells are located in the polar fields

The length of the polar fields always directly corresponded to the age and the size of the animals (Fig. 7). In very young animals, the polar fields are very short and small, and sometimes cannot even be seen on the surface – they are inside the body-wall invagination where the aboral organ is located (Norekian and Moroz, 2016). After hatching, the polar fields grow in length to form an elongated figure eight structure.

**Figure 7.**
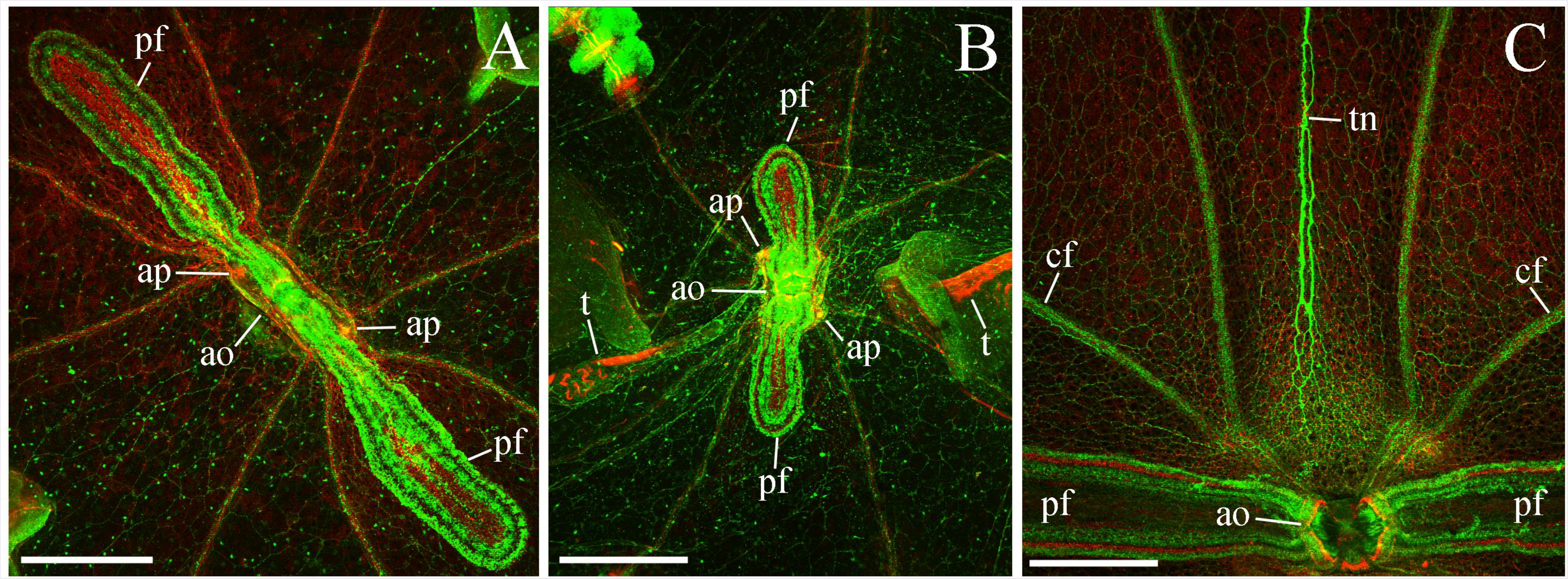
**The organization of the polar fields** (tubulin IR – green, phalloidin – red). There is a direct correlation between age and the size of the polar fields. In adults, the polar fields are large and elongated (**A**), while in young animals the polar fields are more compact (**B**) but with the same organization pattern. **C**: While approaching the aboral organ, tentacular nerve makes multiple branches and merges with the subepithelial network. Abbreviations: *ao* - aboral organ; *pf* - polar field; *cf* - ciliated furrow; *ap* - anal pore; *tn* - tentacular nerve; *t* - tentacle. Scale bars: A, B, C - 300 μm.

The tubulin antibodies intensely stained two bands of numerous short cilia at outside edges of the polar fields (Fig. 7 and 8A, B), efficiently ‘masking’ neural elements inside the structure. However, using the confocal microscopy for isolating optical sections, we identified densely packed neurons through the entire length of the polar fields (Fig. 8B, C; Table 1). These small cells (less than 3 μm in diameter) had short neurites (Fig. 8D) and formed a narrow band next to the outside edge of the polar fields. Many neuron-like cells were bipolar, while others had three or four elongated processes, primarily oriented to the inner parts of the polar fields.

**Figure 8.**
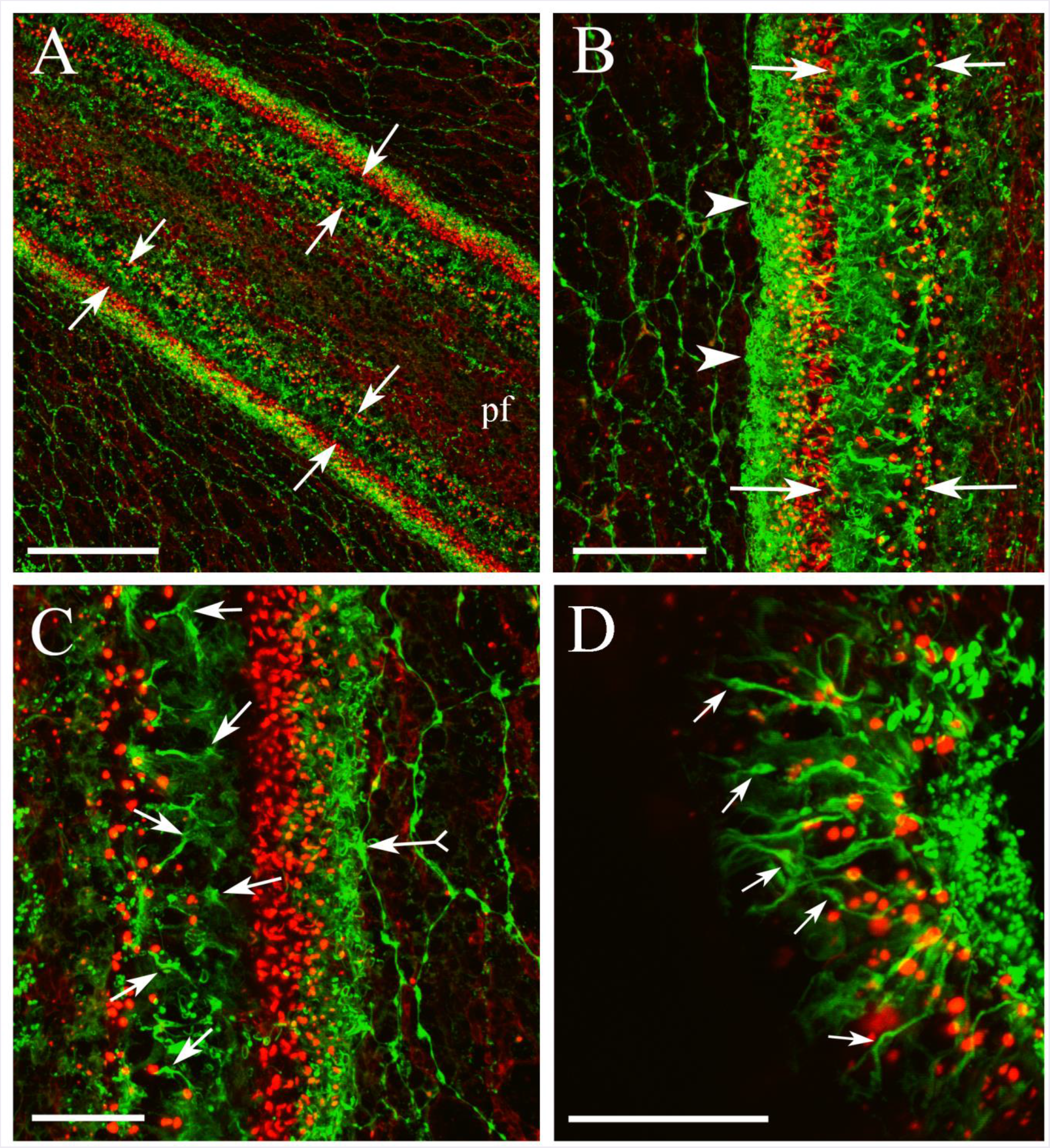
**Neurons in the polar fields** *(pf)* (tubulin IR – green, phalloidin – red). **A, B**: A narrow band next to the outside edge of the polar fields (arrows) contains densely packed tubulin IR neural cells through its entire length. Tubulin antibody also stained numerous short cilia covering the surface of the polar fields (arrowheads). **C, D**: Neurons (arrows) are small (3 μm in diameter) with short processes predominantly toward the inside area the polar fields. The subepithelial polygonal neural network makes a direct connection with the polar fields (arrow with tail). Scale bars: A - 100 μm, B - 40 μm, C, D - 20 μm.

The epithelial polygonal neural network had numerous connections with the polar fields (Fig. 8C). Nevertheless, we could not visualize a direct contact between the polygonal network and neurons inside the polar field because of the brightly stained cilia on edge.

### 3.3 Two Neural Subsystems in *Pleurobrachia bachei*

Tubulin IR revealed two major parts of the diffused nervous system in *Pleurobrachia*: (i) the surface, subepithelial polygonal neural network, and (ii) a large diffuse meshwork of neurons and fibers in the mesoglea, which spread over the internal gelatinous part of the body.

#### 3.3.1 The Subepithelial Neural Net

The subepithelial neural network is a large polygonal mesh of neurites covering the entire body surface under the epithelium, as well as at the surface of the tentacle pockets. This meshwork consisted of individual polygonal units of different sizes (between 20 and 150 μm) and shapes: many units had four ‘corners’, but there were also pentagonal (five corners), hexagonal (six corners), heptagonal (seven corners), and sometimes octagonal (eight corners) configurations (Fig. 9A, B). Higher resolution microscopy identified many tightly packed neuronal processes (Fig. 9B). and direct connections between the subepithelial network and other surface structures such as the polar fields and ciliated furrows (Fig. 9C, D).

**Figure 9.**
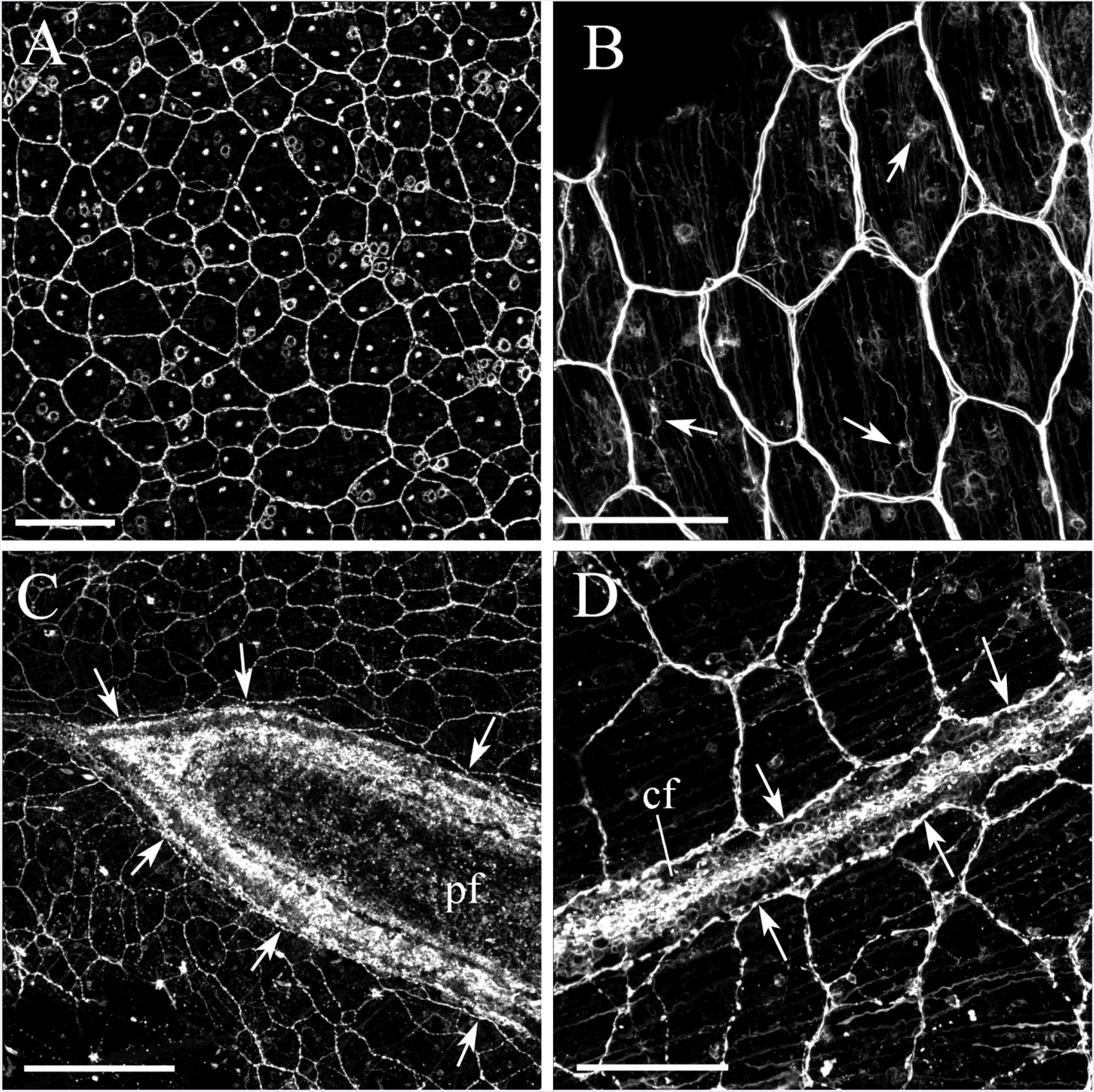
Subepithelial neural network stained with tubulin antibody. **A, B**: Neural net consists of polygonal units of different shapes and sizes. There are individual neurons with clearly visible neurites (arrows). The segments of the neural net composed of tightly packed thin processes, rather than a single thick axon (B). **C, D**: The neural net covers all surface of the body wall, and neuronal terminals contact both the polar fields (C) and ciliated furrows (D), sometimes tightly enclosing them (shown by arrows). Abbreviations: *cf* - ciliated furrow; *pf* - polar field. Scale bars: A - 100 μm, B - 50 μm, C - 120 μm, D - 50 μm.

The subepithelial network is formed by 4-8 μm neurons with multiple processes, which contribute to the structure of the polygonal units (Fig. 10A, B; Table 1). All neurons had a single nucleus (Fig. 10A, B), but DAPI staining also revealed numerous nuclei inside the meshwork itself suggesting that neuronal somata with little cytoplasm area were tightly packed (Fig. 10C). Some of these cells were located in the nods of a mesh and appeared to be tripolar, while others were bipolar (Table 1).

**Figure 10.**
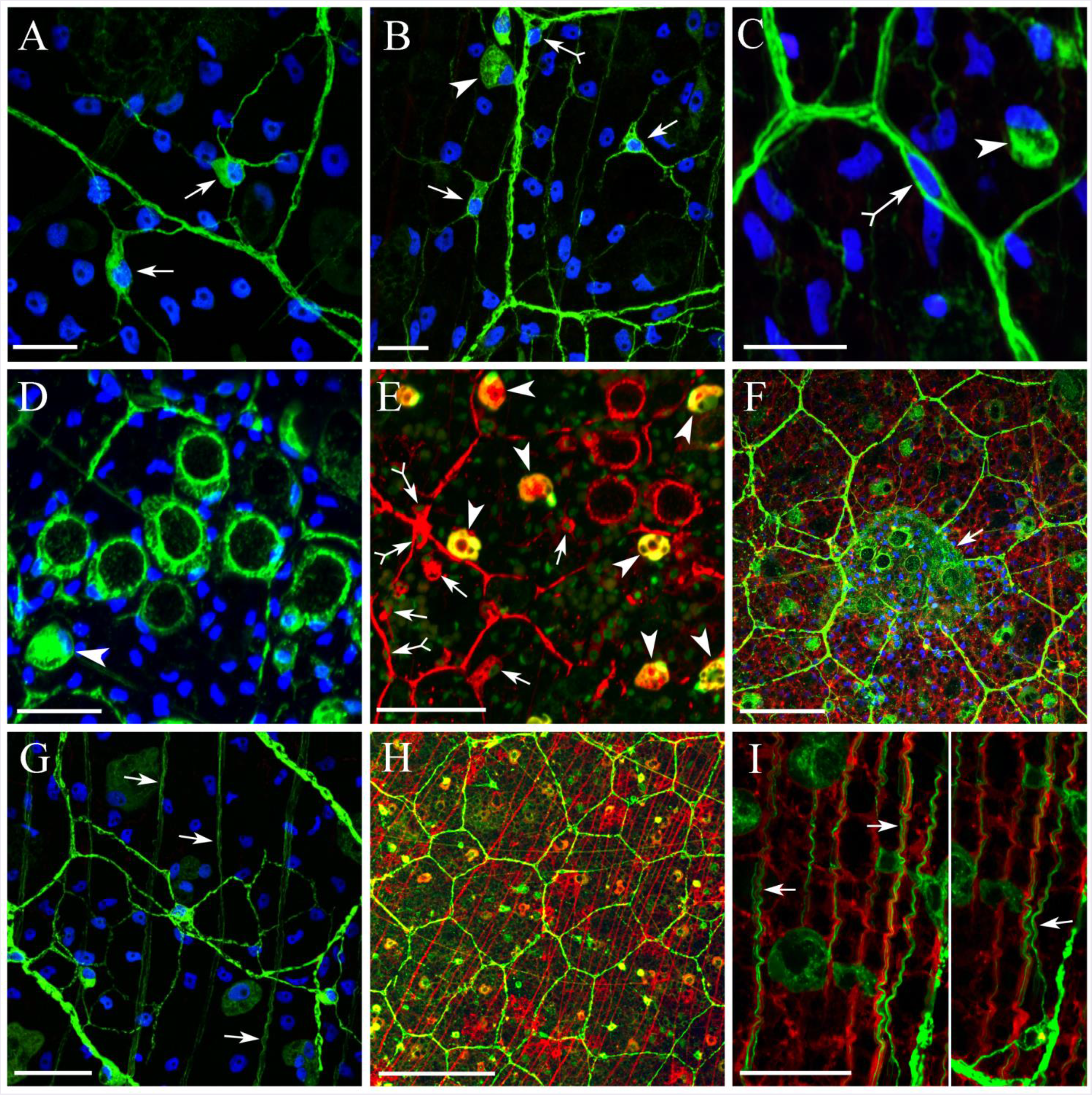
**Distinct elements of the subepithelial neural network** labeled by tubulin antibody (in all photos nuclei were stained by DAPI (blue) except E, where we used SYTOX green). **A, B**: The subepithelial network consists of individual neurons with some cell bodies outside of the segments of the main nerve net (arrows). Most neurons have multiple processes, which contribute to the polygonal architecture. **C**: DAPI staining shows examples of a neuron inside the segment of the polygon (arrowhead with tail) with the central nucleus. **D**: Tubulin antibody also stains several secretory cells at different functional stages. At earlier secretory stages the cells are round-shaped and more brightly stained (arrowhead). At later stages, they look like large ‘pores.’ The secretory cells are frequently grouped in clusters of 2 to 8. **E**: When we used SYTOX green as a nuclear marker instead of DAPI, together with tubulin antibody coupled with a red tag, some secretory cells have the orange or yellow appearance (arrowheads) suggesting the co-localization of both markers. SYTOX green also labels a high concentration of RNA in cell bodies, which might occur at the stage of intense protein synthesis before the secretion. In this image neurons with neurites shown in red (arrows). **F**: In preparations, where the upper epithelial layer was also scanned, a slightly fluorescent cloud (arrow) was detected over the clusters of ‘empty’ secretory cells with pore-like appearance (see D), further suggesting that cells might be captured following the release of their content during fixation. **G**: A part of the subepithelial neural net is a web of very fine processes (arrows) running in parallel to each other slightly below the main neural mesh. **H**: Long individual parietal muscle fibers stained by phalloidin (red) have the same location, slightly below the neural mesh, and similar parallel orientation with fine neural processes described in G. **I**: Higher resolution image of the double labeling of both neural processes (green) and muscles (red) shows that individual neural processes are attached to each muscle fiber following its entire length (arrows). Note that one neurite of a single bipolar neuron enters the subepithelial neural net, and might function as a motoneuron. Scale bars: A, B, C, D - 10 μm, E - 25 μm, F - 50 μm, G - 20 μm, H - 100 μm, I - 20 μm.

Our data suggest that the subepithelial network controls contractions of the surface muscles and body wall movements. Indeed, we identified a web of very fine neuronal processes running in parallel to subepithelial muscles and connected to each other slightly below the focal plane of the principal network (Fig. 10G, H). Moreover, higher resolution double labeling confirmed that several individual neural processes are located at the surface of single muscle fibers over their entire length (Fig. 10I), and we traced these processes to some neurons located between polygonal neurites of the subepithelial network.

In addition to the small neurons with well-defined neurites, we identified larger secretory-like cells (5-10 μm in diameter) of a round shape and without processes - glandular or gland cells (Hernandez-Nicaise, 1991); Fig. 10C, D). These secretory cells were distributed randomly or grouped in clusters of 3 to 8 cells. We think these cells were captured at different secretory stages. At their presumed last secretion phase (e.g., after emptying their content), the cells looked like large aperture-shaped structures with a large circular dark center and approximately 7-10 μm in diameter (Fig. 10D, E).

When SYTOX green was used to stain the cell nuclei instead of DAPI, while tubulin antibody was tagged with a red fluorescent label, the difference between neurons with processes and round-shaped glandular cells became even more obvious. Glandular cells appeared to be orange or yellow, unlike neurons which remained red, apparently as a result of co-localization of tubulin IR and diffuse SYTOX green staining (Fig. 10E). SYTOX green labels not only DNA but also detects a high concentration of RNA in cell bodies, suggesting that secretory cells have much higher RNA concentration (before they empty their content). In some preparations, when we scanned the upper epithelial layer, a slightly auto-fluorescent cloud was observed over the clusters of ‘empty’ secretory cells, thus suggesting that they released their content during fixation, and further supporting our hypothesis that they are indeed secretory cells (Fig. 10F).

#### 3.3.2 Identification of putative sensory cells in the intergument

We identified five types of putative sensory cells in the skin of *Pleurobrachia*: two types were integrated parts of the subepithelial neural network with predominant tubulin IR, and three types represented the epithelial receptors (Fig. 11 and Table 2). SEM observations also revealed several different classes of sensory cilia on the body surface of *Pleurobrachia*, with some of them corresponding to these receptor cell types (Table 2 and Fig. 12).

**Figure 11.**
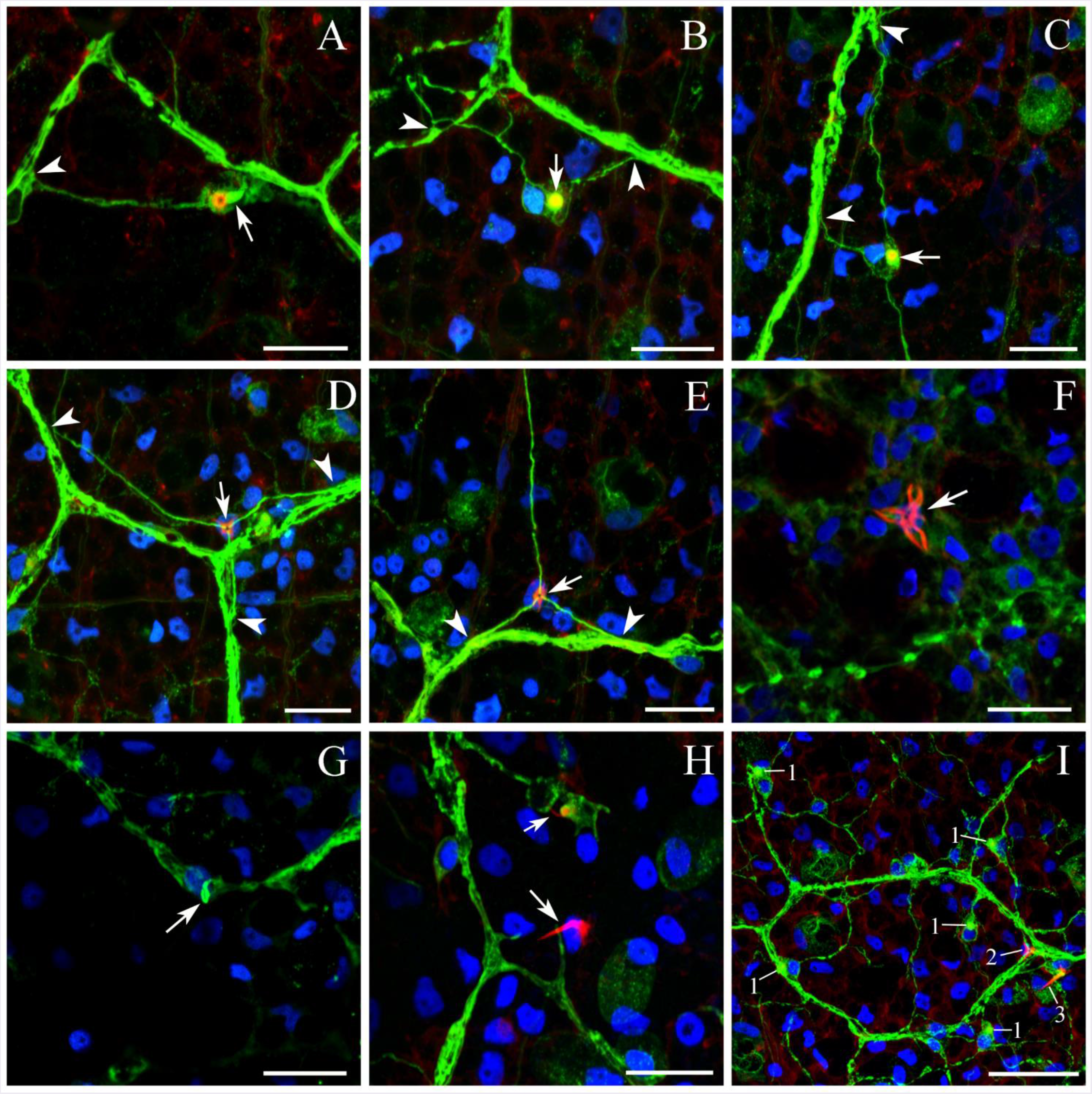
**Five types of sensory cells associated with the subepithelial neural net** (green – tubulin IR; red – phalloidin; blue – nuclear DAPI labeling). **A, B, C**: The first receptor type includes cells with a single, short, thick cilium stained with tubulin antibody (arrows). The base of these cilia is always labeled by phalloidin (red). The overall receptor morphology (with two or three processes within the neural net - arrowheads) and labeling patterns are similar to many neurons in the net (DAPI staining is absent in A). **D, E**: The second type of receptor cells has three radial neurites that are entering the subepithelial neural net (arrowheads). **F**: The third type of receptors contains a compact group of 2 to 7 long cilia labeled by phalloidin. The cilia always originated from one central region (arrow) located on the top of a nucleus. **G**: The forth type of receptors has a single long cilium, which is curled at the top (arrow), and labeled by tubulin antibody only (arrow). **H**: The fifth receptor type has a long cilium labeled by phalloidin only. **I**: Spatial relationship among different receptor types in the subepithelial network: (1) - neuron-like cell with tubulin-stained cilia; (2) - a cell with three radial neural processes joining the network; (3) - a cell with a long phalloidin-labeled cilium. Scale bars: A to H - 10 μm, I - 20 μm.

**Figure 12.**
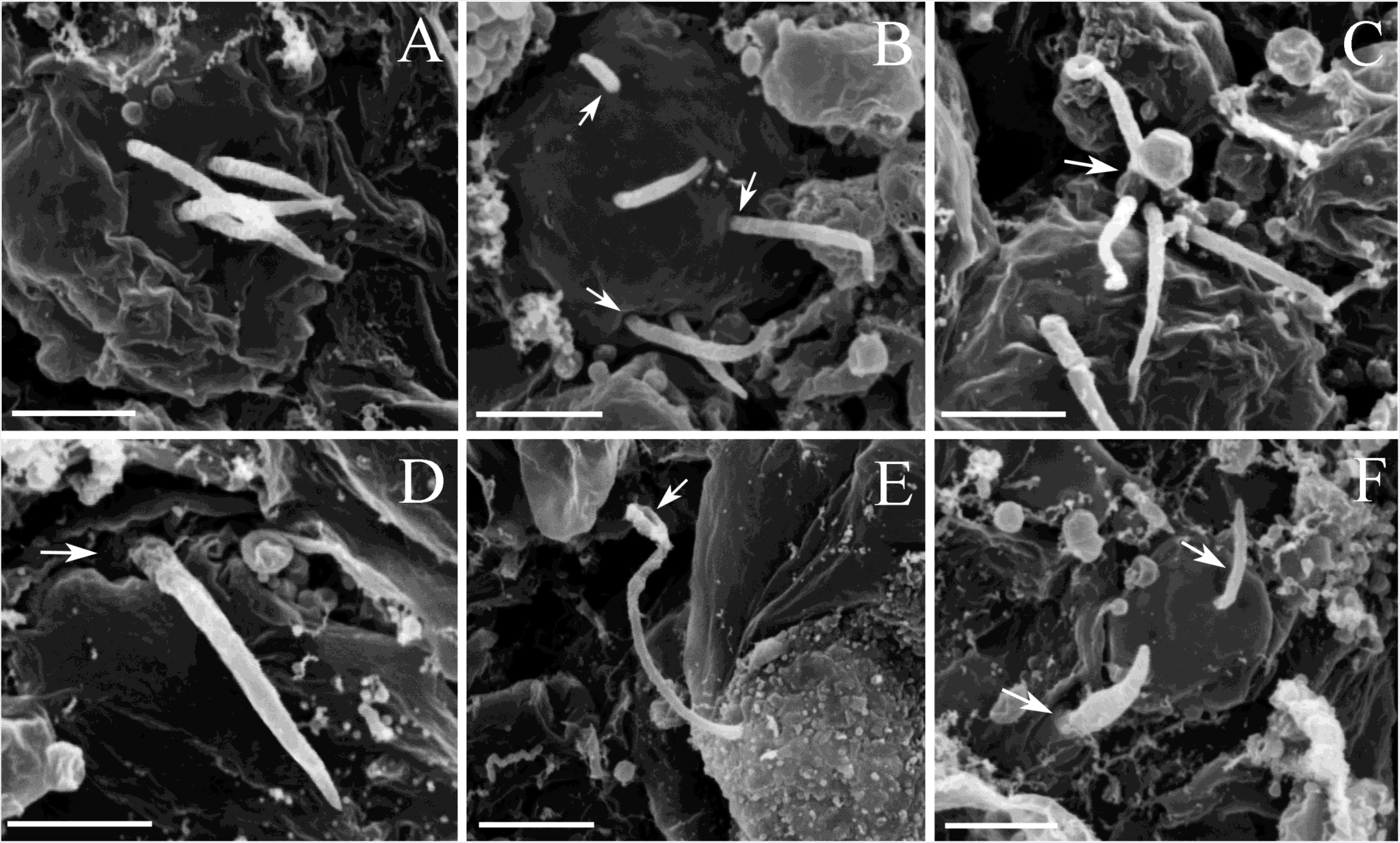
SEM of different ciliated receptors on the surface of the body. **A, B**: Receptors with 2 to 7 small cilia on the top of a single round-shaped cell. Although the length of individual cilia in the group could vary, they all have the same thickness and morphology. **C**: An image of a damaged cell with the base where all cilia from the group join together (arrow). **D**: The receptor with a large cilium and a wide base (arrow). **E**: A cell with a cilium curled on the top (arrow). **F**: Arrows point to individual cilia with different morphology. Scale bars: 2 μm.

**Table 2.** Receptor/sensory cell types identified in *Pleurobrachia*.

|  | *Receptor type* | *Location* | *Number of cilia* | *Visualization* | *Figures* |
| --- | --- | --- | --- | --- | --- |
| 1. | Ciliated neurons between strands of the subepithelial net | Subepithelial; a part of the neural net | 1 cilium | Tubulin AB | Fig. 11 A, B, C |
| 2. | Receptors with tubulin AB labeled processes joining the subepithelial net | Subepithelial; a part of the neural net | ? | Tubulin AB, only processes labeled | Fig. 11 D, E |
| 3. | Receptor with a group of 2-7 cilia | Skin | 2-7 cilia | Phalloidin & SEM | Fig. 11 F |
| 4. | Receptor with a long and curled cilium | Skin | 1 cilium | Tubulin AB & SEM | Fig. 11 G |
| 5. | Receptor with a large thick cilium | Skin | 1 cilium | Phalloidin & SEM | Fig. 11 H |
| 6. | Ciliated receptors on the surface of the tentacles | Tentacle surface | 1 cilium | Phalloidin | Fig. 25 C, D |

The first type of receptors has a relatively thick and short cilium, which brightly stained by tubulin antibody (Fig. 11A, B, C; very rarely, we observed two or three cilia). A wide base root of the ciliated structure was always labeled by phalloidin suggesting the presence of F-actin. The morphology of these cells was very similar to most neurons described in the previous section (e.g., Fig. 11H). They had cell bodies about 5 μm in diameter and usually two or three processes fully integrated within the subepithelial neural net (Fig. 11A, B, C, I; though sometimes there was one or more than three). Also, we observed with SEM many short surface cilia (Fig. 12F). Some of them could belong to the first receptor type described here. However, there was no sufficient information to establish an irrefutable link between them.

The second receptor type included cells located deeper in the body wall, just under the layer of the subepidermal neural network (presumably proprioreceptors). These cells produced three thin neurites (labeled by tubulin antibody), which spread in the radial directions (Fig. 11D, E) and fused with the rest of the network. Very often these receptors were located next to the joints points (‘corners,’ Fig. 11D), and more rarely, receptors had only two visible processes instead of three. Interestingly, the cell body itself of this receptor type was labeled neither by phalloidin nor tubulin antibody. However, a few short phalloidin-stained structures were always located on the top of their DAPI stained nucleus (Fig. 11D, E).

We recognized the third type of receptors as 5 to 7 μm cells with a compact group of 2-7 cilia labeled by phalloidin only (Fig. 11F). The length of individual cilia in the group could vary, but they all had the same thickness and appearance under SEM (Fig. 12A, B, C). These cilia were 3 to 8 μm long and always originated from one central region close to the nucleus. The base where all cilia from the group were joining each other was also revealed by SEM (Fig. 12C).

The fourth type of receptors had a long thick single cilium (up to 7-10 μm) that was labeled by phalloidin only (Fig. 11H, I and Fig.12D).

In contrast, the fifth type of receptors also had a single long cilium, which was labeled by tubulin antibody only (Fig.11G), but with a small base labeled by phalloidin. The very top of the cilium was always curled and looked very similar in SEM preparations (Fig. 12E). The last two receptor types were less abundant compared to others.

#### 3.3.3 Tentacles and their neuro-muscular system

The main food capturing organs in *Pleurobrachia* is the highly mobile tentacles with dozens of tentilla (or tentillae, “little tentacles”) (Figs. 1, 13, 14A). The base of each contractile tentacle is located inside the tentacle pocket, which itself is attached to the posterior pharynx forming the single functional unit (Fig. 14A). Both tentacles and tentilla were completely covered with colloblasts (Fig. 13C, D), glue-like cells that capture prey. As a result, the tentacles had a complex neuro-muscular organization with sensory capabilities and innervation.

**Figure 13.**
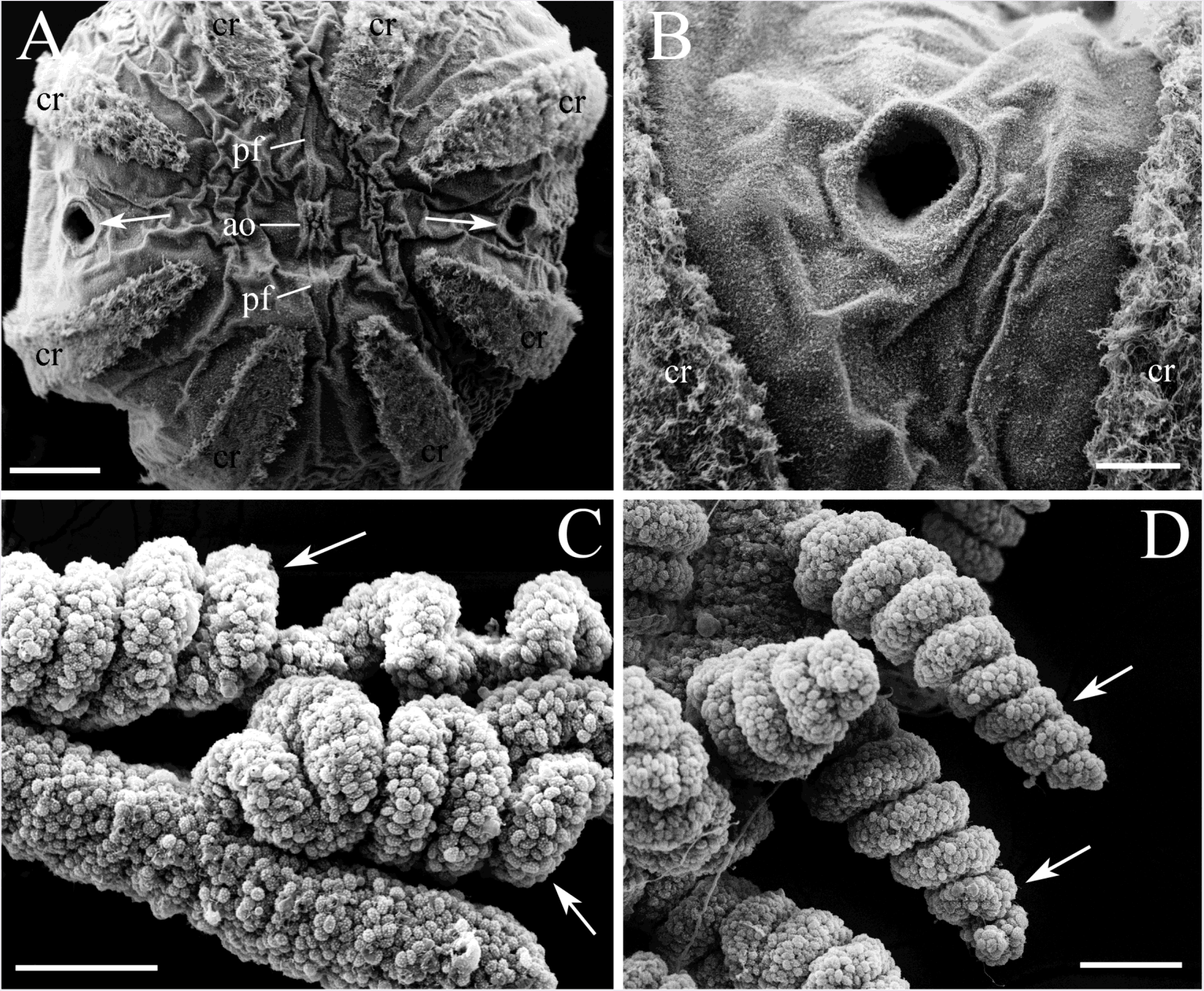
SEM of the tentacles. **A, B**: Narrow openings of the tentacle pockets (arrows) are located on the aboral side between comb rows. **C, D**: Tentacles, and tentilla are fully covered with small round-shaped colloblasts (arrows). Abbreviations: *ao* - aboral organ; *pf* - polar field; *cr* - comb row. Scale bars: A - 1 mm, B - 200 μm, C, D - 50 μm.

**Figure 14.**
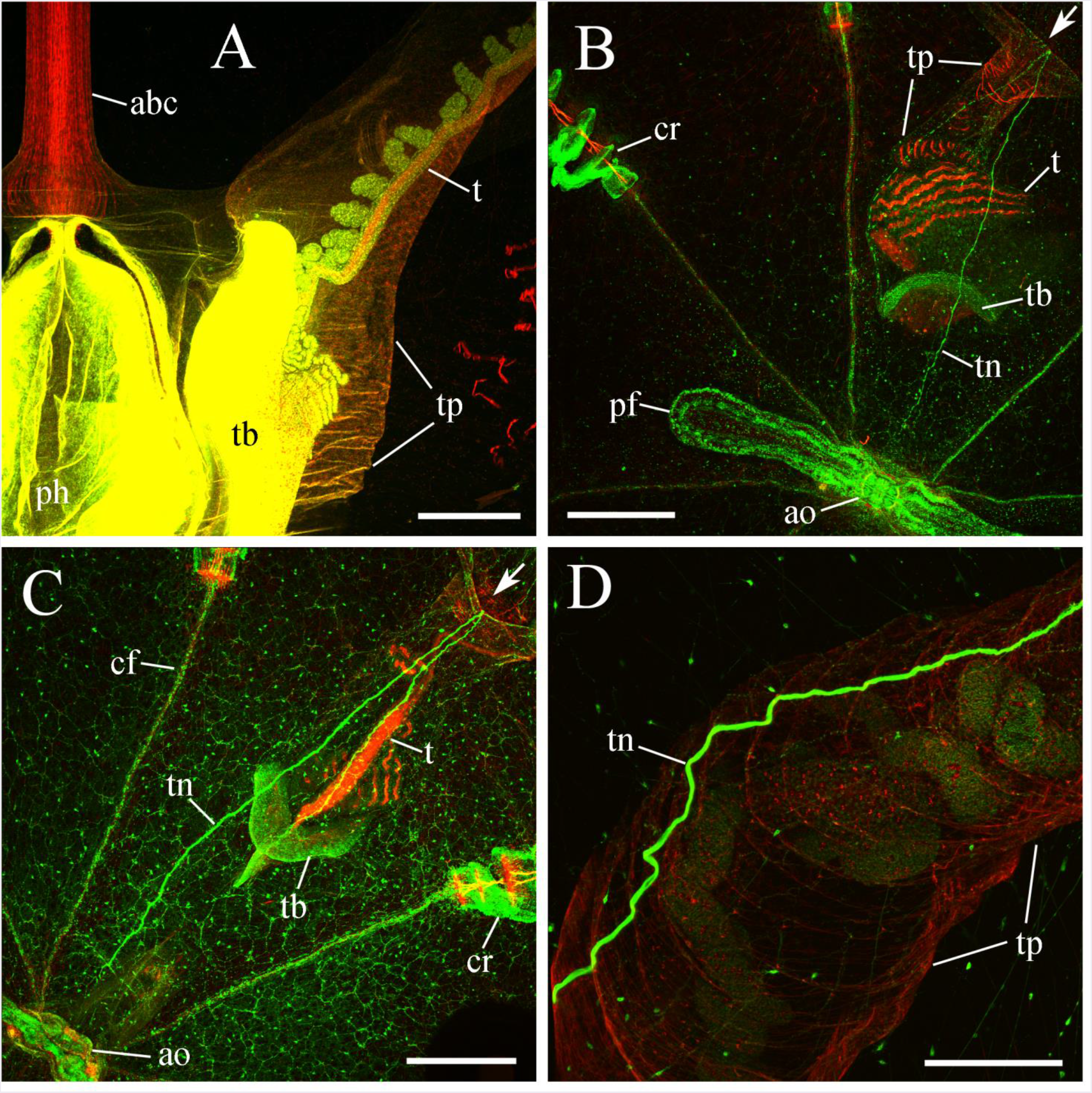
**Aboral tentacular nerve** with tubulin antibody (green) and phalloidin (red) staining. **A**: Each tentacle is located inside a tentacle pocket, which is attached to the posterior pharynx in the center of the animal. SYTOX green provided a yellow background fluorescence in addition to phalloidin staining for this visualization. **B, C**: The aboral tentacular nerves originate nearby of the aboral organ, then run on the surface between the comb rows and turn into the opening (arrow) of the ipsilateral tentacle pocket. The nerves are visualized through the entire length of a tentacle pocket and innervate the base of each tentacle. **D**: Aboral tentacular nerve in the wall of the tentacle pocket at higher magnification. Abbreviations: *ao* – the aboral organ; *abc* – the aboral canal; *ph* – the pharynx; *pf* – the polar field; *cr* - comb rows; *cf* - ciliated furrow; *tb* - tentacle base; *tp* - tentacle pockets; *tn* – tentacular nerve; *t* - tentacle (B and C also show phalloidin-stained muscles in the main tentacle and tentilla). Scale bars: A - 500 μm, B, C - 250 μm, D - 150 μm.

We identified four major nerves in *Pleurobrachia* - two on each side of the animal. All of them are associated with the tentacles and, therefore, were named as the tentacular nerves (Hernandez-Nicaise, 1991). The base of each contractile tentacle is located inside the tentacle pocket, which itself is attached to the posterior pharynx forming the single functional unit (Fig. 13A).

Two symmetrical *aboral* tentacular nerves can connect tentacles and the aboral complex (Fig. 3A, 7C, 14B, C), where nerves began to branch, and gradually merged with the subepithelial network (Fig. 7C). On the opposite side, after crossing half of the body, the tentacle nerves ran into the ipsilateral tentacle pocket (Fig. 14B, C, D), and eventually innervated the base of each tentacle (Fig. 14B, C, D and Video-2 in Supplements).

Two *oral* tentacular nerves originated from the surface network near the mouth. These nerves ran between comb rows to the ipsilateral tentacle pocket, where they branched, and ultimately merged with the subepithelial neural network at the opening of the pocket (Fig. 15A, B, C). Unlike the aboral tentacular nerves, the oral tentacular nerves did not run through the entire tentacle pocket (Fig. 15A, B, C).

**Figure 15.**
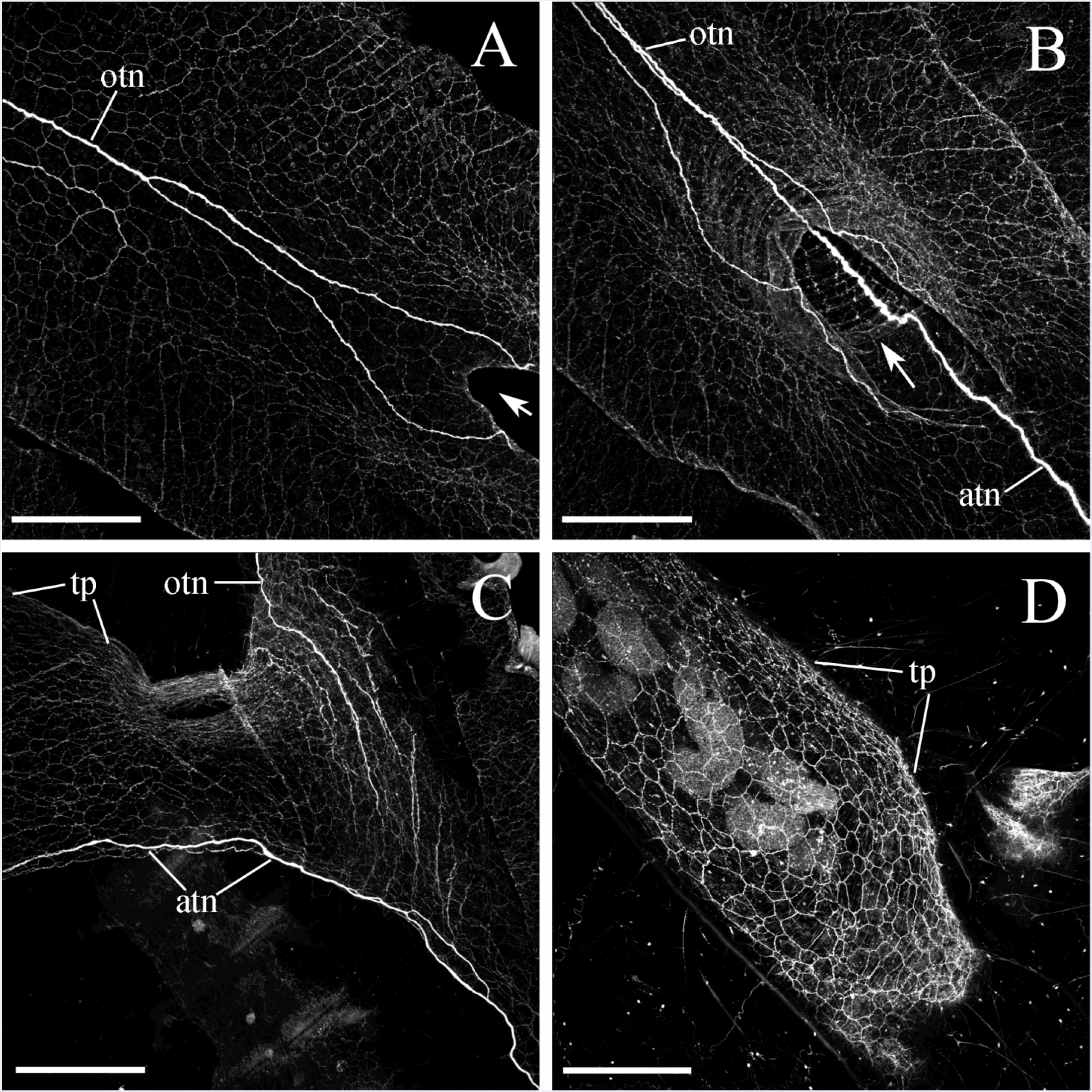
**Aboral and oral tentacular nerves with tubulin antibody staining**. Two *oral* tentacular nerves that originate on both sides of the mouth reach the ipsilateral tentacle pocket opening (arrow), but unlike the *aboral* tentacular nerves, they don’t go deep inside and cross through the entire tentacle pocket to innervate the base the tentacles. The nerves start branching and thinning near the tentacle pocket opening and eventually merge with the neural network. **A, B**: Horizontal view. **C**: Saggital view. Note that all animal body surface area is covered by the polygonal neural network. **D**: The entire surface of each tentacle pocket is also covered with the same polygonal neural network. Abbreviations: *atn* - aboral tentacular nerve; *otn* - oral tentacular nerve; *tp* - tentacle pocket. Scale bars: A, B, C, D - 300 μm.

Each tentacular nerve was localized within the same layer as the subepithelial network and had numerous interconnections with it through its entire length. Thus, the tentacular nerves appeared to be an integrated part of the entire surface network, possibly collecting information from the mouth, the aboral organ, body surface and tentacles to control the feeding, locomotion and defense responses, which are tightly correlated with tentacle movements.

The tentacle pockets themselves contained a dense subepithelial network (Fig. 15D). In several SEM preparations (n=7), we identified neurons formed different polygonal units with clearly visible neural cell bodies and their processes (Fig. 16).

**Figure 16.**
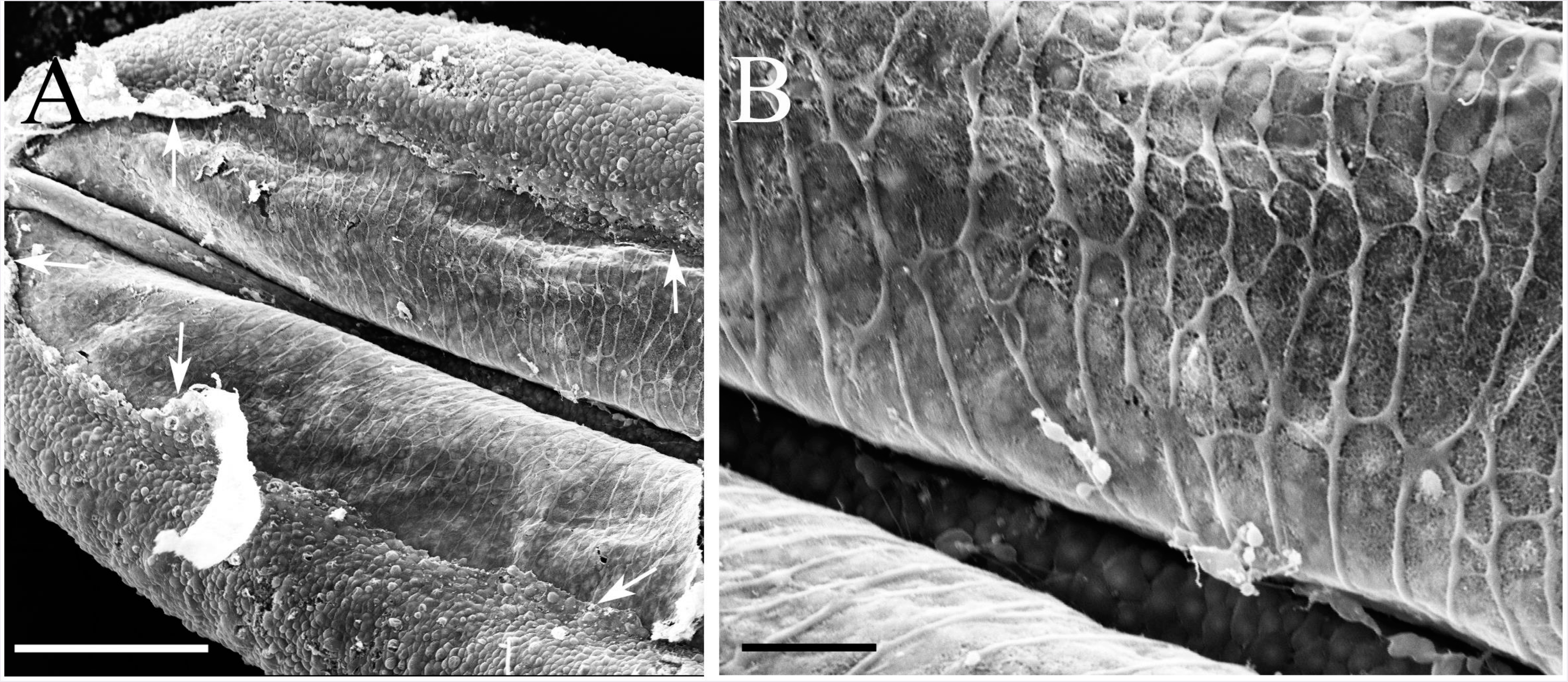
SEM of the polygonal neural network in the tentacle pocket. **A**: The upper epithelium layer is peeled off in some areas (arrows) revealing the underlying layer of the neural network. **B**: The neural network consists of different polygonal units and neural processes. Scale bars: A - 100 μm, B - 20 μm.

Each tentacle itself contained a central nerve (Fig. 17B) with numerous neuronal processes, which ran parallel to the muscles through the entire length (Fig. 17C). In addition to numerous colloblasts, the surface of the tentacles was covered with ~5 μm receptors with a single cilium; each receptor sent a neural processes to the central nerve (Fig. 17C, D; Table 2).

**Figure 17.**
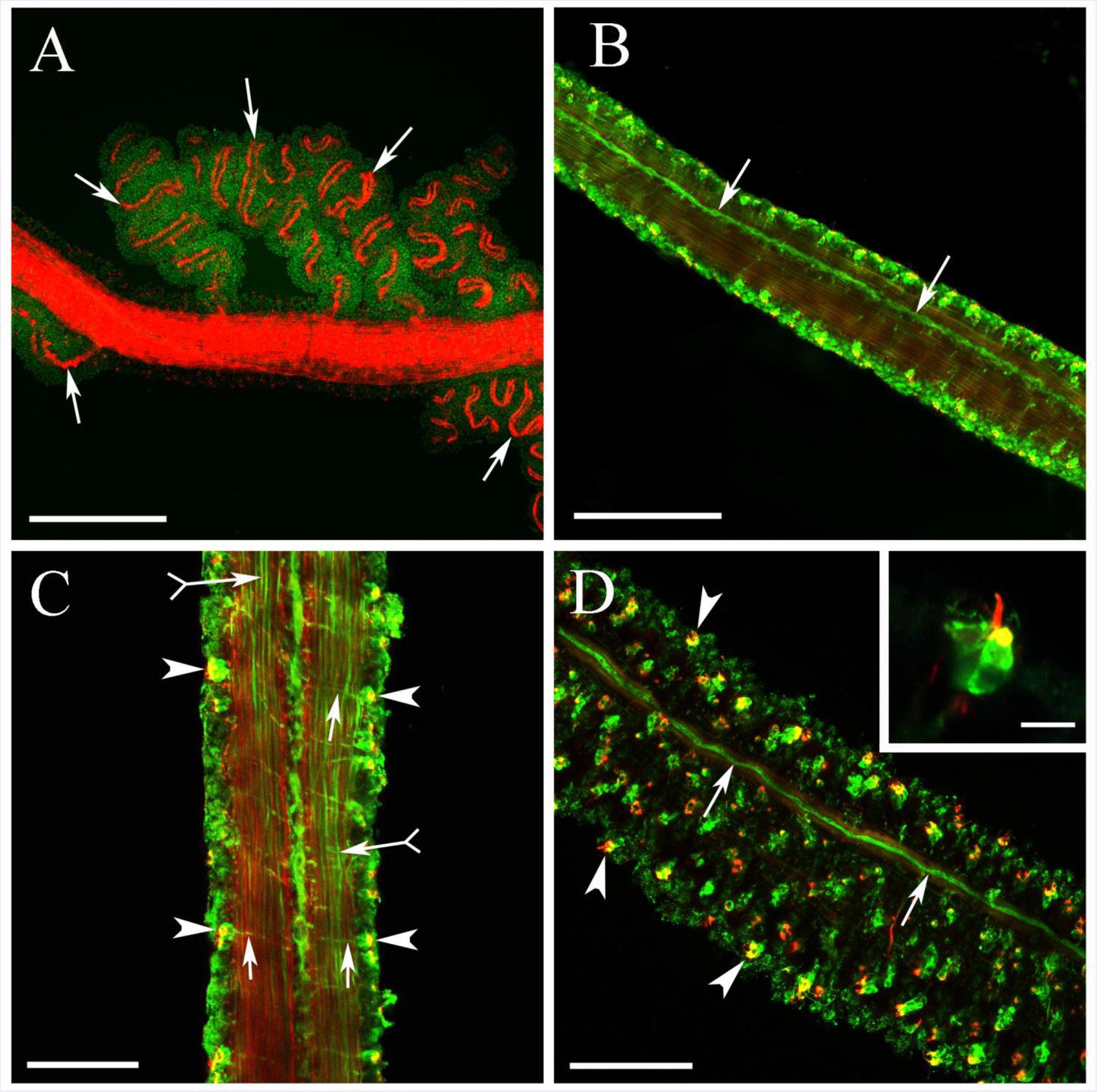
Neuro-muscular systems in the tentacles. **A**: Phalloidin (red) stains the entire main trunk of each tentacle suggesting its muscular nature. It also labels muscle fibers in each tentilla (arrows). Each tentilla has a double strain of muscle fibers, which is usually packed in tight spirals. **B**: The main trunk of each tentacle has a central nerve (arrows) along its entire length (stained by tubulin antibody - green). **C**: Tubulin IR reveals: (i) numerous neuronal processes (arrows with tail) running along the muscle fibers through the length of the tentacle; (ii) receptor cells on the surface (arrowheads). Each of those receptors sends a neurite to the central nerve (arrows). **D**: The tentacle surface is densely covered by these receptors (arrowheads). Large magnification (insert) shows that each receptor cell consists of a round cell body labeled by tubulin antibody (green) and a short cilium labeled by phalloidin (red). Scale bars: A - 150 μm, B - 50 μm, C - 30 μm, D - 30 μm (insert - 5 μm).

Phalloidin staining confirmed the muscular natures of tentacles, although tentilla had different types of double strain muscles (Fig. 17A).

#### 3.3.4 Mesogleal neural system

The entire space between the skin and inner gastroderm in *Pleurobrachia* is occupied by mesoglea, a jelly-like, fully transparent tissue, characteristic of the phylum. Both phalloidin and tubulin antibody labeled very diverse systems of muscles and neurons in the mesoglea; many of these cells had processes, which crossed the mesoglea in all directions (Fig. 18). The vast system of mesogleal neurons represented the second major part of the nervous system in *Pleurobrachia*.

**Figure 18.**
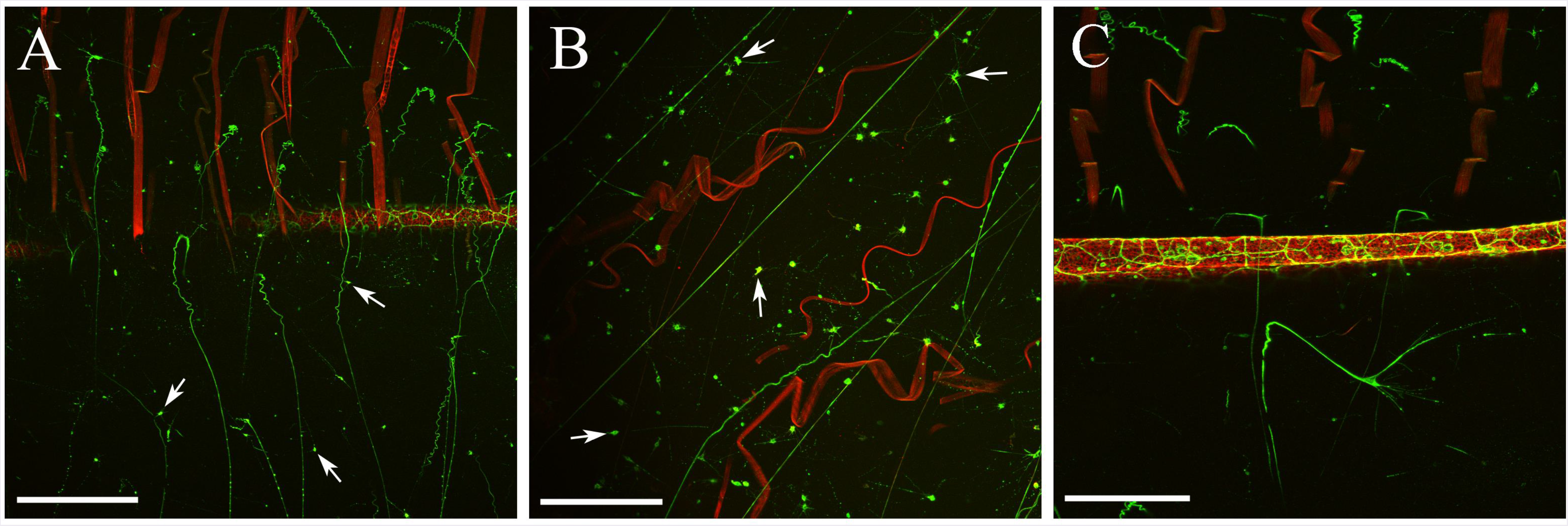
**A, B, C**. The mesoglea areas are very rich in muscle and neural cells as well as their processes. Phalloidin (red) stains the large muscle cells attached to comb rows, as well as thinner muscle cells and processes crossing the mesoglea. Tubulin antibody (green) label many small neuronal cell bodies (arrows) as well as thin neuronal-like processes crossing mesoglea in different directions. Scale bars: A - 300 μm, B, C - 150 μm.

There were approximately 2 to 3 thousand mesogleal neurons in the entire adult *Pleurobrachia* according to our estimates (Fig. 19A). All cell bodies had similar sizes between 3 μm and 7 μm in diameter (for the 1-1.5 cm animals) and could be separated into three types (Table 1).

**Figure 19.**
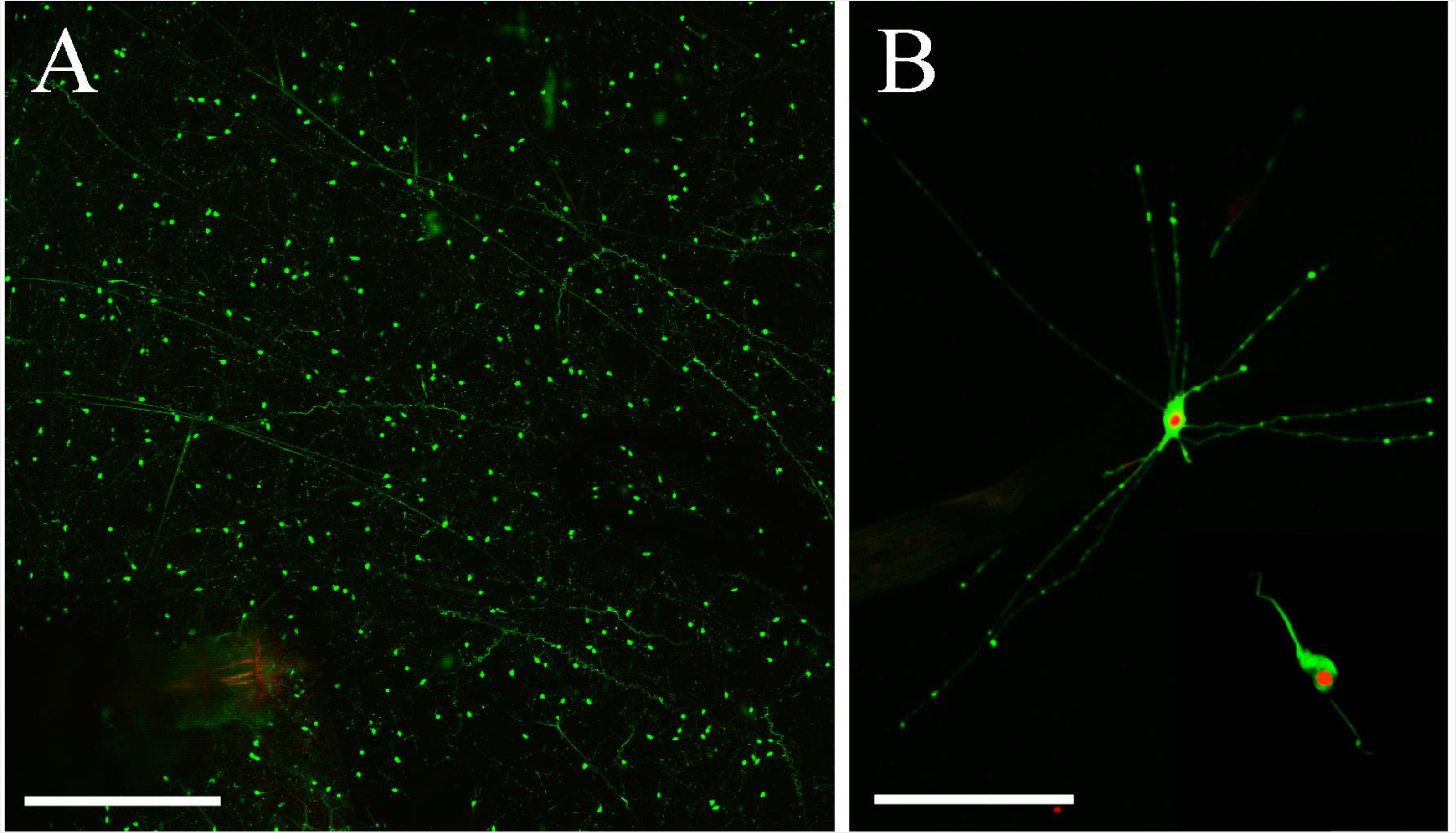
Mesogleal neurons. **A**: Many neurons stained by tubulin antibody (green) are spread over the entire mesoglea. **B**: All presumed neuronal somata had similar sizes but can be separated into different types based on the pattern of their processes. The most numerous type is represented by star-like cells with thin radial processes of similar length projecting in all directions. The second type is bipolar neurons with processes on opposite poles of the cell. Tubulin IR (green), SYTOX (red). Scale bars: A - 300 μm, B - 40 μm.

First, and the most numerous type was represented by cells with a thin processes of similar length projecting in all directions, which gave them a star-like shape (Fig. 19B). The second type of cells was represented by bipolar neurons with two processes on opposite sides (Fig. 19B). The third type included cells with long thin processes crossing the large areas of the mesogleal space (Fig. 20D, E, F).

**Figure 20.**
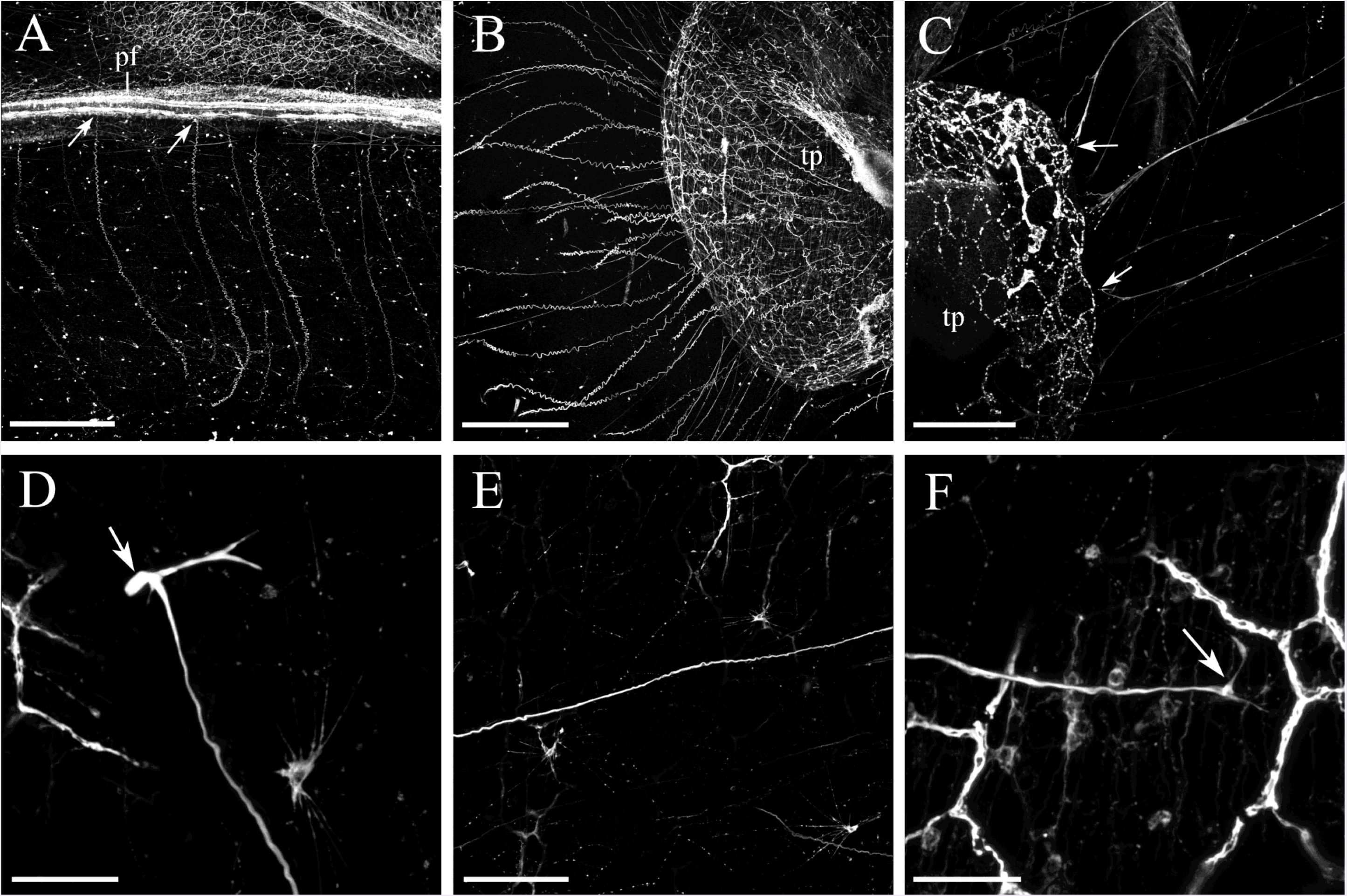
Mesoglial neuronal-like processes stained by tubulin antibody. **A**: Some of crossed large mesogleal regions and attached to the body surface structures including polar fields area (arrows) or the tentacle pockets (**B, C**). **D, E, F**: A single cell body (arrow) located on one side of the animal near the body surface (D) gives rise to a single process that crosses a large area of the mesogleal space (E) and terminates in the subepithelial network layer (arrow) almost on the other side of the body (F). Abbreviations: *pf* - polar field; *tp* - tentacle pocket. Scale bars: A, B - 300 μm, C - 150 μm, D - 25 μm, E - 75 μm, F - 25 μm.

Often, the neuronal-like processes in mesoglea were attached to the surface structures such as the aboral organ and polar fields, the mouth area and regions between the comb rows (Fig. 20) as well as tentacle pockets (Fig. 20B, C). The terminals always branched at the very end significantly increasing the contact area (Fig. 20C and 21A).

**Figure 21.**
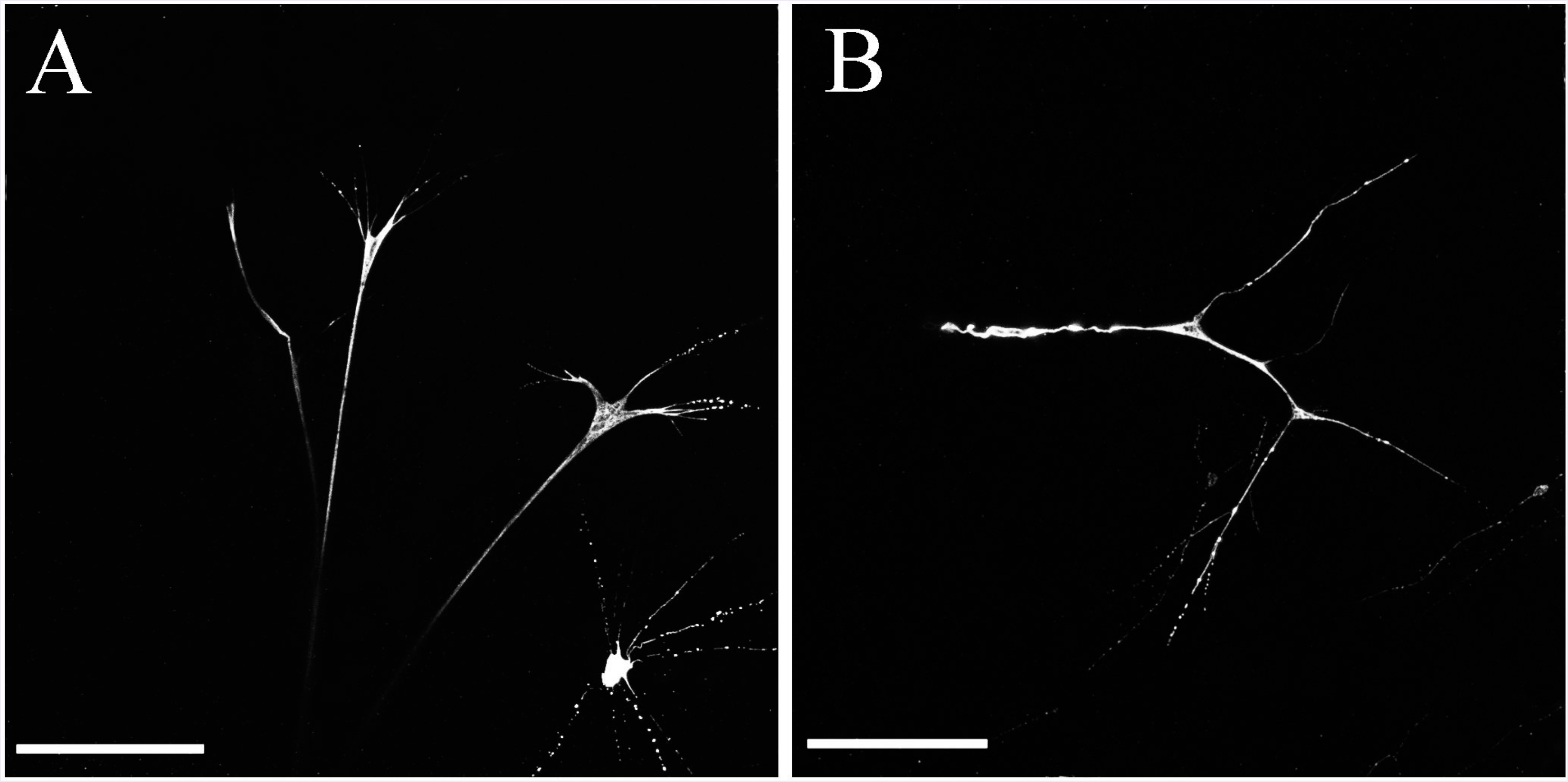
Mesoglial tubulin IR processes at a higher resolution near a target surface. (**A**) and in the middle of the mesoglea without any recognizable target (**B**). Scale bars: A - 75 μm, B - 50 μm.

We detected connections of mesogleal neuronal processes both to the surface polygonal neural network (Fig. 22A) and to the subepithelial nerve net in the tentacle pockets (Fig. 22B). Some of the neural-like terminals, however, ended shortly before making visible contact with the network (Fig. 22B) and the branching of the filaments occurred in the center of the mesoglea (Fig. 21B and 18C).

**Figure 22.**
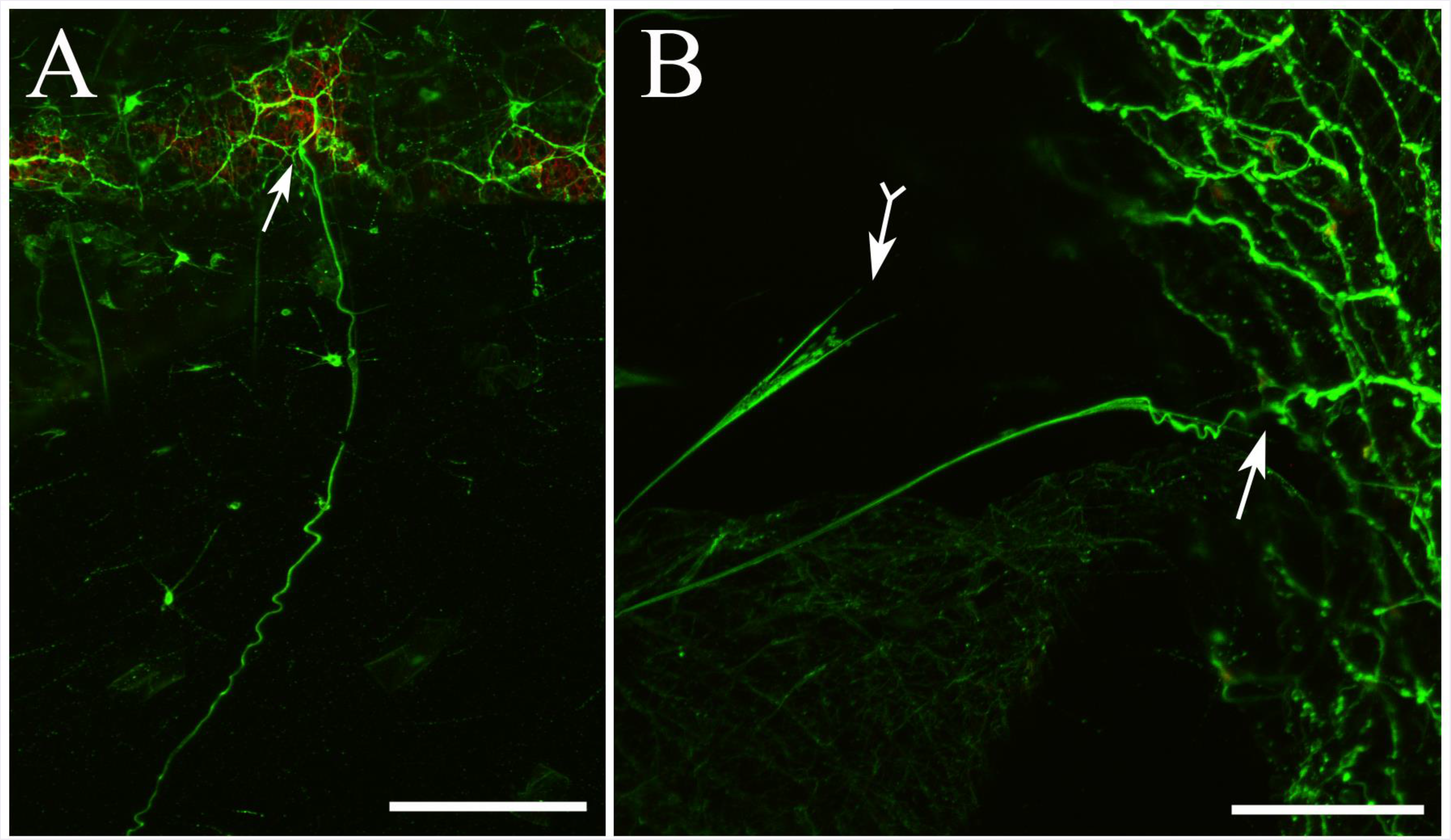
High resolution imaging shows a direct connection (arrows) between mesogleal neuronal processes and the surface polygonal neural net (**A**) and the subepithelial net in the tentacle pockets (**B**). Many of the processes, however, terminate shortly before making a direct visible contact with the polygonal network (arrow with tail). Scale bars: A - 100 μm, B - 50 μm.

In selected SEM experiments, the *Pleurobrachia* body was intentionally broke open after SEM drying process to get access to the inside organs. Cells and their processes (Figs. 23, 24) were preserved very well, while the mesogleal gelatinous tissue was washed away during the procedure. For example, there were many processes that at the point of contact with the pharynx surface had a different morphology than the muscle fibers (Fig. 23) and looked more like tubulin IR neurites described above (Fig. 24A, B). Unlike muscle fibers, which were firmly attached to the surface of the pharynx (Fig. 23C, D), these terminals produced several very thin and long branches that were barely connecting to the surface and were not capable of providing strong mechanical binding (Fig. 24C, D). These processes could be a part of the neural-like integrative system.

**Figure 23.**
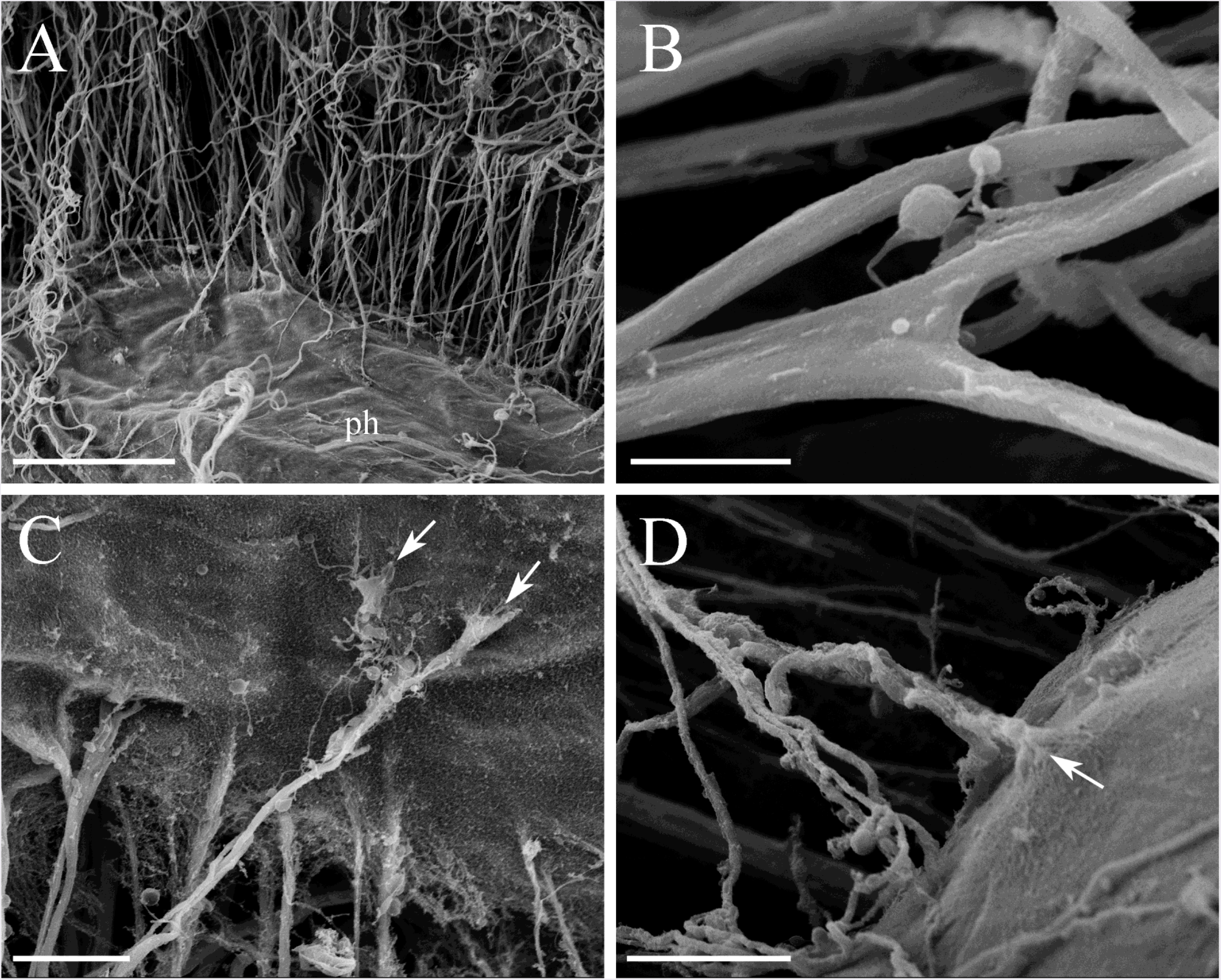
SEM of the pharynx muscles. **A**: A thick-walled pharynx (*ph*) with a well-preserved muscle distribution after a drying process. **B**: The muscles have variable thickness, and on a rare occasion some muscle fibers bifurcate into two branches. **C, D**: The muscle fibers are attached to the surface of the pharynx. Note that there is always some widening and branching at the point of contact to increase the binding surface (arrows). Scale bars: A - 100 μm, B - 10 μm, C, D - 20 μm.

**Figure 24.**
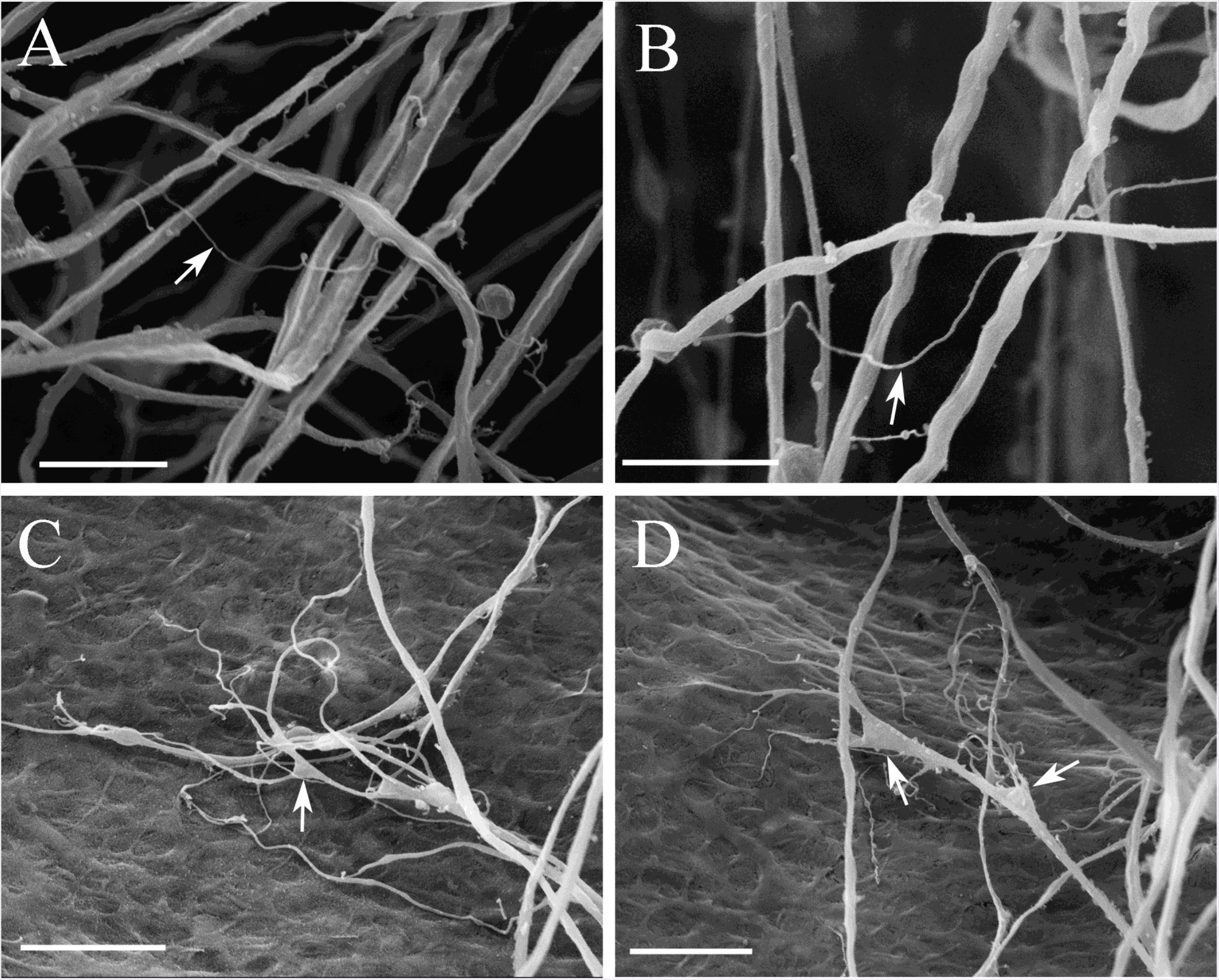
SEM of mesogleal neuronal-like processes among muscle fibers attached to the pharynx. **A, B**: In addition to thick muscle fibers, there are many very thin processes (arrows), which might be neuronal processes. **C, D**: At the point of contact with the pharynx surface, many neuronal-like processes have a morphology similar to the tubulin-labeled terminals described earlier. Unlike muscle cells, these processes produce several very thin and long branches (arrows) that are nor embedded into the surface and do not establish secure binding characteristic for muscles. These elements are presumably a part of the mesogleal neural system. Scale bars: A, B - 10 μm, C, D - 20 μm.

Using SEM, we detected a few mesogleal neuron-like cells associated with muscles (Fig. 25). The largest of these cells were 5-7 μm in diameter and had long thin processes (Fig. 25B, E, F). Some of the cells were multipolar, while others were bipolar with extensively branching processes (Fig. 25E, F). The mesoglial cells were found not only in the space among the muscle fibers, but also attached to the surface of the pharynx (Fig. 25C).

**Figure 25.**
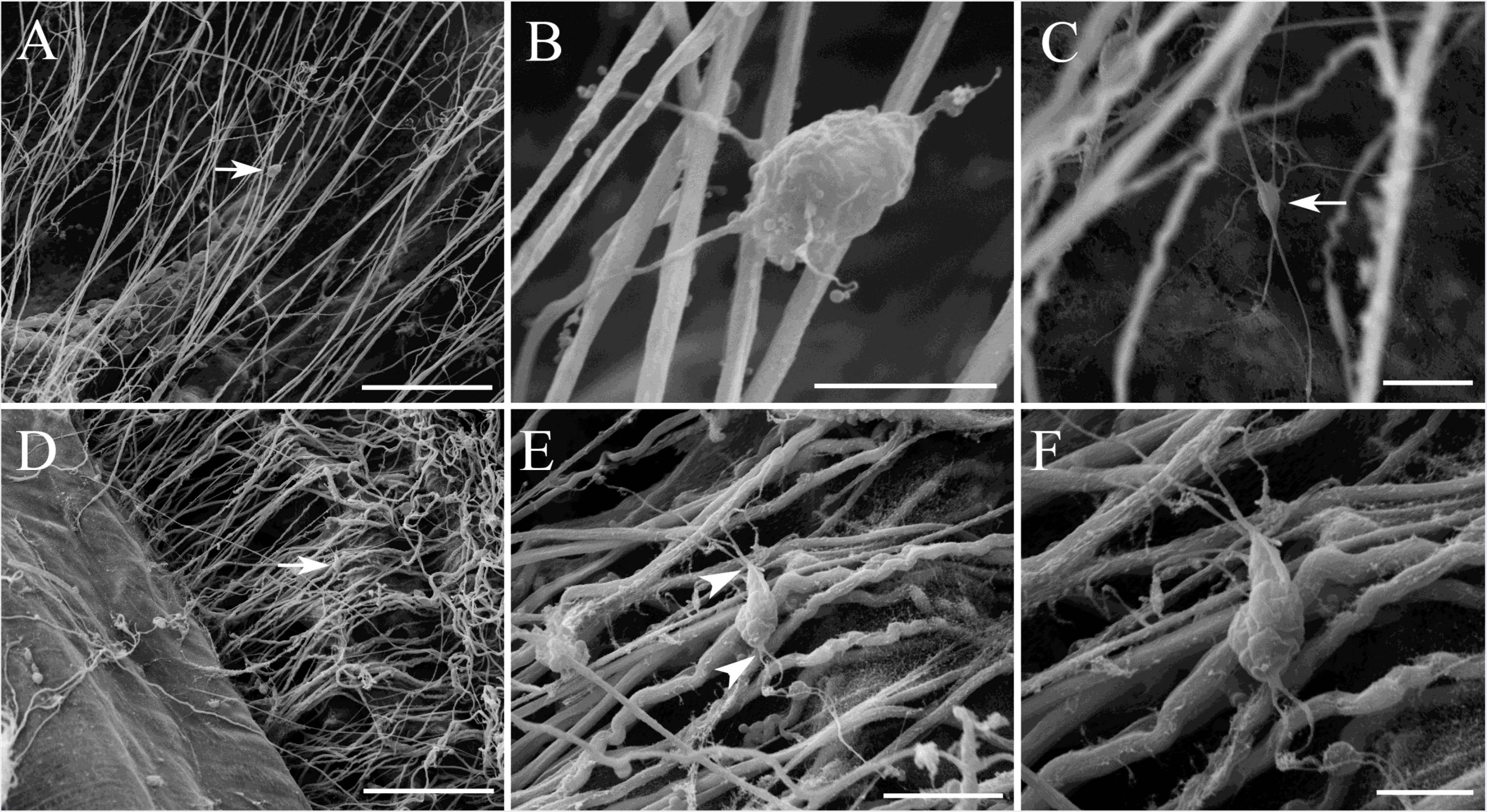
SEM of mesogleal neurons among muscles in the pharynx area. **A, B**: A large multipolar neuron (arrow) remains attached to the muscle fibers; low and high magnification. **C**: A neuron (arrow) on the surface of the pharynx with very long thin processes extending in different directions. **D, E, F**: A bipolar mesoglial neuron (arrow) among the muscle fibers (arrowheads) with extensively branching processes at each end; shown at different magnifications. Scale bars: A - 100 μm, B, C - 10 μm, D - 100 μm, E - 20 μm, F - 10 μm.

### 3.4 Identification of muscular and neural elements in comb rows

Swim cilia in ctenophores are one of the longest cilia in the animal kingdom with eight comb rows mediating complex locomotion of *Pleurobrachia*. SEM confirms that the swim cilia are fused together into individual plates - combs or ctenes (Fig. 26A). Each comb row is connected to the aboral organ (the major gravity sensor and integrative center (Tamm, 1982; Tamm, 2014)) by highly specialized ciliated furrows (Figs. 1A, B; 2A, C, D and 27A,B). Two neighboring or “sister” furrows fused near the aboral organ (Figs. 3A, 27A). At the earlier stages of development, the animals have only four ciliated furrows and comb rows, but on the 7-10th day, each furrow and each comb row begin to split into two “sister” structures, making eight homologous ciliated rows in adults (Norekian and Moroz, 2015).

**Figure 26.**
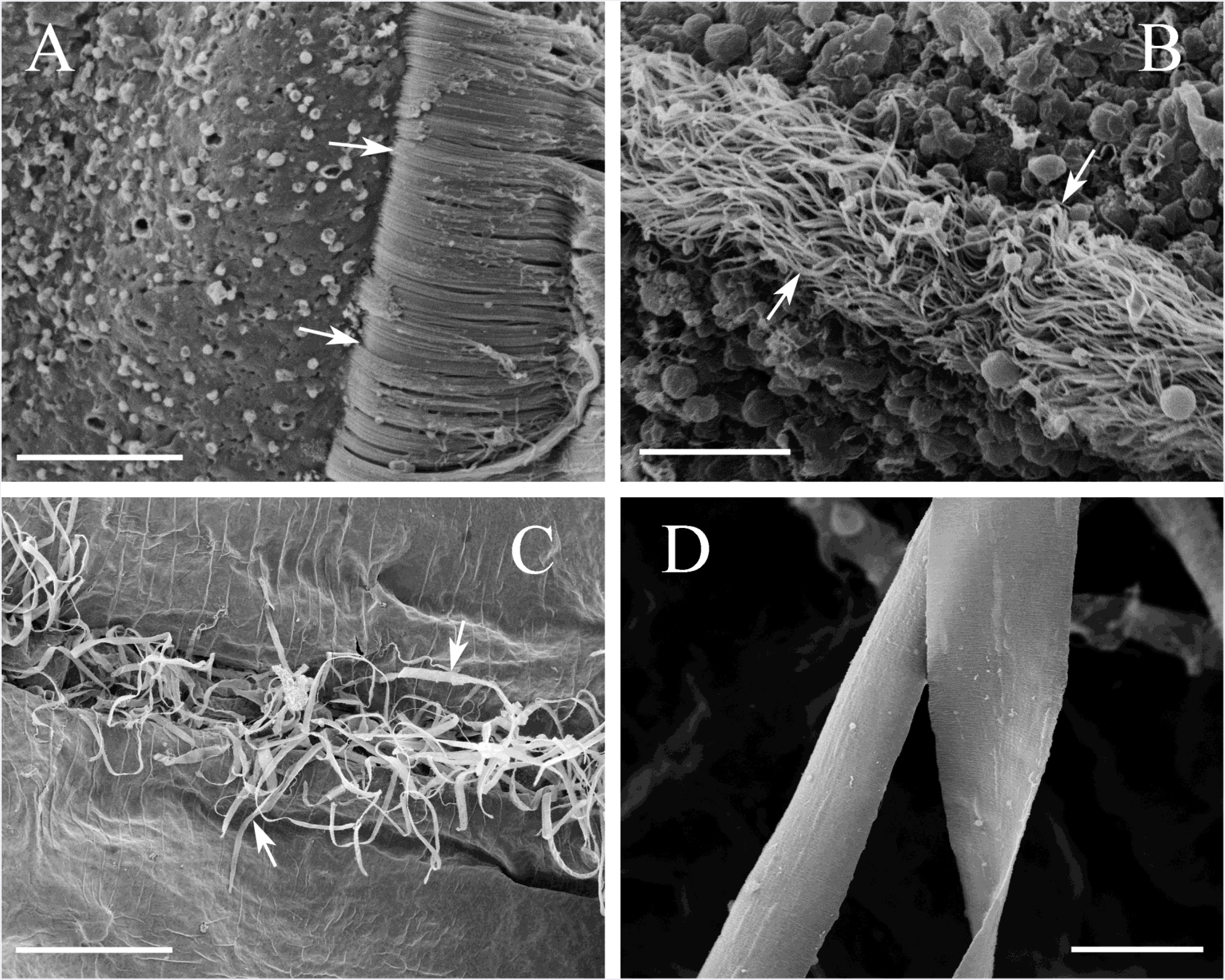
SEM of the combs and ciliated furrows. **A**: Comb plates represent a group of long swim cilia fused together (arrows). **B**: Ciliated furrow contains a narrow band of densely packed thin cilia (arrows). **C**: Each comb row had numerous flat and wide muscle cells (arrows) attached from the mesoglea side. These are the widest muscle cells in *Pleurobrachia* (D). Scale bars: A - 20 μm, B - 10 μm, C - 400 μm, D - 20 μm.

**Figure 27.**
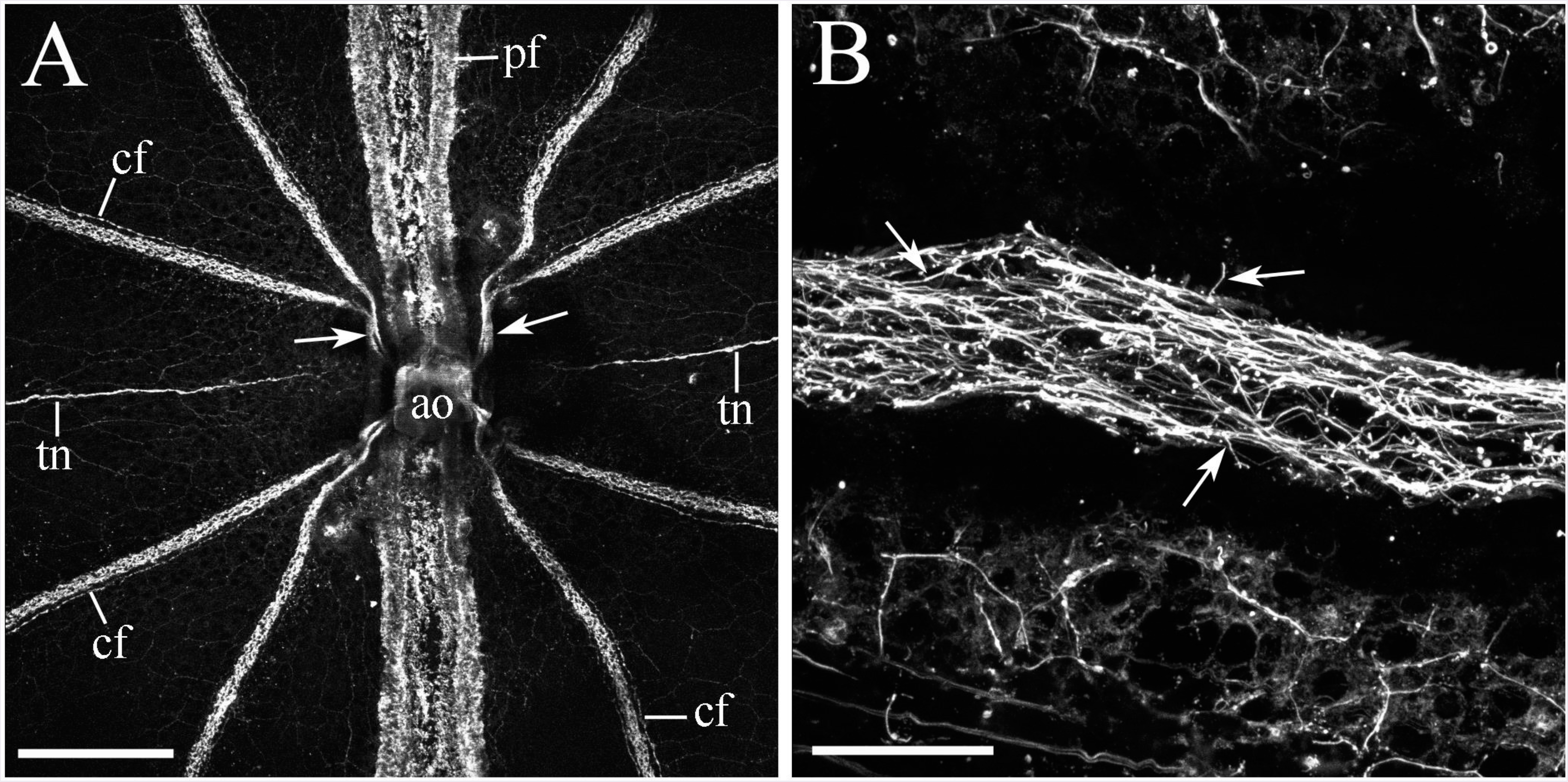
Ciliated furrows stained by tubulin antibody. **A**: The eight ciliated furrows connect their respective comb rows with the aboral organ. Note that two “sister” furrows (which were derived from one band in their early development) are merged close to the aboral organ (arrows). **B**: The ciliated furrows have a high density of motile cilia (arrows), which are also labeled by the tubulin antibody, but not phalloidin. Abbreviations: *ao* – the aboral organ; *pf* - polar field; *cf* - ciliated furrow; *tn* - tentacular nerve. Scale bars: A - 250 μm, B - 30 μm.

The ciliated furrows were densely covered with cilia (Figs. 26B, 27B). Only tubulin antibody (but not phalloidin) did label the cilia in the furrows and ctenes. Tubulin IR also revealed the polygonal neural net between individual comb plates (Fig. 29B).

Phalloidin labeled three distinct structures in the comb rows: (i) the modified muscle-like (non-contractive) fibers connecting neighboring comb plates (Fig. 28B, C, and 29A, B), (ii)-the basal cushion or polster at the base of each comb plate (Fig. 28B, C and 29A), (iii) and true muscles attached to the comb rows (Fig. 28D and 29C, D). The muscle-like fibers connecting the plates have many branches or extension within the same orientation (Fig. 28B, C and 29A). Most of them in the center mediated a direct mechanical connection between plates (Fig. 28B, C and 29A).

**Figure 28.**
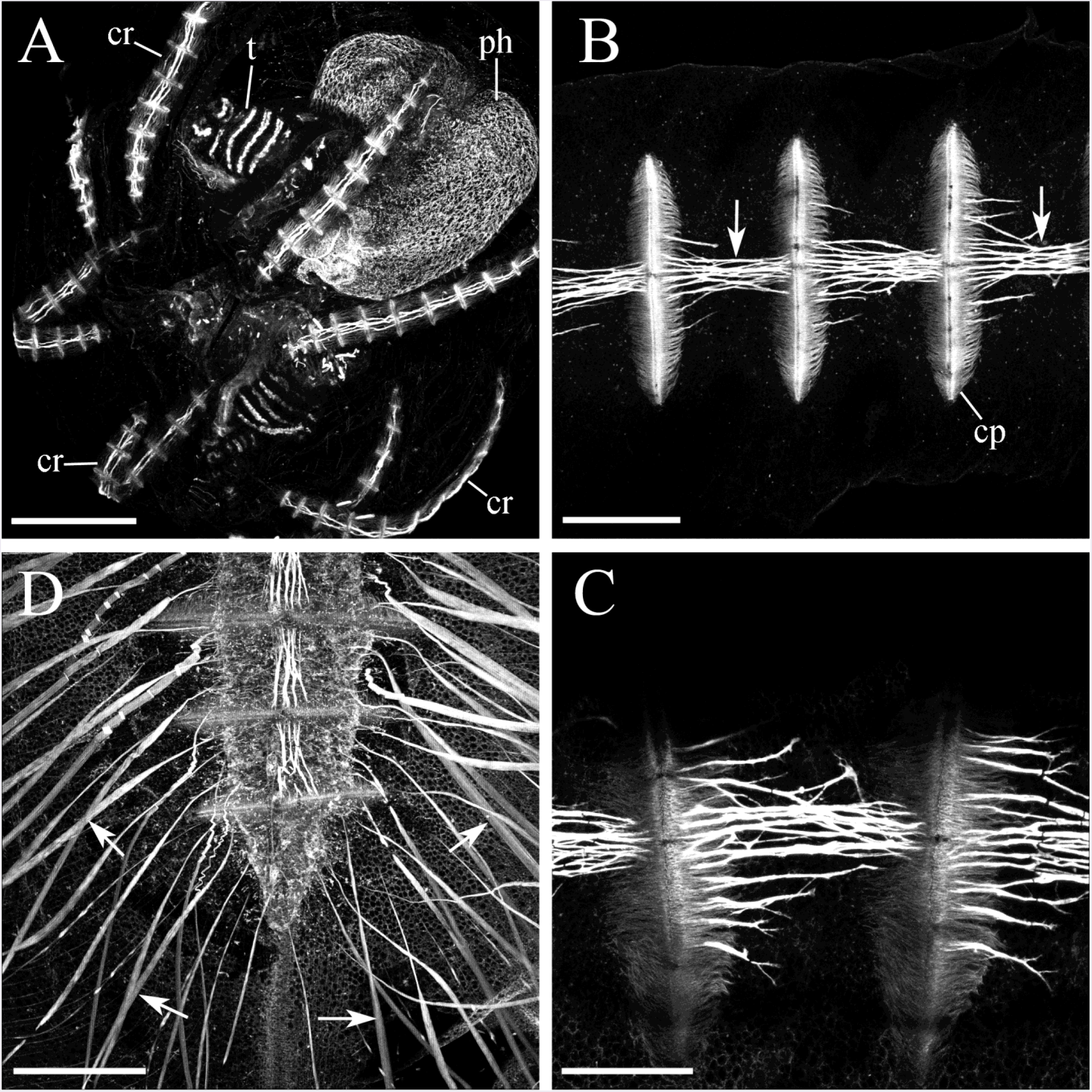
Phalloidin stains two main cell types associated with the comb rows. **A**: The entire small *Pleurobrachia* is stained with phalloidin. Comb rows are visible in addition to tentacles and pharynx that is completely outlined by phalloidin-stained cilia. **B**: A group of muscle-like elements that connect neighboring comb plates (their basal cushions are also labeled with phalloidin). **C**: These muscle-like non-contractive fibers often form branches within the same orientation along the comb row. **D**: The array of very long individual flat muscles (arrows) attached to comb rows (center). Abbreviations: *cr* - comb row; *cp* - comb plate; *t* - tentacle; *ph* - pharynx. Scale bars: A - 300 μm, B - 250 μm, C - 150 μm, D - 300 μm.

**Figure 29.**
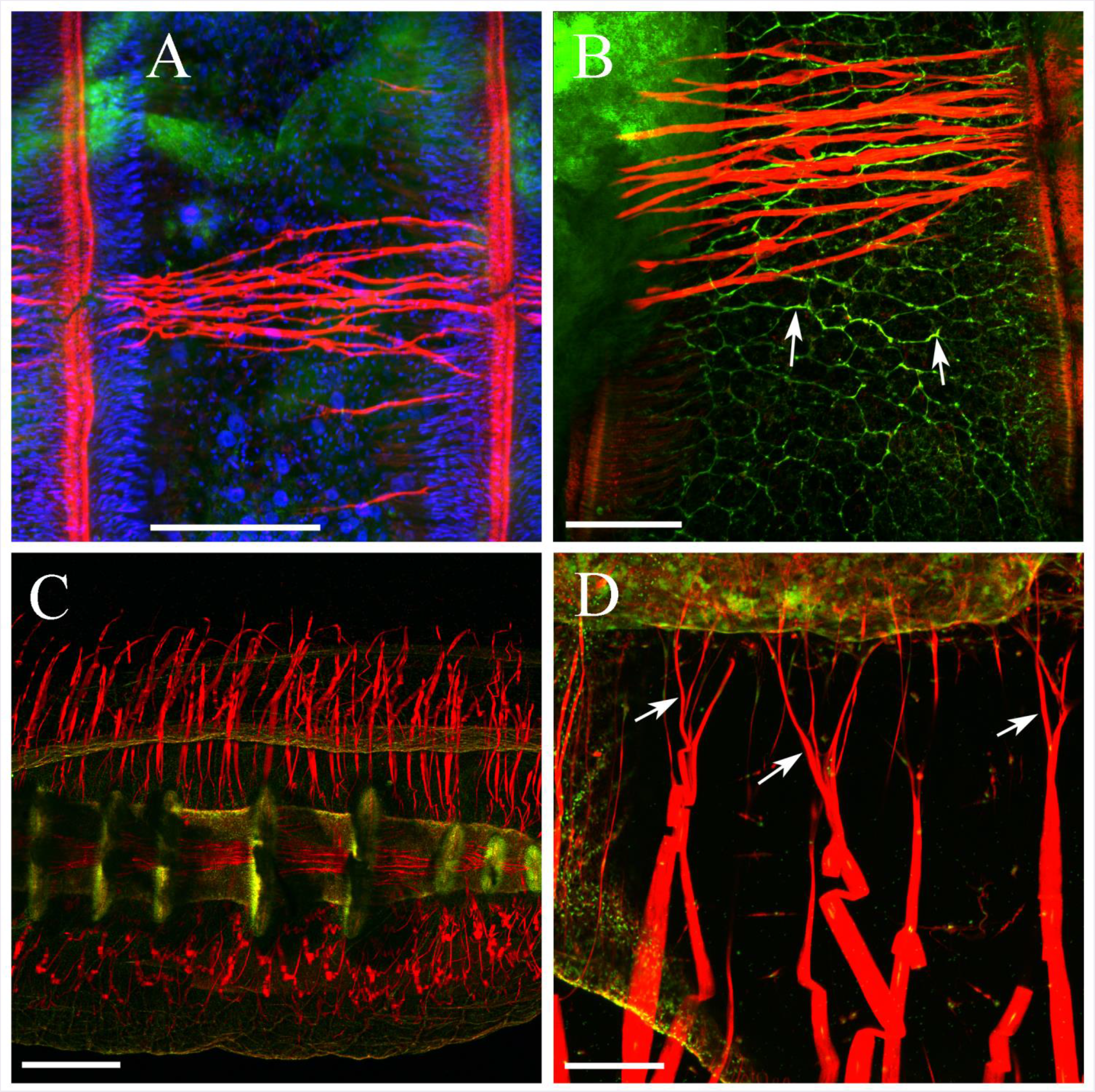
**Muscles and muscle-like elements in combs** (red -phalloidin staining). **A**: Most of the musclelike non-contracting fibers are long enough to establish connections between comb plates. However, DAPI (blue) labeling did not reveal nuclei in these phalloidin-labeled fibers (in contrast to the base of the comb plate). **B**: Tubulin IR (green) revealed the polygonal neural net (arrows) inside and between individual comb plates. **C**: Long flat muscles are attached to each comb row, from the aboral end to the oral side. **D**: Each flat muscle cell is 20-30 μm wide and about 2-3 mm long (for 1-2 cm large animals) with branching at the contact sites with the comb row (arrows). SYTOX green labels several nuclei in each muscle cells. Scale bars: A - 50 μm, B - 50 μm, C - 500 μm, D - 100 μm.

Hundreds of very long individual muscle cells were attached to both sides of comb rows and extended deep into the mesoglea (Figs. 28D, 29C, D, and SEM in Fig. 26C, D). Each muscle cell was 20-30 μm wide with branches close to the cilia row (Figs. 26D and 29D). These numerous muscles were evenly distributed through the entire length of each comb and could retract the comb rows as a defensive withdrawal response in *Pleurobrachia*.

### 3.5 Organization of the digestive system: muscles, cilia, and innervation

*Pleurobrachia* is an active predator frequently preying on different copepods: sometimes two or even three whole copepods could be seen inside the pharynx (Fig. 31C). Phalloidin clearly labeled muscles in copepods.

The mouth in *Pleurobrachia* is slightly elongated and covered with hundreds of short cilia (Fig. 30A, B), which facilitate movements of food particles. The oral complex continues into the ectodermal stomodaeum or pharynx (Fig. 1) with folded walls for efficient digestion (Fig. 30C). The pharynx has many muscles, attached to it from its mesogleal side (Fig. 30C, D). The pharynx opens into a short stomach chamber (Fig. 30D, E). These two digestive compartments are separated by a narrow sphincter, which isolates the pharynx from the canal system. The sphincter area is densely covered by long cilia (Fig. 30E). The thin-walled stomach extends into the aboral canal that then continues to the aboral complex, where it branches into two shorter canals with two muscular anal pores used for excretion (Fig. 2A and 30F).

**Figure 30.**
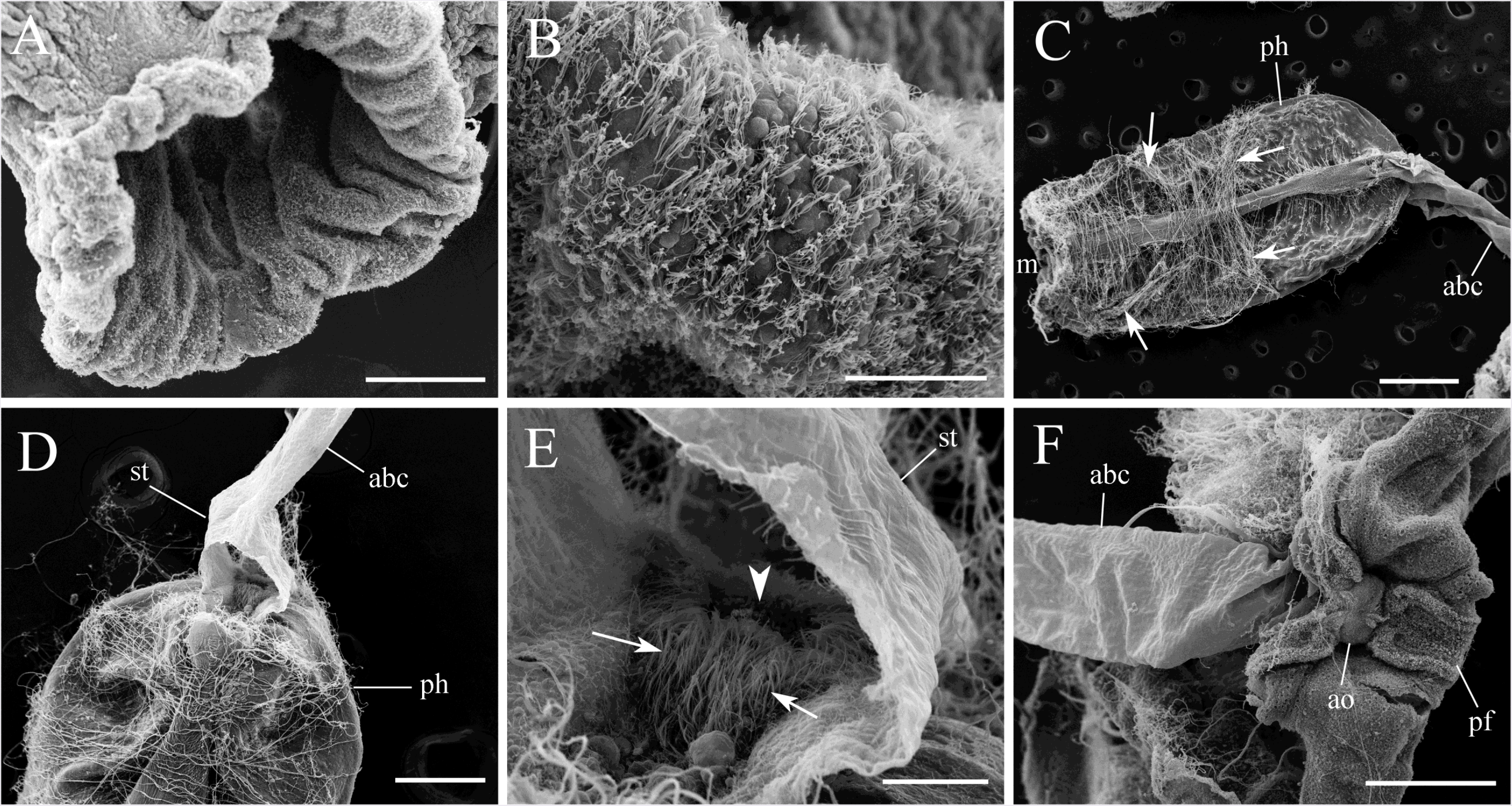
SEM of the digestive system. **A**: Elongated mouth. **B**: The entire area inside the mouth is covered with short cilia. **C**: The pharynx is always well preserved during SEM drying process with thin muscle filaments attached to it (arrows). **D**: The pharynx continues into a short thin-walled stomach chamber, and then to the aboral canal. **E**: The pharynx and stomach are separated by a sphincter (arrowhead), which is densely covered with long cilia inside the stomach (arrows). **F**: The aboral canal and the aboral organ (see details in Fig. 31F). Abbreviations: *ao* - aboral organ; *abc* - aboral canal; *m* - mouth; *pf* - polar field; *ph* - pharynx; *st* - stomach. Scale bars: A - 200 μm, B - 40 μm, C - 500 μm, D - 200 μm, E - 50 μm, F - 200 μm.

**Figure 31.**
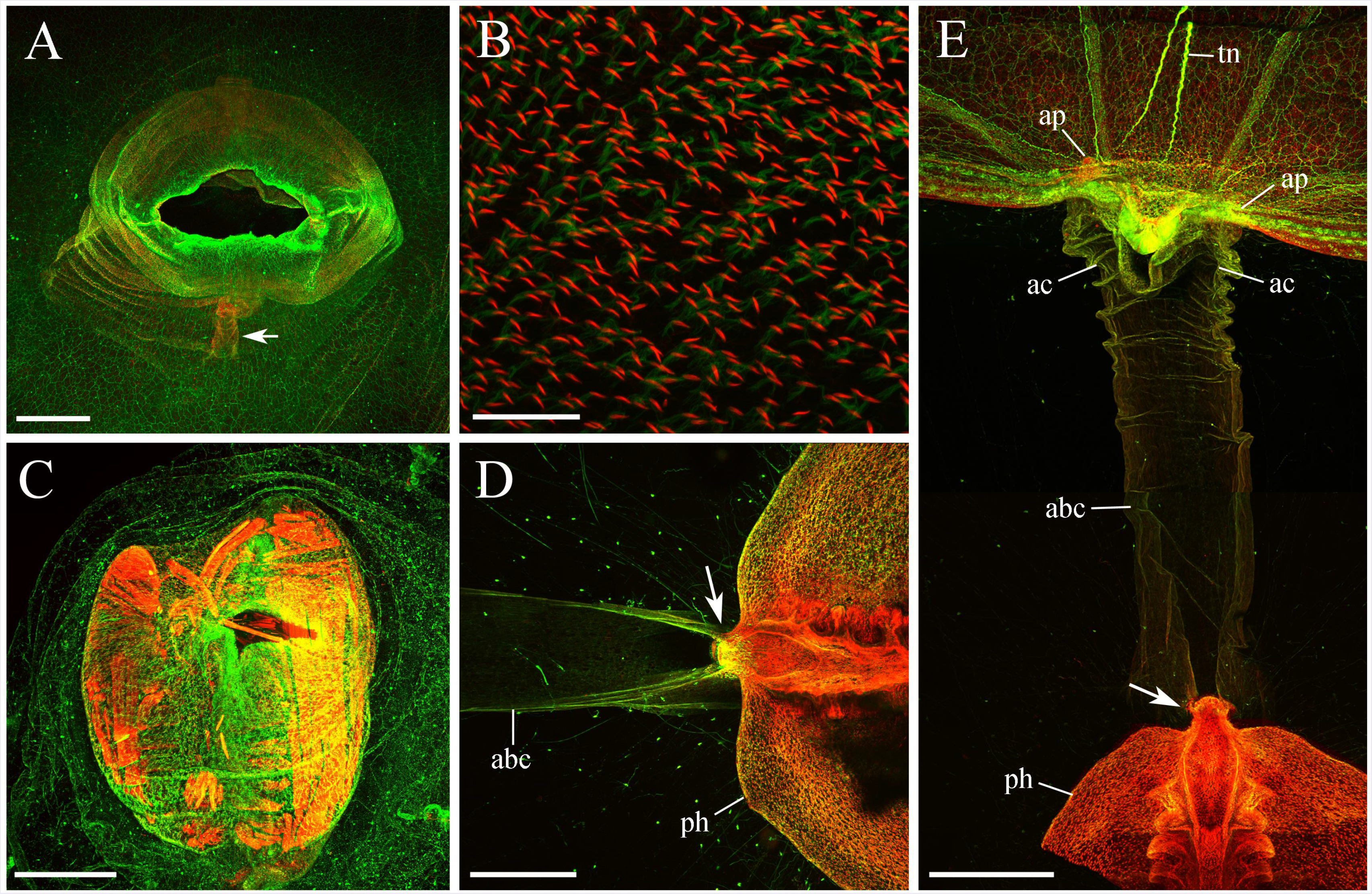
**Components of the digestive system** (double labeling by tubulin antibody (green) and phalloidin (red)). **A**: The neural net covers the entire mouth area. Two furrows are noticeable inside the mouth on the opposite sides (arrow). **B**: Internal surfaces of the mouth and pharynx are covered by short phalloidin-stained cilia. **C**: Two copepods inside the pharynx of a small *Pleurobrachia* with their muscle groups labeled by phalloidin. **D, E**: Pharynx connects to the short stomach chamber and aboral canal via a small constriction, or sphincter (arrow). The aboral canal continues to the aboral organ, where it branches into two short anal canals terminating into two anal pores on the opposite sides of the aboral organ. Abbreviations: *abc* - aboral canal; *ac* - anal canal; *ap* - anal pore; *ph* - pharynx; *tn* - tentacular nerve. Scale bars: A - 500 μm, B - 50 μm, C, D, E - 300 μm.

Neural plexus that covered the entire body of *Pleurobrachia* continues to the “lips” area (Fig. 31A) with two furrows inside the mouth. The mouth was outlined by numerous cilia, which brightly stained by phalloidin (Fig. 31B, E and 32B), but not tubulin antibody. A large group of muscles encircled the mouth (Fig. 32A, B) and mediated its closure. In contrast, fewer muscle filaments had radial orientation, presumably leading to the opening of the mouth (Fig. 32B).

**Figure 32.**
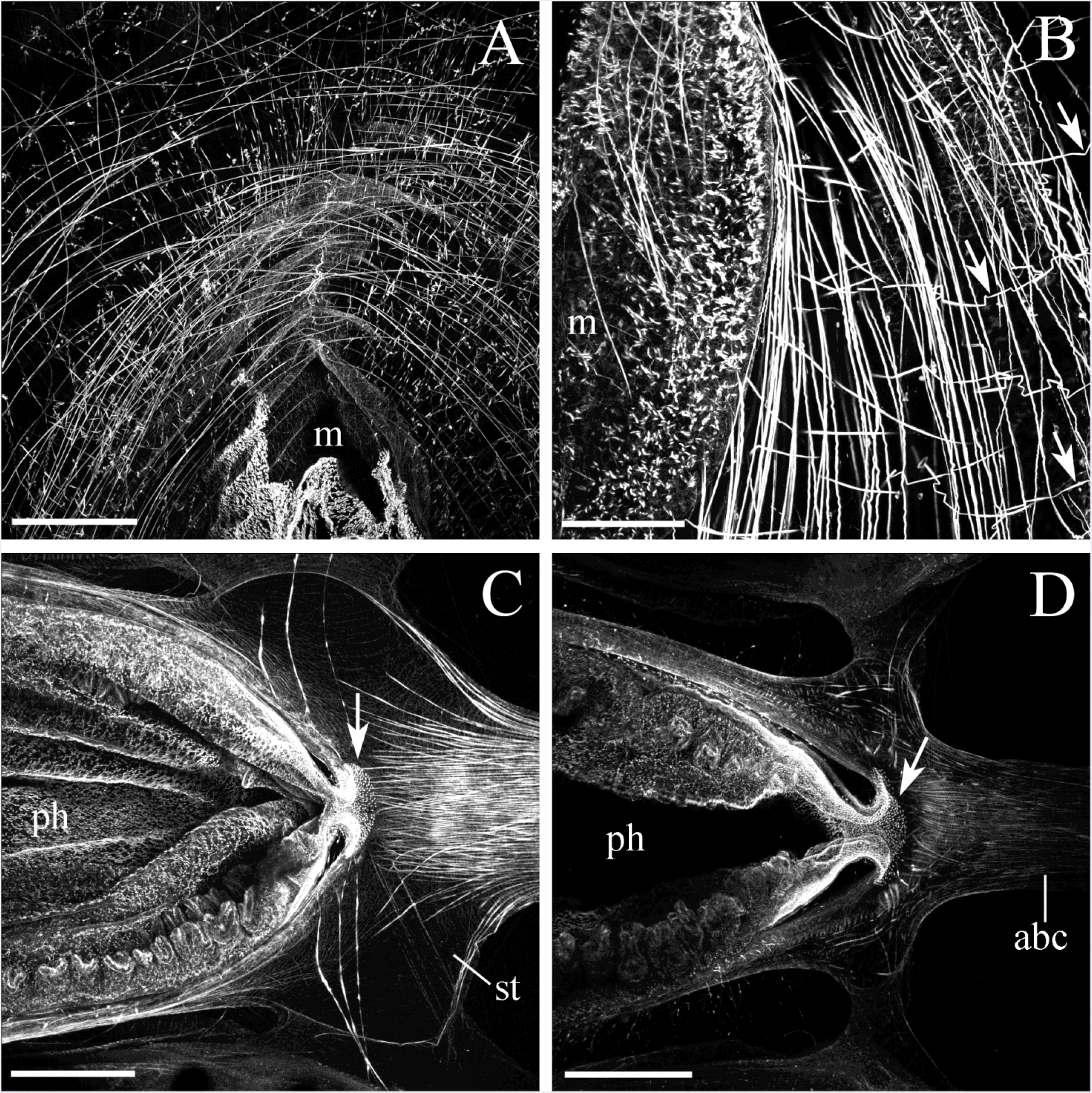
**Muscles and cilia of the digestive system** (phalloidin staining). **A, B**: Numerous circular and less abundant radial (arrows) muscles of the mouth. **C, D**: Muscular pharynx has thick folded walls densely outlined by short cilia. The sphincter (arrow) between the pharynx and stomach chamber is also covered by cilia. Note also thin muscles originating within the walls of the stomach and extended into the aboral canal. Abbreviations: *abc* - aboral canal; *m* - mouth; *ph* - pharynx; *st* - stomach. Scale bars: A - 300 μm, B - 150 μm, C, D - 300 μm.

Like the mouth, the elongated pharynx was covered with numerous short cilia (Fig. 31D, E and 32C, D). The pharynx was connected to the short stomach chamber via a small constriction, or sphincter, which could isolate the pharynx from the thin-walled stomach (Fig. 31D, E and 32C, D). It appears that this sphincter allows only partially digested food or small particles to go through the system, while large parts like chitinous remains of the copepods would remain in the pharynx and be ejected via the mouth. The sphincter was also covered with cilia stained by phalloidin (Fig. 32C, D) similarly to oral parts of the digestive system.

The short, thin-walled stomach had the opening into the whole gastrovascular canal system. The first part includes the aboral canal that extended toward the aboral organ (Fig. 31E). The wall of the aboral canal contained a layer of thin longitudinal muscles (Fig. 13A, 32C and 33D). The stomach chamber also connected to two short transverse canals, which turned into two tentacular and four interradial canals, running in radial directions through the mesoglea (Hernandez-Nicaise 1991). Each interradial canal branches into two adradial canals and reaching every comb rows subsequently forming the meridional canals (Fig. 34A, B).

**Figure 33.**
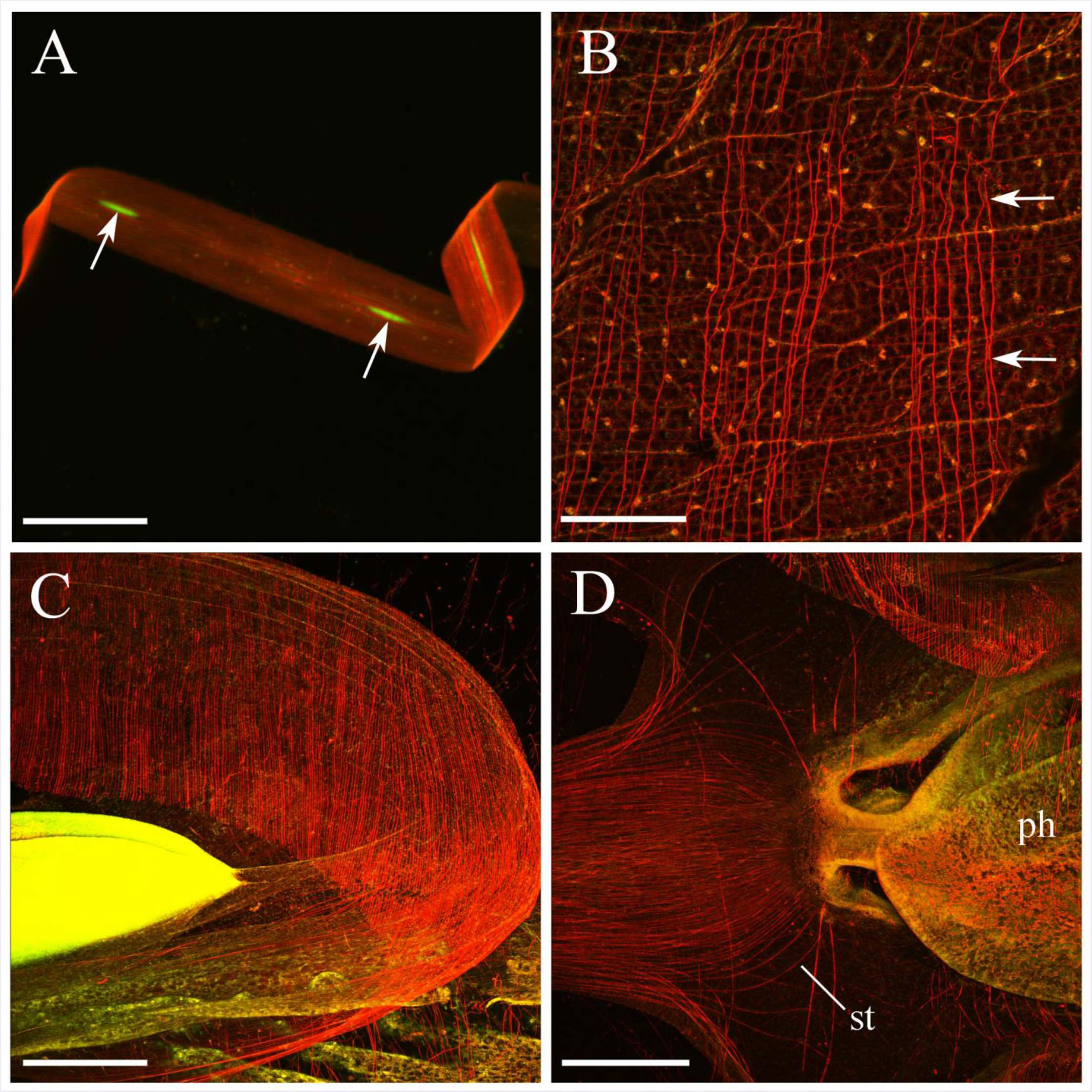
The diversity of muscle groups stained by phalloidin (red). **A**: Flat and wide muscle cell, which is attached to the comb row. Each cell has many nuclei (arrows) stained by SYTOX green over its entire length. **B**: External parietal muscles (arrows) in the outer body wall. **C**: The wall of the tentacle pocket with circular and longitudinal muscles. **D**: Muscles of the stomach and the aboral canal. Abbreviations: *ph* - pharynx; *st* - stomach. Scale bars: A - 40 μm, B - 150 μm, C, D - 300 μm.

**Figure 34.**
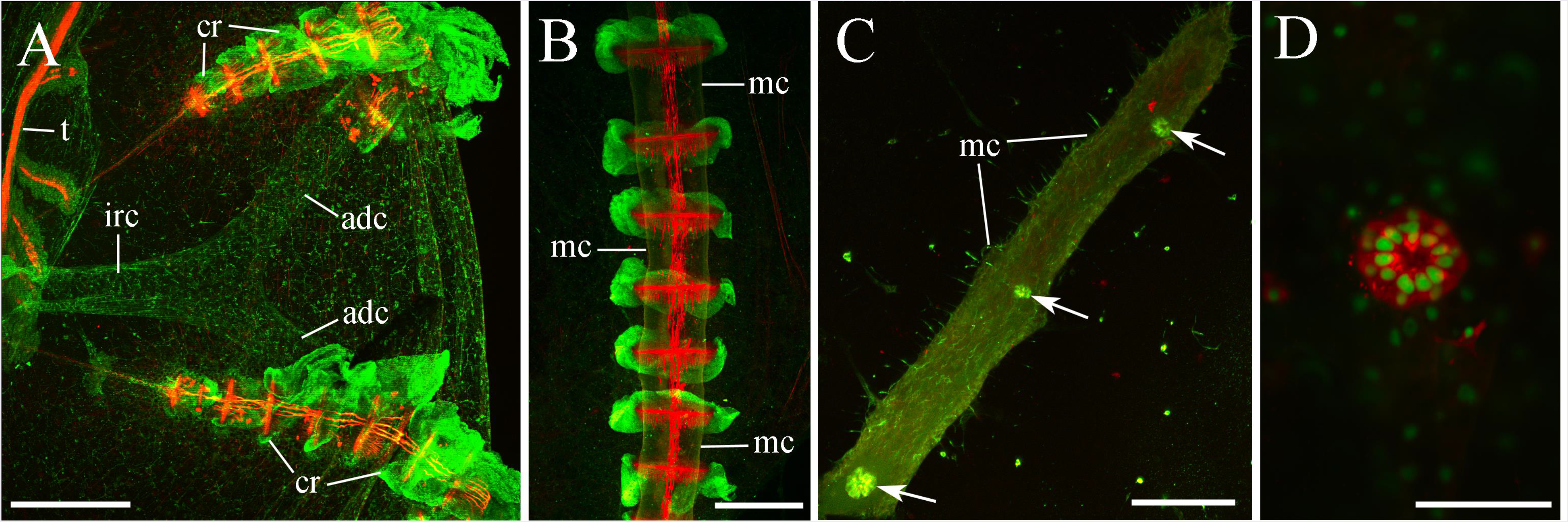
**Gastrovascular canals in *Pleurobrachia*** (tubulin IR is green, phalloidin is red). **A**: Interradial canal branches in two adradial canals, which subsequently open into meridional canals running under each comb row. **B**: Meridional canals are localized under each comb row. **C**: Meridional canal has a series of pore complexes (ciliated rosettes) along its length. **D**: Each pore consists of at least 10-12 cells forming a circle. SYTOX green stains the nuclei of these cells in green, while tubulin IR is red in this photo. Abbreviations: *irc* - interradial canal; *adc* - adradial canal; *mc* - meridional canal; *cr* - comb row; *t* - tentacle. Scale bars: A - 300 μm, B - 500 μm, C - 50 μm, D - 20 μm.

The interradial and adradial canals were lined up with cilia (Fig. 35C) facilitating the nutrient flow along the gastrovascular system. These cilia were specifically labeled by tubulin antibody (Fig. 35A, B), but not phalloidin.

**Figure 35.**
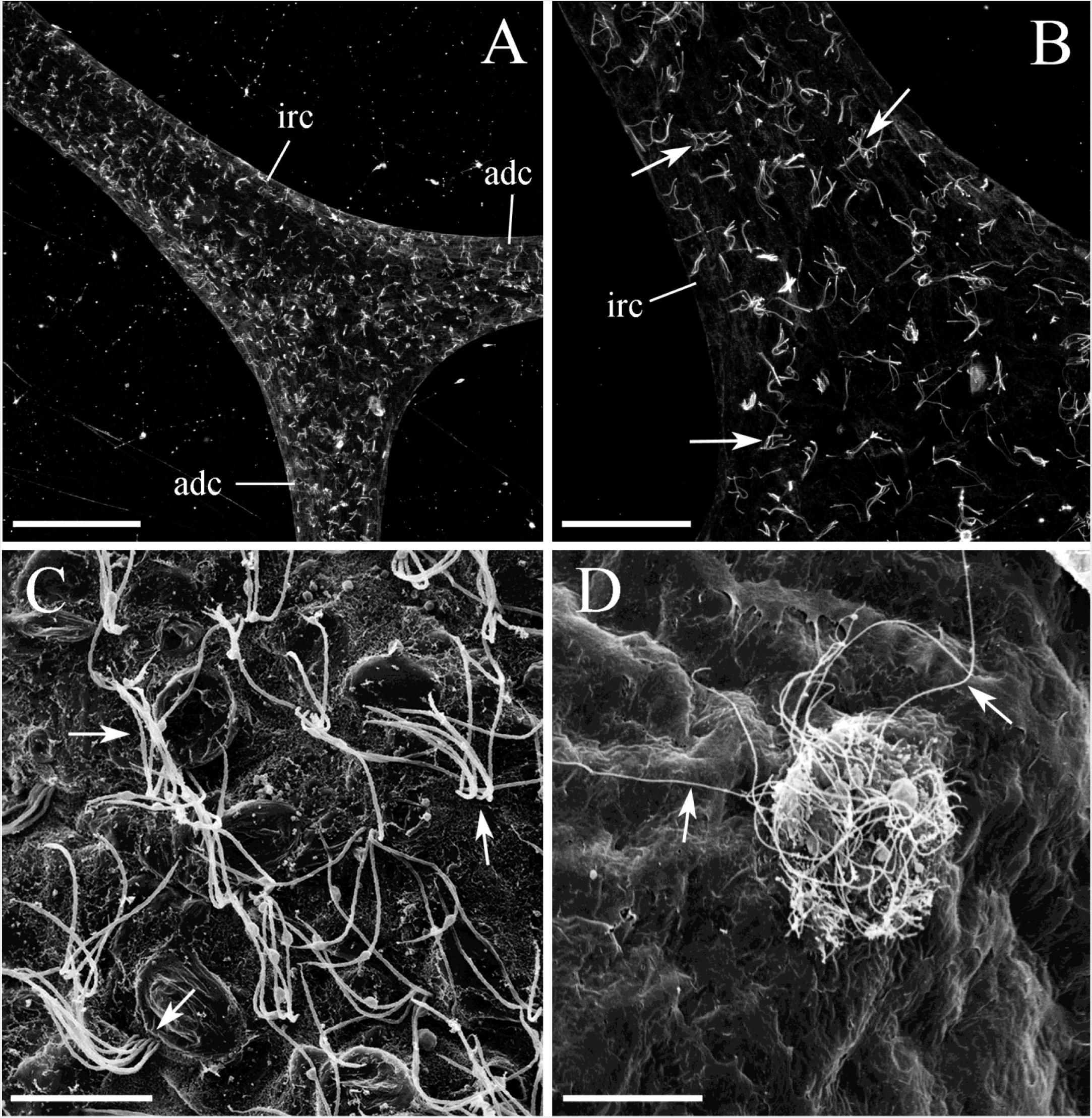
**A, B**: The interradial and adradial canals with cilia visualized by tubulin antibody staining (arrows). **C**: SEM image of cilia (arrows) outlining the inner surface of the meridional canals. **D**: SEM image of a single pore on the outside surface of a meridional canal facing the mesoglea. Note long thin cilia of the pore (arrows). Abbreviations: *irc* - interradial canal; *adc* - adradial canal. Scale bars: A - 150 μm, B - 50 μm, C - 10 μm, D - 20 μm.

Tubulin antibody also stained several individual pores located along each meridional canal (Fig. 34C), called ciliated rosettes (Hernandez-Nicaise 1991). These pores were found through the entire length of every canal under the comb rows facing the mesogleal area of the body away from the combs. SYTOX green staining, which was used to label the nuclei in the cells, revealed that each pore consisted of at least 10-12 cells forming a circle (Fig. 34D). Numerous long cilia attached to every single rosette from the mesoglea side (Fig. 35D). Some of the pores were completely closed, while others were wide open in our fixed preparations. These pores provided a connection between the gastrovascular canal system and mesogleal area of the body.

Using SYTOX green or DAPI for nuclei staining had an unusual side-effect in our studies providing a nice visualization of some anatomical structures. For example, SYTOX green in our preparations brightly stained parts of the reproductive system (Fig. 36). *Pleurobrachia* is a hermaphrodite, like most of the ctenophores, with bands of male and female gonads residing in the walls of the meridional canals: the ovary is located on one side and the testis on the other (Fig. 36B, C).

**Figure 36.**
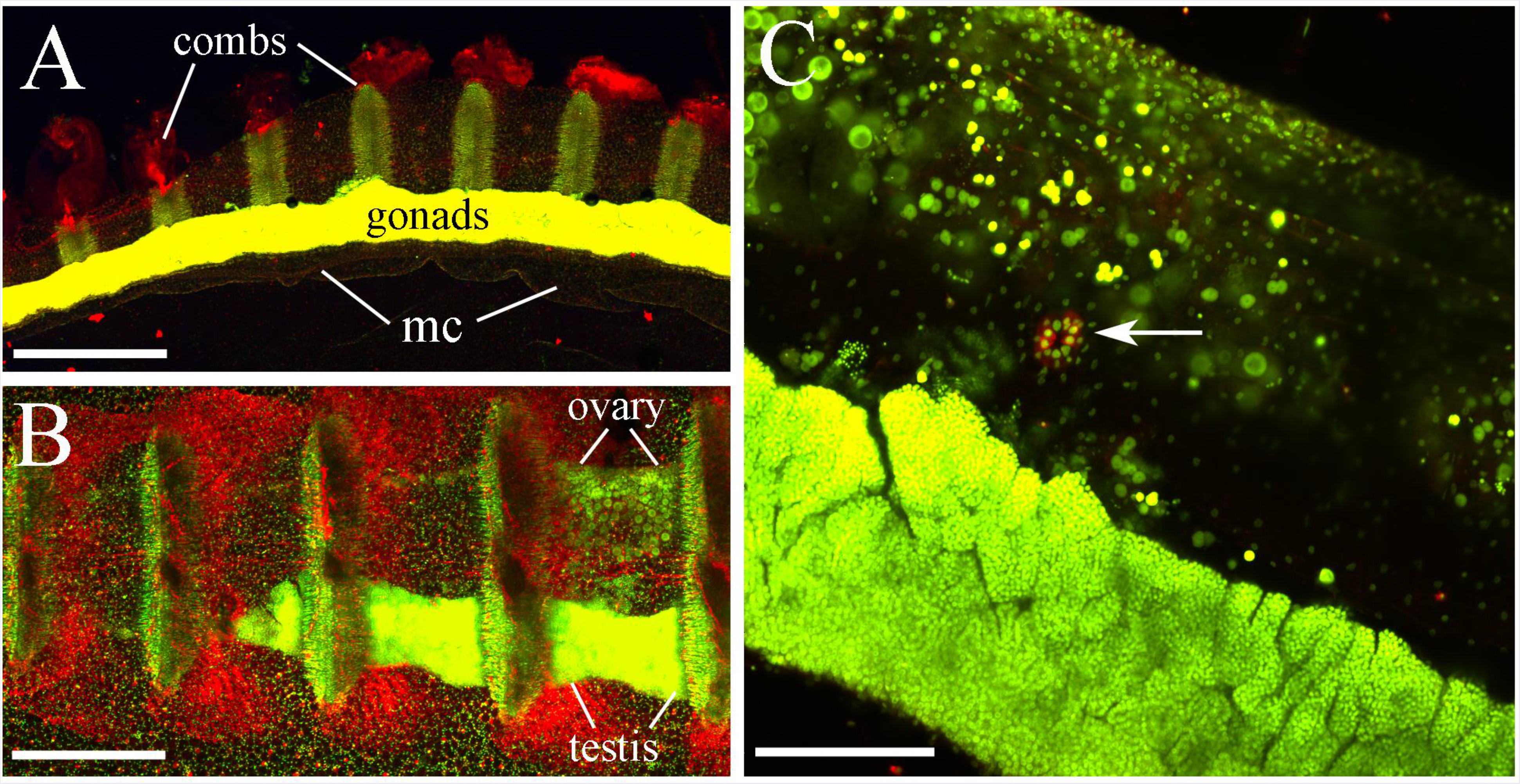
**Reproductive system revealed by SYTOX green staining** (shown in green-yellow color, while tubulin IR is red). **A**: Sagittal view of the comb row shows brightly stained gonads within the meridional canal (*mc*). **B**: Horizontal view of the comb row shows ovary and testis in two separate longitudinal bands of the meridional canal under the comb plates. **C**: Higher magnification of the meridional canal with a testis at the bottom and ovaries on the top. The arrow points to a pore. Scale bars: A - 500 μm, B - 250 μm, C - 50 μm.

### 3.6 The diversity of muscle systems in *Pleurobrachia*

Table 4 summarizes the information about more than a dozen muscle types in *Pleurobrachia*. In fact, the diversity of muscles and muscle type cells (frequently labeled by phalloidin) exceed diversity of neuronal or receptor subtypes (see Tables 1 and 2) and comparable with the diversity of ciliated cells (Table 3).

**Table 3.** Types of cilia in *Pleurobrachia*, and their markers.

|  | *Cilia type* | *Marker* | *Figures* |
| --- | --- | --- | --- |
| 1. | Cilia on neurons between strands of subepithelial network | Tubulin AB | Fig. 11 A, B, C |
| 2. | Receptors with a group of 2-7 cilia | Phalloidin | Fig. 11 F |
| 3. | Receptors with a long and curled cilium | Tubulin AB | Fig. 11 G |
| 4. | Receptors with a large and thick cilium | Phalloidin | Fig. 11 H |
| 5. | Receptors on the surface of the tentacles | Phalloidin | Fig. 25 C, D |
| 6. | Aboral organ dome cilia | only SEM | Fig. 2 A, B |
| 7. | Cilia on the edges of polar fields | Tubulin AB | Fig. 2 C, D; 8 B |
| 8. | Cilia of the ciliated furrows | Tubulin AB | Fig. 26 A, B; 29 B |
| 9. | Swim cilia of ctenes | Tubulin AB | Fig. 29 A; 34 A, B |
| 10. | Cilia inside the mouth and pharynx | Phalloidin | Fig. 30 A; 31 B, E |
| 11. | Cilia around the pharynx-stomach sphincter | Phalloidin | Fig. 30 E; 32 C, D |
| 12. | Cilia covering the gastrovascular canals | Tubulin AB | Fig. 35 A, B, C |
| 13. | Ciliated rosettes in the meridional canals | only SEM | Fig. 35 D |

**Table 4.** Muscle cell types and groups in *Pleurobrachia*.

|  | *Muscle group* | *Figures* |
| --- | --- | --- |
| 1. | Long muscles crossing the mesogleal region, some attached from inside to the body wall, pharynx and tentacle pockets | Fig. 16; 21 |
| 2. | Long parietal muscles in the body wall forming rectangular mesh | Fig. 33 B; 37 A, B |
| 3. | Muscles attached to the aboral organ and mediating its withdrawal | Fig. 6 C |
| 4. | Muscle-like cells related to four balances in the aboral organ | Fig. 5 B |
| 5. | Muscles attached to the polar fields and mediating their withdrawal | Fig. 6 D |
| 6. | Muscles of the main trunk of the tentacles | Fig. 25 A |
| 7. | Double strains of muscles in tentilla | Fig. 25 A |
| 8. | Muscles in the wall of a tentacle pocket | Fig. 33 C |
| 9. | Muscle-like non-contractile fibers connecting comb plates | Fig. 27 B, C; 28 A, B |
| 10. | Long, wide and flat muscles attached to comb rows | Fig. 27 D; 28 C, D; 29 C, D |
| 11. | Circular muscles around the mouth | Fig. 32 A, B |
| 12. | Radial muscles around the mouth | Fig. 32 B |
| 13. | Muscles inside the thick wall of a pharynx | Fig. 32 C, D |
| 14. | Thin longitudinal muscles in the walls of stomach and aboral canal | Fig. 32 C; 33 D; 37 C |
| 15. | Muscles of a sphincter between pharynx and stomach | Fig. 31 E; 32 C |

Muscle cells attached to the cilia rows were among the largest and, perhaps, the most powerful muscle groups in *Pleurobrachia* (Figs. 26C, D and 29D). These multinucleated cells were flat, 20-30 μm wide (Fig. 33A), and extend into the mesoglea. Although *Pleurobrachia* does not change the body shape, these flat muscles, as well as body wall muscles, could support the maintenance of the hydroskeletal and mediated local movements such as defense withdrawal of comb rows.

The body wall of the *Pleurobrachia* contained a number of external parietal muscles, which formed a loose rectangular network of fibers (Figs. 33B and 37A). Most of them were running in the oral-aboral direction, while some filaments were localized in the perpendicular direction (Fig. 37A). Near the aboral complex, the pattern of muscles was different. Muscle fibers were running in parallel to each ciliated furrow and then were ‘bent’ towards the neighboring ciliated furrow (Fig. 37B). Phalloidin revealed many muscles in the wall of each tentacle pocket (Fig. 33C). Contraction of these muscles might help to extend the tentacles. There were also many distinct groups of muscles in the wall of the stomach and circular muscles in the aboral canal (Fig. 33D and 37C).

**Figure 37.**
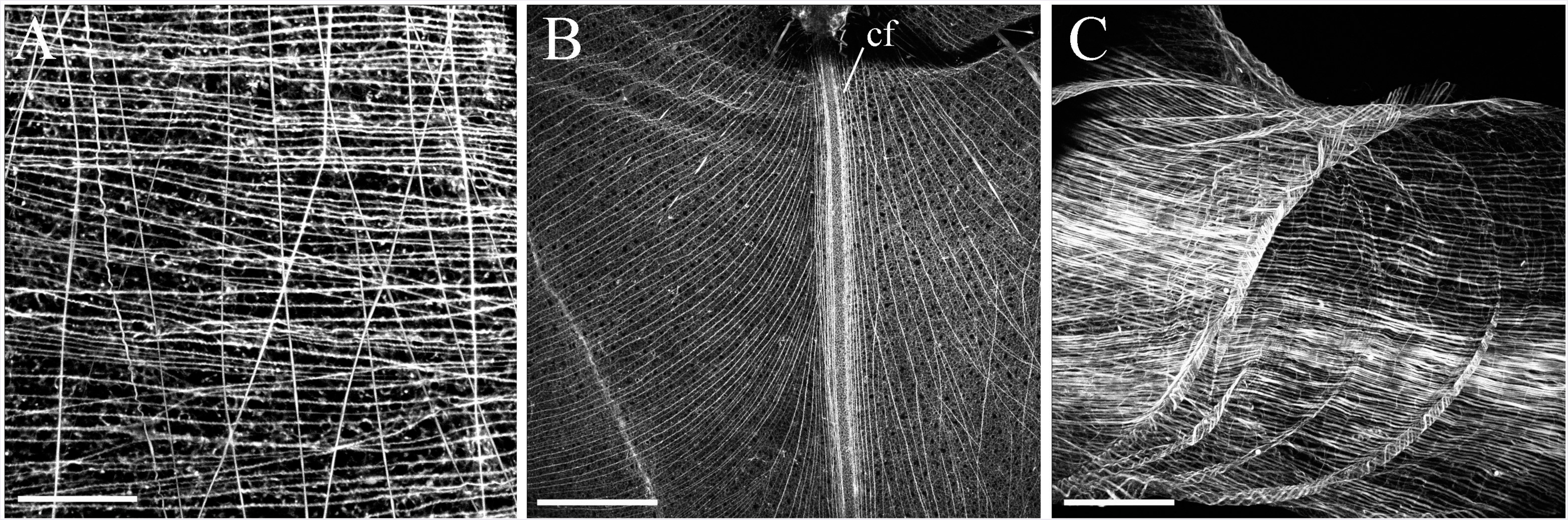
Elongated muscles in *Pleurobrachia* (phalloidin labeling). **A**: Thin external parietal muscles forming a loose rectangular mesh of fibers in the body wall. **B**: Closer to the aboral organ muscles are running in parallel to each ciliated furrow *(cf)* and then making a loop towards the neighboring ciliated furrow. **C**: The majority of muscles in the wall of the aboral canal are longitudinal with very few circular filaments. Scale bars: A - 70 μm, B - 300 μm, C - 120 μm.

We also identified a broad spectrum of muscle filaments crossing the mesogleal region and attached to the body wall, pharynx or tentacle pockets (Fig. 18).

There were also many distinct groups of muscles in the wall of the stomach and circular muscles in the aboral canal (Fig. 33D and 37C) and other parts of the digestive system as revealed by SEM microscopy (Fig. 23).

Deciphering tissue organization and a cell type diversity in the mesoglea was challenging. Numerous muscle fibers identified in the mesoglea, including the ones attached to the pharynx, were stained by phalloidin only and not labeled by tubulin antibody. They were usually thicker and different than neuronal filaments. In contrast, the mesoglea also contained many very thin neuronal filaments that were labeled only by tubulin antibody and were not stained by phalloidin.

However, a small fraction of the cells with long processes that were labeled by tubulin antibody also showed phalloidin staining in double-labeling experiments (Fig. 38A, B). In the freshly stained preparations (within two days of processing), when the concentration of phalloidin was at the highest end, and the antifade mounting medium was used, the proportion of filaments stained by both phalloidin and tubulin antibody could reach 20-30%. Even a few cells (but not all) with neuron-like morphology were labeled by phalloidin under these conditions. In contrast, the neurons within the subepidermal neural net were never labeled by phalloidin.

**Figure 38.**
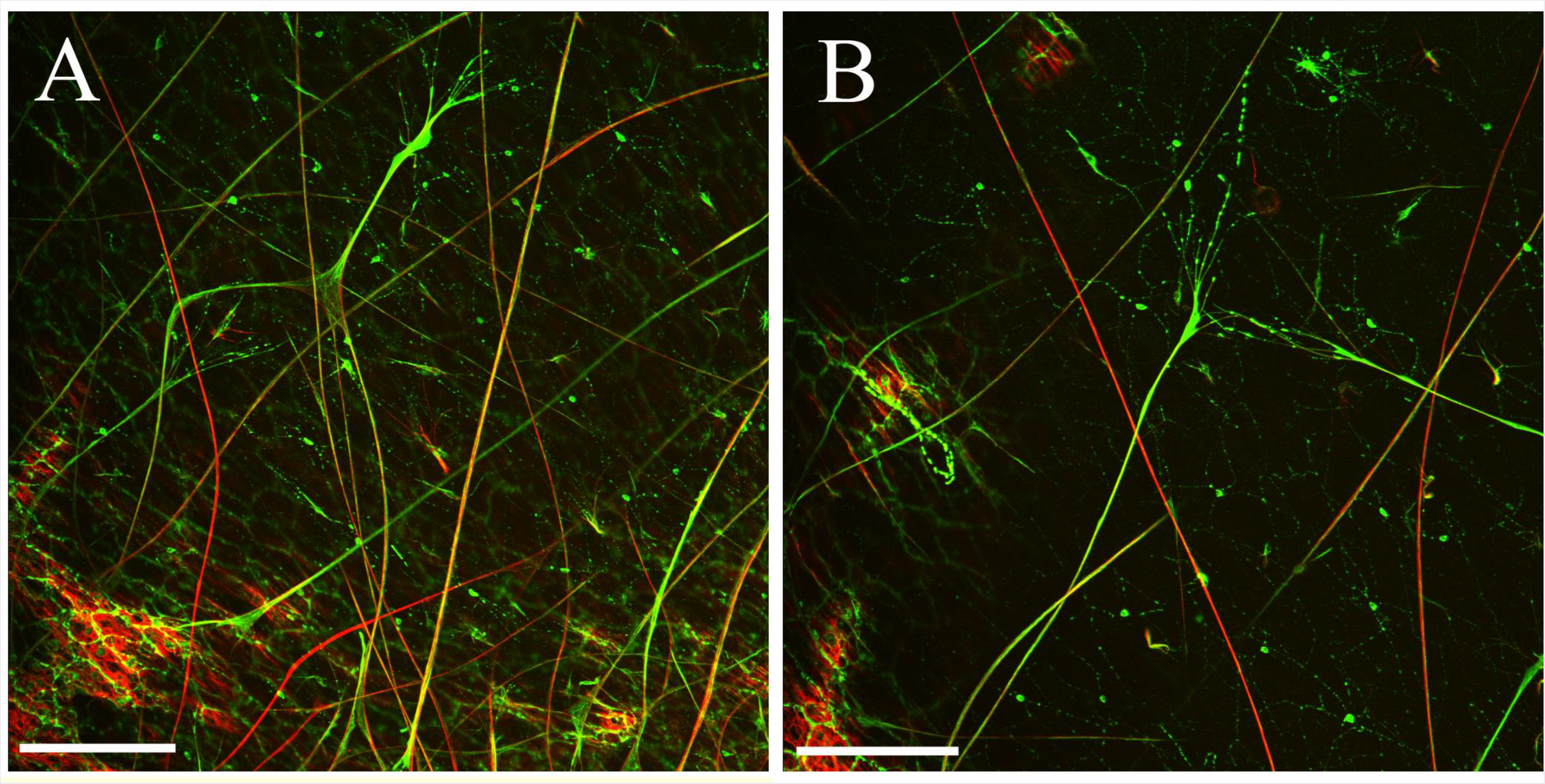
Co-localization of tubulin IR (green) and phalloidin (red) in selected mesoglea cells. **A, B**: Some of the tubulin-IR neuronal-like processes also demonstrate phalloidin staining. Scale bars: A, B - 100 μm.

## 4 DISCUSSION

Origins and early evolution of neural systems is a highly debated topic, and ctenophores are one of the key reference species (Striedter et al., 2014) in any reconstruction of the genealogy of cell types in Metazoa (Moroz, 2018). Recent advantages of comparative neuroscience and single-cell genomic point out to insufficient knowledge about the cellular diversity in basal metazoans in general, and ctenophores in particular. In fact, recent single-cell RNA-seq profiling of two ctenophores (Moroz, 2018; Sebe-Pedros et al., 2018) indicated that many cells could not be reliably recognized using conventional approaches. The absence of cell-specific molecular markers for ctenophore neurons and muscles is a noticeable bottleneck in the field. As a first step, a systematic reference microscopic atlas is needed for *Pleurobrachia* and related species. This work is based on earlier studies (Hernandez-Nicaise, 1991; Jager et al., 2011; Jager et al., 2013; Moroz et al., 2014; Norekian and Moroz, 2016) and provide the detailed atlas of the neuro-muscular organization of *Pleurobrachia bachei*.

Using two markers (tubulin IR and phalloidin), we conservatively recognized at least 45 distinct morphological types of cells related to sensory, neuronal and effector systems (Tables 1–4) as well as male and female gonads, colloblasts, pores in the canals, epithelial cells, balances, phalloidin stained cells in aboral organ, secretory cells, etc. Combined with electron microscopical observations (Horridge, 1965; Hernandez-Nicaise, 1968; 1973b; a; c; Hernandez-Nicaise, 1991), we estimate that at least 80 morphologically and ultrastructurally distinct cell types present in ctenophores, including at least 9 types of neurons and neuronal-like cells, 6 types of putative receptors and a large diversity of muscles and muscle-like elements – many of them described here for the first time.

The overall patterns of neuro-muscular systems in *Pleurobrachia bachei* is similar to those reported in a related species (Jager et al., 2011), but here, we recognized novel types of muscular, ciliated and neuronal-like cells as well as six distinct receptors (vs. one or two types described previously – see also (Hernandez-Nicaise, 1991)). These data have been complemented by scanning electron microscopy, which is challenging, if not impossible, on such fragile species as *Mnemiopsis* or *Bolinopsis*.

There is an unexpectedly complex organization of the tentacular nerves and innervation of the tentacle pockets, which might reflect the role of tentacles in prey detection and feeding including a stereotyped rotation behavior, characteristic for this species (Tamm, 1982).

Importantly, we identified connections between the subepithelial and mesogleal networks, suggesting their functional coupling (see also (Hernandez-Nicaise, 1968; Hernandez-Nicaise, 1991)), despite remarkable differences in 3D organization and cellular composition. The subepithelial network was never labeled by phalloidin, but some mesogleal neuron-like elements could be stained by both the tubulin antibody and phalloidin with at least three distinct morphological types. Furthermore, we detected connections between the subepithelial nerve net and the polar fields as well as ciliated conductive tracts – which provides strong morphological support for the integration of all parts of the neuro-muscular system.

Our mapping suggests that neurons in the subepithelial net can perform at least three canonical functions: (a) be motoneurons for muscles (Fig. 11I, with glutamate as a neuromuscular transmitter (Moroz et al., 2014)), (b) be a primarily sensory cells with at least two, or more modalities (e.g. mechanosensation with different strength, and a possible chemoreception (Kass-Simon and Hufnagel, 1992)), (c) be interneurons since we identified many cells with branched neurites as well as connections to the mesogleal net, polar fields, ciliated furrows, combs, tentacle nerves, and mouth. It is also possible that a simpler nerve net in ctenophores and their neurons might combine multiple functions (Hernandez-Nicaise, 1974); and be physiologically as well as chemically highly heterogeneous recruiting different neuropeptides and non-conventional low molecular weight transmitters (see discussion in (Moroz, 2015; Moroz and Kohn, 2015).

Control of comb beating occurs under three mechanisms (i) neuronal inputs; (ii) cilia-based coupling and conductance; and (iii) coordination, conduction, and coupling by non-contractive muscle cells including those cells between comb plates described in the current study (Figs. 27 B, C and 28 A, B). These inter-comb bridges can be an analog to the Purkinje fibers, the electrical conductive system in the mammalian heart when muscle derivatives ‘lost’ contractility and ‘become’ neuroid-like elements mediating synchronous conduction and coupling of the entire organ. The similar conjecture can be applied to enigmatic and quite diverse mesoglea cell populations with mixed patterns of phalloidin/tubulin IR.

Tubulin labeling and distinct morphology releal the presence of at least four types of neurons related to the subepithelial net with connections to combs, the aboral organ, tentacles and other parts of the feeding system such as pharynx. We also classified at least three types of tubulin IR mesogleal neurons, some of them shared morphological characteristics with phalloidin (only)-stained muscles. Mesogleal muscles might act as part of the body hydroskeleton and mediate retractions of combs or support functions of the pharynx or other organs. However, there are some muscle-like/neuron-like cells, not only with the shared morphology but also can be double labeled both with both tubulin AB and the F-actin marker - phalloidin. In other words, these cells have a ‘dual’ nature and, perhaps, shared origins.

Could neurons evolve from muscle-type ancestor lineages? An instant hypothesis: the mesogleal neurons and some other ‘neuroid’ populations are derived from muscle-like cells in ctenophores. Moreover, over the course of evolution, ‘true’ contractive muscle cells might be transformed into neuronal-like elements similar to fibers between combs or fibers in the mesoglea. Equally possible, that muscle and neuronal cells might have a common ancestral cell lineage in evolution and, therefore, share a core gene regulatory network. Considering that mesoderm and muscles in ctenophores can evolve independently from those in bilaterians (Moroz et al., 2014) - the described diversity of cell types in ctenophores can be a fertile ground for testing hypotheses of convergent evolution on the cellular level.

It is equally possible that neural-like elements might share the common ancestry with ciliated cells, which reach their utmost diversity of forms and functions in Ctenophora - more than in any other clade of the animal kingdom. We treat these evolutionary reconstructions, not as an alternative, but rather complementary hypotheses of the convergent evolution of cell phenotypes in Metazoa. Moreover, the hypothesis of convergent evolution of neurons and muscles in ctenophores is congruent with any outcome of the ongoing discussion about the Ctenophora-sister (Whelan et al., 2015b; Arcila et al., 2017; Shen et al., 2017; Whelan et al., 2017) vs. Porifera-sister (Pisani et al., 2015; Feuda et al., 2017; Simion et al., 2017) phylogenetic hypotheses. Indeed, many genes involved in the neuronal and muscular specification and functions had not been identified in the sequenced ctenophore genomes or, if such gene orthologs are present, they are not dependable neuronal or muscular markers (Moroz et al., 2014; Moroz, 2015; Moroz and Kohn, 2015; 2016).

Life of ctenophores is based on cilia (Tamm, 2014), and ctenophores took full advantage to explore cilia to achieve countless innovations in structures and functions including signal transmission as in the ciliated furrows. In summary, both ciliated cells and muscles have greater overall morphological diversity than neurons. However, we suspect that there is a hidden molecular diversity among all cell types and mostly in ctenophore neurons, which can be revealed in the ongoing and future systematic comparisons of cell-specific genomic, proteomic and metabolomic studies (Moroz, 2018). The presented atlas of the neuro-muscular organization in *Pleurobrachia* would be one of many reference platforms to understand the complex life of comb jellies better.

## Acknowledgments

We thank FHL for their excellent collection and microscope facilities including the new Nikon Laser Scanning confocal microscope; Dr. Victoria Foe for the use of the BioRad confocal microscope and Dr. Adam Summers for the use of SEM. We also thank Dr. Claudia Mills and Dr. Billie Swalla for useful ctenophore discussions.

This work was supported by the United States National Aeronautics and Space Administration (grant NASA-NNX13AJ31G), the National Science Foundation (grants 1146575, 1557923, 1548121 and 1645219) and National Institute of Health (grants R01GM097502, R01MH097062-01A1).

## Role of the authors

All authors had full access to all the data in the study and take responsibility for the integrity of the data and the accuracy of the data analysis. TPN and LLM share authorship equally. Research design: TPN, LLM. Acquisition of data: TPN, LLM. Analysis and interpretation of data: TPN, LLM. Drafting of the article: TPN, LLM. Funding: LLM.

## LITERATURE CITED

Alamaru A, Hoeksema BW, van der Meij SET, Huchon D. 2017. Molecular diversity of benthic ctenophores (Coeloplanidae). Sci Rep 7(1):6365.

Arcila D, Orti G, Vari R, Armbruster JW, Stiassny MLJ, Ko KD, Sabaj MH, Lundberg J, Revell LJ, Betancur RR. 2017. Genome-wide interrogation advances resolution of recalcitrant groups in the tree of life. Nat Ecol Evol 1(2):20.

Aronova M. 1974. Electron microscopic observation of the aboral organ of Ctenophora. I. The gravity receptor. Z Mikrosk Anat Forsch 88(2):401–412.

Borowiec ML, Lee EK, Chiu JC, Plachetzki DC. 2015. Extracting phylogenetic signal and accounting for bias in whole-genome data sets supports the Ctenophora as sister to remaining Metazoa. BMC Genomics 16:987.

Brusca RC, Brusca GJ. 2003. Invertebrates. Sunderland, Massachusetts: Sinauer Associates, Inc. 936 p.

Cavalier-Smith T. 2017. Origin of animal multicellularity: precursors, causes, consequences-the choanoflagellate/sponge transition, neurogenesis and the Cambrian explosion. Philos Trans R Soc Lond B Biol Sci 372(1713).

Dunn CW, Leys SP, Haddock SH. 2015. The hidden biology of sponges and ctenophores. Trends Ecol Evol 30(5):282–291.

Feuda R, Dohrmann M, Pett W, Philippe H, Rota-Stabelli O, Lartillot N, Worheide G, Pisani D. 2017. Improved Modeling of Compositional Heterogeneity Supports Sponges as Sister to All Other Animals. Curr Biol 27(24):3864-3870 e3864.

Halanych KM, Whelan NV, Kocot KM, Kohn AB, Moroz LL. 2016. Miscues misplace sponges. Proc Natl Acad Sci U S A 113(8):E946–947.

Hernandez-Nicaise M-L. 1991. Ctenophora. In: F.W. Harrison FW, Westfall JA, eds. Microscopic Anatomy of Invertebrates: Placozoa, Porifera, Cnidaria, and Ctenophora. Vol 2. New York: Wiley. p 359418.

Hernandez-Nicaise ML. 1968. Specialized connexions between nerve cells and mesenchymal cells in ctenophores. Nature 217(5133):1075–1076.

Hernandez-Nicaise ML. 1973a. [The nervous system of ctenophores. I. Structure and ultrastructure of the epithelial nerve-nets]. Z Zellforsch Mikrosk Anat 137(2):223–250.

Hernandez-Nicaise ML. 1973b. [The nervous system of ctenophores. II. The nervous elements of the mesoglea of beroids and cydippids (author’s transl)]. Z Zellforsch Mikrosk Anat 143(1):117–133.

Hernandez-Nicaise ML. 1973c. The nervous system of ctenophores. III. Ultrastructure of synapses. J Neurocytol 2(3):249–263.

Hernandez-Nicaise ML. 1974. Ultrastructural evidence for a sensory-motor neuron in Ctenophora. Tissue Cell 6(1):43–47.

Horridge GA. 1965. Relations between Nerves and Cilia in Ctenophores. Am Zool 5:357–375.

Jager M, Chiori R, Alie A, Dayraud C, Queinnec E, Manuel M. 2011. New insights on ctenophore neural anatomy: immunofluorescence study in *Pleurobrachia pileus* (Muller, 1776). J Exp Zool B Mol Dev Evol 316B(3):171–187.

Jager M, Dayraud C, Mialot A, Queinnec E, le Guyader H, Manuel M. 2013. Evidence for involvement of Wnt signalling in body polarities, cell proliferation, and the neuro-sensory system in an adult ctenophore. PLoS One 8(12):e84363.

Kass-Simon G, Hufnagel LA. 1992. Suspected chemoreceptors in coelenterates and ctenophores. Microsc Res Tech 22(3):265–284.

King N, Rokas A. 2017. Embracing Uncertainty in Reconstructing Early Animal Evolution. Curr Biol 27(19):R1081–R1088.

Kozloff EN. 1990. Invertebrates. Philadelphia: Sounders College Publishing. 866 p.

Lowe B. 1997. The role of Ca2+ in deflection-induced excitation of motile, mechanoresponsive balancer cilia in the ctenophore statocyst. J Exp Biol 200(Pt 11):1593–1606.

Moroz LL. 2014. The genealogy of genealogy of neurons. Commun Integr Biol 7(6):e993269.

Moroz LL. 2015. Convergent evolution of neural systems in ctenophores. J Exp Biol 218(Pt 4):598–611.

Moroz LL. 2018. NeuroSystematics and Periodic System of Neurons: Model vs Reference Species at Single-cell Resolution. ACS Chem Neurosci.

Moroz LL, Kocot KM, Citarella MR, Dosung S, Norekian TP, Povolotskaya IS, Grigorenko AP, Dailey C, Berezikov E, Buckley KM, Ptitsyn A, Reshetov D, Mukherjee K, Moroz TP, Bobkova Y, Yu F, Kapitonov VV, Jurka J, Bobkov YV, Swore JJ, Girardo DO, Fodor A, Gusev F, Sanford R, Bruders R, Kittler E, Mills CE, Rast JP, Derelle R, Solovyev VV, Kondrashov FA, Swalla B J, Sweedler JV, Rogaev EI, Halanych KM, Kohn AB. 2014. The ctenophore genome and the evolutionary origins of neural systems. Nature 510(7503):109–114.

Moroz LL, Kohn AB. 2015. Unbiased View of Synaptic and Neuronal Gene Complement in Ctenophores: Are There Pan-neuronal and Pan-synaptic Genes across Metazoa? Integr Comp Biol 55(6):1028–1049.

Moroz LL, Kohn AB. 2016. Independent origins of neurons and synapses: insights from ctenophores. Philos Trans R Soc Lond B Biol Sci 371(1685):20150041.

Nielsen C. 2012. Animal Evolution: Interrelationships of the living phyla. Oxford: Oxford University Press.

Norekian TP, Moroz LL. 2016. Development of neuromuscular organization in the ctenophore Pleurobrachia bachei. J Comp Neurol 524(1):136–151.

Pisani D, Pett W, Dohrmann M, Feuda R, Rota-Stabelli O, Philippe H, Lartillot N, Worheide G. 2015. Genomic data do not support comb jellies as the sister group to all other animals. Proc Natl Acad Sci U S A 112(50):15402–15407.

Presnell JS, Vandepas LE, Warren KJ, Swalla BJ, Amemiya CT, Browne WE. 2016. The Presence of a Functionally Tripartite Through-Gut in Ctenophora Has Implications for Metazoan Character Trait Evolution. Curr Biol 26(20):2814–2820.

Ryan JF, Pang K, Schnitzler CE, Nguyen AD, Moreland RT, Simmons DK, Koch BJ, Francis WR, Havlak P, Program NCS, Smith SA, Putnam NH, Haddock SH, Dunn CW, Wolfsberg TG, Mullikin JC, Martindale MQ, Baxevanis AD. 2013. The genome of the ctenophore Mnemiopsis leidyi and its implications for cell type evolution. Science 342(6164):1242592.

Sebe-Pedros A, Chomsky E, Pang K, Lara-Astiaso D, Gaiti F, Mukamel Z, Amit I, Hejnol A, Degnan BM, Tanay A. 2018. Early metazoan cell type diversity and the evolution of multicellular gene regulation. Nat Ecol Evol 2(7):1176–1188.

Shen XX, Hittinger CT, Rokas A. 2017. Contentious relationships in phylogenomic studies can be driven by a handful of genes. Nat Ecol Evol 1(5):126.

Simion P, Philippe H, Baurain D, Jager M, Richter DJ, Di Franco A, Roure B, Satoh N, Queinnec E, Ereskovsky A, Lapebie P, Corre E, Delsuc F, King N, Worheide G, Manuel M. 2017. A Large and Consistent Phylogenomic Dataset Supports Sponges as the Sister Group to All Other Animals. Curr Biol 27(7):958–967.

Steinmetz PR, Kraus JE, Larroux C, Hammel JU, Amon-Hassenzahl A, Houliston E, Worheide G, Nickel M, Degnan BM, Technau U. 2012. Independent evolution of striated muscles in cnidarians and bilaterians. Nature 487(7406):231–234.

Striedter GF, Belgard TG, Chen CC, Davis FP, Finlay BL, Gunturkun O, Hale ME, Harris JA, Hecht EE, Hof PR, Hofmann HA, Holland LZ, Iwaniuk AN, Jarvis ED, Karten HJ, Katz PS, Kristan WB, Macagno ER, Mitra PP, Moroz LL, Preuss TM, Ragsdale CW, Sherwood CC, Stevens CF, Stuttgen MC, Tsumoto T, Wilczynski W. 2014. NSF workshop report: discovering general principles of nervous system organization by comparing brain maps across species. J Comp Neurol 522(7):1445–1453.

Tamm SL. 1982. Ctenophora. Electrical conduction and behavior in “simple” invertebrates. Oxford: Clarendon Press. p 266–358.

Tamm SL. 2014. Cilia and the life of ctenophores. Invertebrate Biology 133(1):1–46.

Tamm SL. 2015. Functional consequences of the asymmetric architecture of the ctenophore statocyst. Biol Bull 229(2):173–184.

Tamm SL. 2016a. No surprise that comb jellies poop. Science 352(6290):1182.

Tamm SL. 2016b. Novel Structures Associated with Presumed Photoreceptors in the Aboral Sense Organ of Ctenophores. Biol Bull 231(2):97–102.

Tamm SL, Tamm S. 2002. Novel bridge of axon-like processes of epithelial cells in the aboral sense organ of ctenophores. J Morphol 254(2):99–120.

Telford MJ, Moroz LL, Halanych KM. 2016. Evolution: A sisterly dispute. Nature 529(7586):286–287.

Wehland J, Willingham MC. 1983. A rat monoclonal antibody reacting specifically with the tyrosylated form of alpha-tubulin. II. Effects on cell movement, organization of microtubules, and intermediate filaments, and arrangement of Golgi elements. J Cell Biol 97(5 Pt 1):1476–1490.

Wehland J, Willingham MC, Sandoval IV. 1983. A rat monoclonal antibody reacting specifically with the tyrosylated form of alpha-tubulin. I. Biochemical characterization, effects on microtubule polymerization in vitro, and microtubule polymerization and organization in vivo. J Cell Biol 97(5 Pt 1):1467–1475.

Whelan NV, Kocot KM, Halanych KM. 2015a. Employing Phylogenomics to Resolve the Relationships among Cnidarians, Ctenophores, Sponges, Placozoans, and Bilaterians. Integr Comp Biol 55(6):1084–1095.

Whelan NV, Kocot KM, Moroz LL, Halanych KM. 2015b. Error, signal, and the placement of Ctenophora sister to all other animals. Proc Natl Acad Sci U S A 112(18):5773–5778.

Whelan NV, Kocot KM, Moroz TP, Mukherjee K, Williams P, Paulay G, Moroz LL, Halanych KM. 2017. Ctenophore relationships and their placement as the sister group to all other animals. Nat Ecol Evol 1(11):1737–1746.

Wulf E, Deboben A, Bautz FA, Faulstich H, Wieland T. 1979. Fluorescent phallotoxin, a tool for the visualization of cellular actin. Proc Natl Acad Sci U S A 76(9):4498–4502.

